# Supramolecular hydrogel viscoelasticity regulates in situ tertiary lymphoid neogenesis

**DOI:** 10.64898/2026.07.20.739456

**Authors:** Ryan Ramsey Hosn, Oriana Marrone Mantovani, Artemis Margaronis, James S. Wang, Ethan Kriss, Claire Wu, Michael Kissner, José L. McFaline-Figueroa, Santiago Correa

**Affiliations:** Department of Biomedical Engineering, Columbia University, New York, NY 10027, USA; Columbia Stem Cell Initiative, Columbia University, New York, NY 10032, USA; Herbert Irving Comprehensive Cancer Center, Columbia University, New York, NY 10032, USA; Irving Institute for Cancer Dynamics, Columbia University, New York, NY 10027, USA

## Abstract

Under chronic inflammation, the immune system can spontaneously assemble lymphoid tissues called tertiary lymphoid structures (TLSs). When TLSs arise in tumors, they are associated with improved response to immunotherapy, galvanizing interest in inducing them with biomaterials. These efforts have focused primarily on sustained release of TLS-inducing cues, leaving the influence of the material’s mechanical properties on TLS neogenesis unexamined. Here we varied the viscoelastic properties of an injectable liposomal hydrogel while delivering CCL21 and LIGHT at comparable rates. We found that soft hydrogels were rapidly infiltrated and remodeled, progressing from a myeloid-rich infiltrate to vascularized, B-cell-rich TLS-like tissue, with the lymphoid-to-myeloid ratio rising tenfold between days 7 and 14. In contrast, stiff hydrogels resisted remodeling, showed a fourfold longer in vivo half-life, and sustained a myeloid-dominant niche. In soft hydrogels, single-cell transcriptomics identified a transient interferon-associated myeloid response preceding this organization. Finally, soft but not stiff hydrogels reduced early tumor burden threefold in a prophylactic cancer vaccine setting. These results identify viscoelasticity as a key parameter in biomaterials-mediated lymphoid tissue engineering.

## Introduction

Tertiary lymphoid structures (TLSs) are ectopic immune tissues that form within tumors and have emerged as a critical biomarker in cancer immunotherapy^1,2^, associated with significantly improved survival and favorable response to checkpoint blockade^3–6^. Those associations have driven efforts to induce TLSs deliberately^7–9^ and spurred interest in engineered materials as a translationally relevant approach to TLS induction^10–12^. These efforts are complicated by the heterogeneity of TLSs themselves, which range from loose, immature aggregates to organized structures that mirror lymph node architecture, with adjacent B- and T-cell zones, antigen-presenting and stromal networks, and specialized vasculature that sustains immune-cell recruitment^13–15^. TLS maturation, particularly the formation of germinal centers, is associated with stronger prognostic benefit^16–18^. Reproducing that architecture requires cellular recruitment, stromal activation, vascular remodeling and lymphocyte organization to proceed together^19^.

Biomaterials have shown early promise as a route to TLS induction^10,20^, but the mechanical properties of the material itself have received surprisingly little attention. Implanted and injectable materials that release lymphoid-organizing cues have generated TLS-like tissues with antitumor activity in ectopic and intratumoral settings^21–27^, and these efforts have focused predominantly on sustained release of TLS-inducing biochemical cues, leaving unclear which material properties govern success or failure. Across many cell types, matrix stiffness, viscoelasticity and stress relaxation shape how cells adhere, spread, migrate and differentiate^28–30^. They also shape the structures those cells build^31,32^. These relationships are far better defined in stromal^31,33,34^ and progenitor^28,29,32^ systems than in immune cells^31,35–38^, and have not been explored for TLS neogenesis.

We hypothesized that the mechanical properties of a material would shape the immune tissue that forms within it, independent of the biochemical cues it delivers. To test this, we used an injectable liposome-crosslinked supramolecular hydrogel^39^ in which network mechanics and cargo release are controlled by separate molecular handles. Network viscoelasticity is set by the alkyl-chain length of the hydrophobically modified polymer, whereas retention of the protein cargo is governed by the liposomal crosslinkers. Changing alkyl-chain length alone therefore yielded formulations that differed in their viscoelastic properties, in how readily host cells can deform and replace them, and in how long they persist in vivo, while releasing the same cues at comparable rates. Each formulation carried CCL21 to recruit the lymphocytes and antigen-presenting cells that build TLSs^10,22,40^, LIGHT to engage the lymphotoxin-β receptor signaling that drives TLS organization^21,41,42^, and ovalbumin (OVA) as a defined model antigen.

Despite receiving identical biochemical cues, soft and stiff hydrogels produced markedly different immune niches. We established TLS-inducing niches in the subcutaneous space at defined times and followed their development longitudinally, combining histology, immunofluorescence imaging, flow cytometry and single-cell RNA sequencing. Soft hydrogels were infiltrated and progressively replaced, assembling vascularized, B-cell-rich tissue that carried TLS-associated transcriptional programs, whereas stiff hydrogels relegated cells to their periphery and remained myeloid-dominant, with little lymphoid organization. Single-cell profiling identified a transient interferon-associated myeloid response that preceded lymphoid organization in soft hydrogels. In an antigen-matched melanoma model, a single prophylactic implantation of the soft formulation improved early tumor control, whereas stiff hydrogels performed comparably to empty controls. These results identify viscoelasticity as a key parameter shaping the immune niche that forms within a biomaterial, and one that may be especially consequential for efforts to induce TLSs.

## Results and Discussion

### 1. Hydrogel mechanics set persistence and spatial remodeling

To investigate how hydrogel mechanics influence the development of biomaterial-induced immune niches, we used injectable supramolecular hydrogels in which hydrophobically modified hydroxypropyl methylcellulose (HPMC) is physically crosslinked by anionic liposomes^39^ (**Supplementary Fig. 1**). HPMC modified with 12-carbon (C12) or 18-carbon (C18) alkyl chains formed hydrogels with distinct mechanical properties. Because alkyl-chain length tunes the polymer-liposome network while liposome composition and cargo loading are held constant, this platform allowed network mechanics to be varied under matched initial biochemical conditions. At 10 rad/s, the HPMC-C12 and HPMC-C18 formulations exhibited storage moduli of 58.9 ± 10.0 Pa and 246.8 ± 25.9 Pa, respectively (p < 0.0001; **Fig. 1A**–**C**). For simplicity, we refer to HPMC-C12 as the soft hydrogel and HPMC-C18 as the stiff hydrogel; subsequent measurements further defined their distinct viscoelastic behavior. The stiff formulation showed a lower tanδ and weaker frequency dependence, consistent with a more persistent elastic network (**Supplementary Table 1**). The networks diverged further under large and sustained deformation. The stiff hydrogel yielded at 437 ± 17.5 % strain compared with 90.8 ± 9.4 % for the soft hydrogel (p = 0.0004; **Fig. 1D**–**E**), while creep under 12.5 Pa stress was markedly greater in the soft network (13.6 versus 0.084 % s^-1^; **Supplementary Fig. 2**). Both formulations remained shear-thinning and recovered after deformation, supporting minimally invasive delivery (**Fig. 1F**; **Supplementary Fig. 2**). More detailed linear and nonlinear rheology is provided in the Supplementary Information.

**Figure 1.**
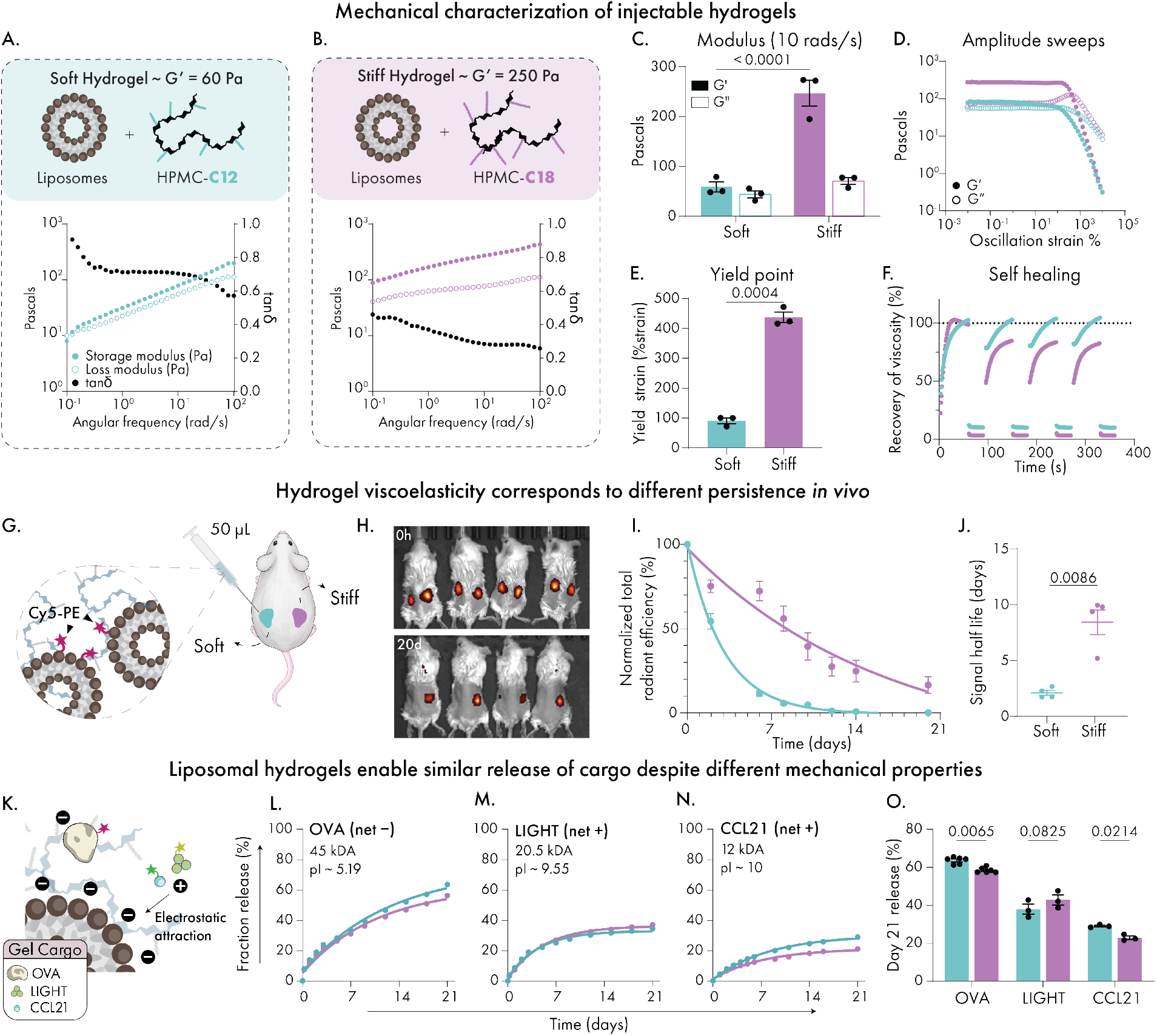
Mechanical tuning of supramolecular hydrogels regulates *in vivo* persistence while maintaining broadly comparable protein release. (**A**–**B**) Schematic representation and frequency-dependent mechanical characterization of supramolecular hydrogels formed by combining liposomes with hydrophobically modified HPMC polymers bearing C12 (soft) or C18 (stiff) alkyl chains. Representative storage (G′), loss (G″) moduli, and tan δ profiles demonstrate solid-like behavior for both formulations across the measured frequency range. (**C**) Ǫuantification of storage and loss moduli at 10 rad s⁻¹. (**D**) Representative oscillatory amplitude sweeps of soft and stiff hydrogels. (**E**) Yield strain quantification demonstrating enhanced resistance to deformation in stiff hydrogels. Rheological quantification in (C) and (E) was performed using n = 3 independently prepared hydrogel batches per formulation. (**F**) Step-strain self-healing assays showing viscosity recovery following repeated shear. (**G**) Schematic of subcutaneous injection of Cy5-labeled empty hydrogels for longitudinal in vivo persistence studies. In vivo persistence analysis included n = 4 biologically independent mice, each receiving one soft hydrogel on the left flank and one stiff hydrogel on the right flank. (**H**) Representative IVIS images collected at 0 h and day 20. (**I**) Ǫuantification of normalized radiant efficiency over time. (**J**) Signal half-life quantification comparing soft and stiff hydrogels. (**K**) Schematic illustrating charge-dependent retention of protein cargos within the liposome-crosslinked hydrogel network. (**L**–**N**) In vitro cumulative release profiles of ovalbumin (OVA), LIGHT and CCL21 from soft and stiff hydrogels over 21 days (n = 3). (**O**) Cumulative release fractions at day 21 for OVA, LIGHT and CCL21 from soft and stiff hydrogels. OVA data represent n = 6 measurements, whereas LIGHT and CCL21 each represent n = 3. Data are represented as mean ± SEM. Statistical significance in (C) and (O) was assessed using two-way ANOVA followed by Šídák’s multiple-comparisons test. Differences in yield strain (E) and signal half-life (J) were assessed using unpaired two-tailed Student’s t-test. Exact *p* values are shown where indicated.

These differences in hydrogel viscoelasticity translated into distinct persistence *in vivo*. Empty hydrogels containing Cy5-labeled liposomes were implanted subcutaneously and followed longitudinally by IVIS (**Fig. 1G**–**J**; **Supplementary Fig. 3A**–**C**). Material-associated signal from soft hydrogels decayed with a half-life of 2.1 ± 0.2 days and was undetectable by approximately two weeks. Stiff hydrogels persisted with a half-life of 8.4 ± 1.1 days and retained 24.6 ± 6.4% of their initial signal at day 14 and 16.3 ± 5.0% by week three. At endpoint, empty soft hydrogels were no longer macroscopically identifiable, whereas residual material from stiff hydrogels remained (**Supplementary Fig. 3D**). These differences are consistent with the greater resistance to creep of the stiff network and with prior work linking reduced creep to longer subcutaneous depot persistence^43^.

Despite their distinct mechanical properties, soft and stiff hydrogels exhibited broadly similar release kinetics. Anionic liposomal crosslinkers were designed to promote retention of the positively charged LIGHT and CCL21 through dynamic electrostatic interactions (**Fig. 1K**; **Supplementary Table 2**). Accordingly, over 21 days, both proteins reached cumulative release plateaus below approximately 40%, whereas OVA approached 60%, with comparable release across soft and stiff formulations (**Fig. 1L**–**O**). At day 21, cumulative release from soft and stiff hydrogels was 62.5% and 57.5% for OVA, 37.9% and 42.8% for LIGHT, and 29.1% and 22.9% for CCL21, respectively. Although differences in OVA and CCL21 were statistically significant, their magnitude was modest, and the overall release trajectories remained broadly similar between formulations. Thus, the formulations exhibited broadly comparable intrinsic release behavior, although their divergent *in vivo* persistence could subsequently alter local cue availability as the materials are remodeled.

We next examined how these differences shaped remodeling *in vivo*. Cy5-labeled soft and stiff hydrogels containing OVA (35 μg), CCL21 (0.4 μg) and LIGHT (0.4 μg) per 50 μL gel (OCL) were collected at days 7, 14, and 21 and analyzed with matched histological sections (**Fig. 2A**–**B**; **Supplementary Fig. 4**). At day 7, residual hydrogel occupied a similar fraction of soft and stiff explants, but its spatial relationship with host tissue already differed. Soft explants contained cell- and collagen-rich tissue throughout the hydrogel, whereas denser cellular and collagen-rich remodeling in stiff explants was concentrated toward the host-material interface. Residual hydrogel in soft explants decreased to 13.4 ± 7.1% of cross-sectional area by day 14 and was undetectable by day 21, while stiff explants retained 44 ± 14.2% and 47.3 ± 7.2% at the same timepoints (**Fig. 2C**). Soft explant mass declined in parallel, while nuclear density within histologically retained tissue increased from 392 ± 50 nuclei mm^-2^ to 691 ± 37 nuclei mm^-2^ between days 7 and 21, consistent with progressive replacement of the hydrogel by host tissue (**Fig. 2D**–**E**). Representative higher magnification day-21 HCE and Masson’s trichrome views further illustrate regions of dense tissue remodeling in soft explants and within the peripheral remodeled zone of stiff explants (**Fig. 2F**). Despite the peripheral concentration of dense remodeled tissue in stiff hydrogels, DAPI⁺ nuclei staining on preserved gel core sections revealed cellular access to the persistent Cy5^+^ scaffold region (**Supplementary Fig. 5**). The inclusion of OCL cargo in soft hydrogels led to the formation of a persistent ectopic tissue that remained explantable at later timepoints, whereas empty soft hydrogels were no longer macroscopically identifiable.

**Figure 2.**
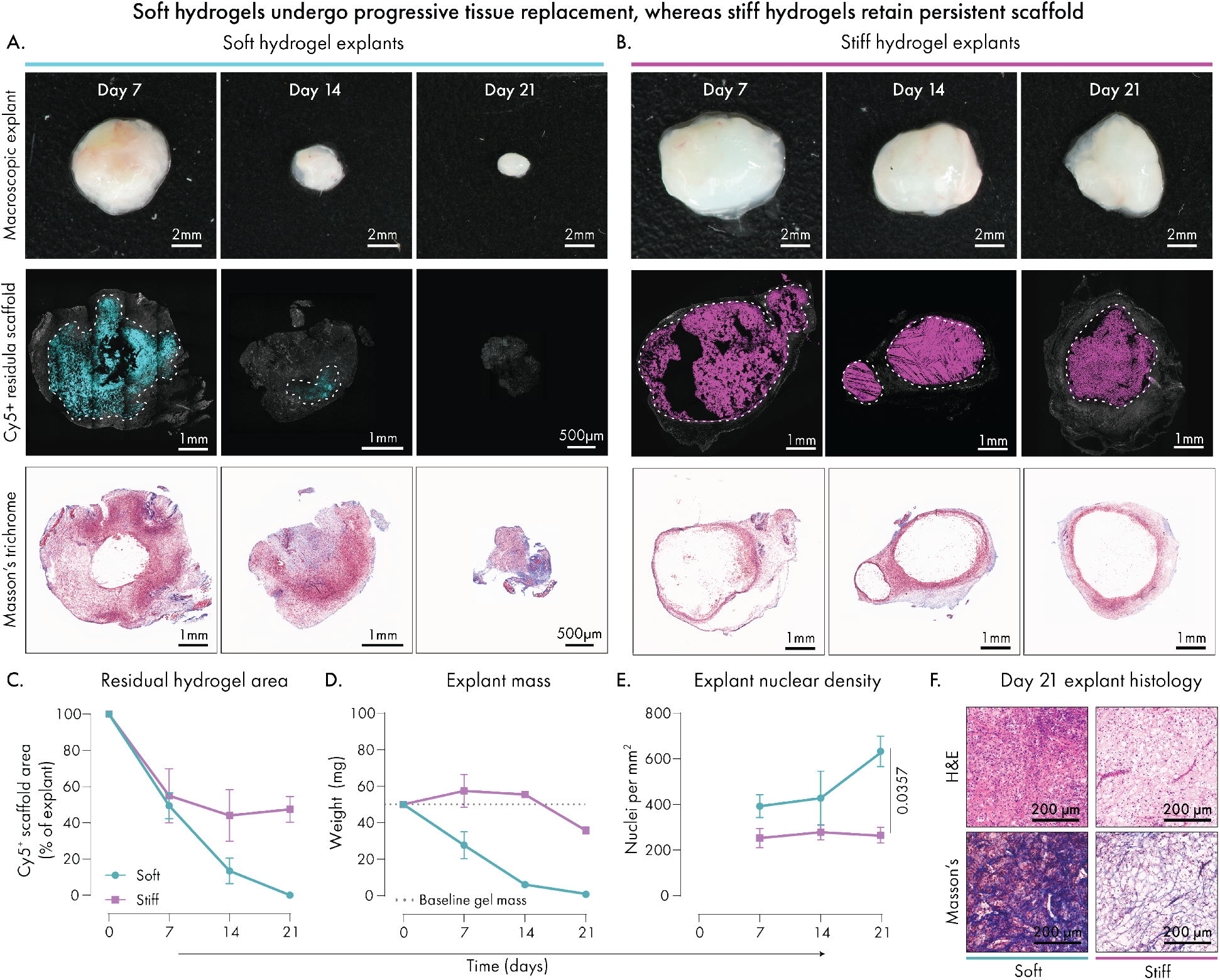
Hydrogel mechanical properties regulate spatial remodeling, material persistence, and cellular infiltration in vivo. (**A**–**B**) Representative macroscopic explant images, Cy5-labeled residual hydrogel sections, and corresponding Masson’s Trichrome-stained serial sections from soft (C12; A) and stiff (C18; B) hydrogels retrieved at days 7, 14, and 21 following subcutaneous injection. Hydrogels contained TLS-associated cytokines CCL21 and LIGHT together with model antigen ovalbumin. Cy5 signal delineates the residual hydrogel within the explant, with dotted outlines indicating the Cy5⁺ scaffold region. (**C**) Ǫuantification of residual hydrogel area, expressed as Cy5⁺ fraction of total explant cross-section over time. Day 0 was defined as 100% residual hydrogel for normalization and represents the theoretical pre-remodeling state rather than an experimentally sectioned sample. (**D**) Explant mass over 21 days following injection. The dotted line indicates the nominal initial hydrogel mass based on the injected 50 μL volume. (**E**) Ǫuantification of nuclear density within explant cross-section (nuclei per mm²) at each timepoint, derived from HCE-stained sections. (**F**) Representative day-21 HCE and Masson’s Trichrome images from soft and stiff explants illustrating cellular and matrix organization. (C–E) data are presented as mean ± SEM from n = 3 independent biological explants per formulation and timepoint, except for nuclear density in soft hydrogels at day 21 (E), where n = 5. Statistical comparisons of nuclear density at day 21 were performed using a Mann–Whitney U test. Exact *p* values are shown where indicated.

These observations establish two distinct remodeling regimes from otherwise matched formulations. The soft network behaves as a sacrificial template that supports distributed remodeling and is progressively replaced by host tissue, whereas the stiff network remains accessible to infiltrating cells but retains a persistent material, with dense tissue remodeling preferentially concentrated toward its periphery. Together with the rheological measurements, these in vivo remodeling patterns are consistent with a more compliant, deformation-permissive soft network and a more deformation-resistant stiff network. This divergence in material persistence and spatial remodeling provides the physical context for the distinct immune niches that subsequently emerge.

### 2. Hydrogel mechanics direct divergent immune niche architecture in vivo

With these distinct remodeling regimes established, we next asked whether matched OCL cargo would give rise to different immune tissues. By day 21, soft hydrogels contained B220⁺ B cell-rich clusters positioned adjacent to CD3⁺ T-cell regions and interspersed with networks of CD11c⁺ antigen-presenting cells (**Fig. 3A**–**B**). Stiff hydrogels instead showed diffuse, poorly compartmentalized lymphocyte aggregates (**Fig. 3C**–**D**). Thus, the soft niche displayed a compartmentalized architecture characteristic of TLS-like structures, whereas the persistent stiff material supported a less organized lymphoid architecture consistent with an immature TLS-like state^17,44^.

**Figure 3.**
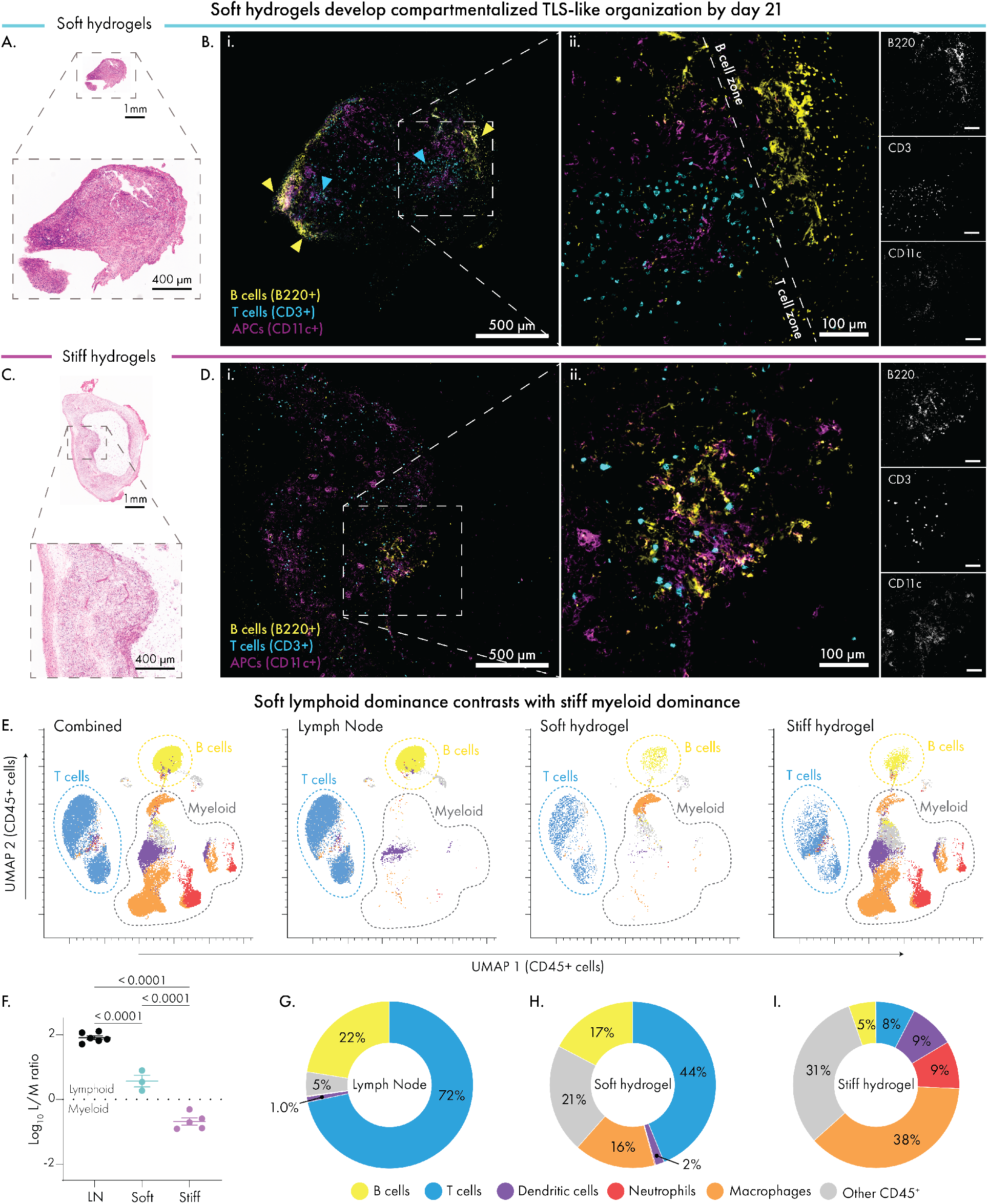
Hydrogel mechanics regulate immune composition and spatial organization at day 21 in vivo. Soft (C12) and stiff (C18) 50 μL hydrogels containing ovalbumin (OVA; 35 μg), CCL21 (0.4 μg), and LIGHT (0.4 μg) were implanted subcutaneously in BALB/c mice and analyzed after 21 days. Inguinal lymph nodes were collected as a reference lymphoid tissue. (**A**, **C**) Representative HCE-stained cross-sections of soft (A) and stiff (C) hydrogel explants, shown as whole-explant overviews and 4× enlarged regions. Explants were sectioned through their central plane for spatial characterization. (**B**, **D**) Immunofluorescence imaging of matched serial sections from soft (B) and stiff (D) hydrogel explants stained for B cells (B220⁺, yellow), T cells (CD3⁺, cyan), and CD11c⁺ antigen-presenting cells (magenta). Whole-explant overviews (i) and higher-magnification views of boxed regions (ii) are shown with corresponding single-channel images at right. Soft hydrogels contained spatially distinct B-cell- and T-cell-rich regions, indicated by yellow and cyan arrowheads, respectively. In contrast, stiff hydrogels showed diffuse intermixing of B cells, T cells, and CD11c⁺ cells without clear zonal segregation. (**E**) Unsupervised UMAP embeddings of flow-cytometry-profiled CD45⁺ cells shown for the combined dataset and separately for lymph nodes, soft hydrogels, and stiff hydrogels. Cells are colored according to annotated immune population. (**F**) Log₁₀-transformed lymphoid-to-myeloid cell ratio calculated from raw immune-cell counts. Lymphoid cells were defined as B and T cells, whereas myeloid cells comprised macrophages, neutrophils, and dendritic cells. Values above and below zero indicate lymphoid- and myeloid-dominant compositions, respectively. Raw immune-cell counts contributing to this ratio are shown in Supplementary Fig. 6. Data are presented as mean ± SEM. (**G–I**) Mean immune composition of lymph nodes (G), soft hydrogels (H), and stiff hydrogels (I), expressed as the percentage of CD45⁺ cells. Replicate-level quantification with mean ± SEM is shown in Supplementary Fig. 6. For flow-cytometry analyses (E–I), replicates consisted of n = 3 biologically independent soft hydrogel explants, n = 5 biologically independent stiff hydrogel explants, and n = 6 lymph nodes collected from individual mice. Statistical significance in (F) was assessed using one-way ANOVA followed by Tukey’s multiple-comparisons test; exact *p* values are shown.

Flow cytometry showed parallel compositional divergence. UMAP^45^ visualization of CD45⁺ cells identified lymphoid populations in both materials, but macrophages and neutrophil populations were more prominent in stiff hydrogels (**Fig. 3E**). The log_10_ lymphoid-to-myeloid ratio was positive in soft hydrogels (0.57 ± 0.18) but negative in stiff hydrogels (-0.68 ± 0.11) compared with 1.91 ± 0.06 in lymph nodes (**Fig. 3F**). Soft hydrogels contained 44 ± 5% T cells and 17 ± 2% B cells among CD45⁺ cells, approaching the lymphoid-rich composition of reference lymph nodes, whereas stiff hydrogels contained 8 ± 2% and 5 ± 1% respectively (**Fig. 3G**–**I**). Conversely, macrophages comprised 38 ± 5% of leukocytes in stiff hydrogels versus 16 ± 5% in soft hydrogels, while neutrophils accounted for 9.3 ± 3.0% of CD45⁺ cells in stiff hydrogels but were nearly absent from soft hydrogels (< 0.1%).

The T-cell compartment reinforced this distinction. Soft hydrogels exhibited a CD4-dominant ratio similar to lymph nodes and contained a greater fraction of naïve T cells, whereas stiff hydrogels showed a lower CD4:CD8 ratio and a greater effector-memory representation (**Supplementary Fig. 6**). Given the short circulating half-life of neutrophils, their continued presence at day 21, together with macrophage enrichment and a more activation-experienced T-cell state, is consistent with sustained inflammatory engagement and ongoing innate recruitment within the persistent stiff hydrogel^46,47^. In contrast, soft hydrogels more closely recapitulated a lymphoid-tissue-like immune composition^48^.

Because compartmentalized TLS-like organization was observed only in soft hydrogels within the 21-day window, we next focused on its temporal evolution across days 7, 14 and 21 (**Fig. 4**; **Supplementary Fig. 7**). At day 7, explants already contained small, loosely organized immune-cell aggregates, predominantly composed of mixed B and T cells, with less frequent T-cell-dominant and occasional CD11c⁺-rich aggregates (**Fig. 4A**; **Supplementary Figs. 7**, **8**). By day 14, organization was most pronounced with large B cell-rich aggregates positioned next to more diffuse T-cell regions, with CD11c⁺ cells within and around the B-cell zones (**Fig. 4B**). At day 21, organized lymphoid regions persisted, but B-cell-rich zones appeared smaller and less compact while T-cell zones remained prominent across the explant (**Fig. 4C**). This contraction of B cell-rich organization may reflect early stages of involution of the TLS-like niche.

**Figure 4.**
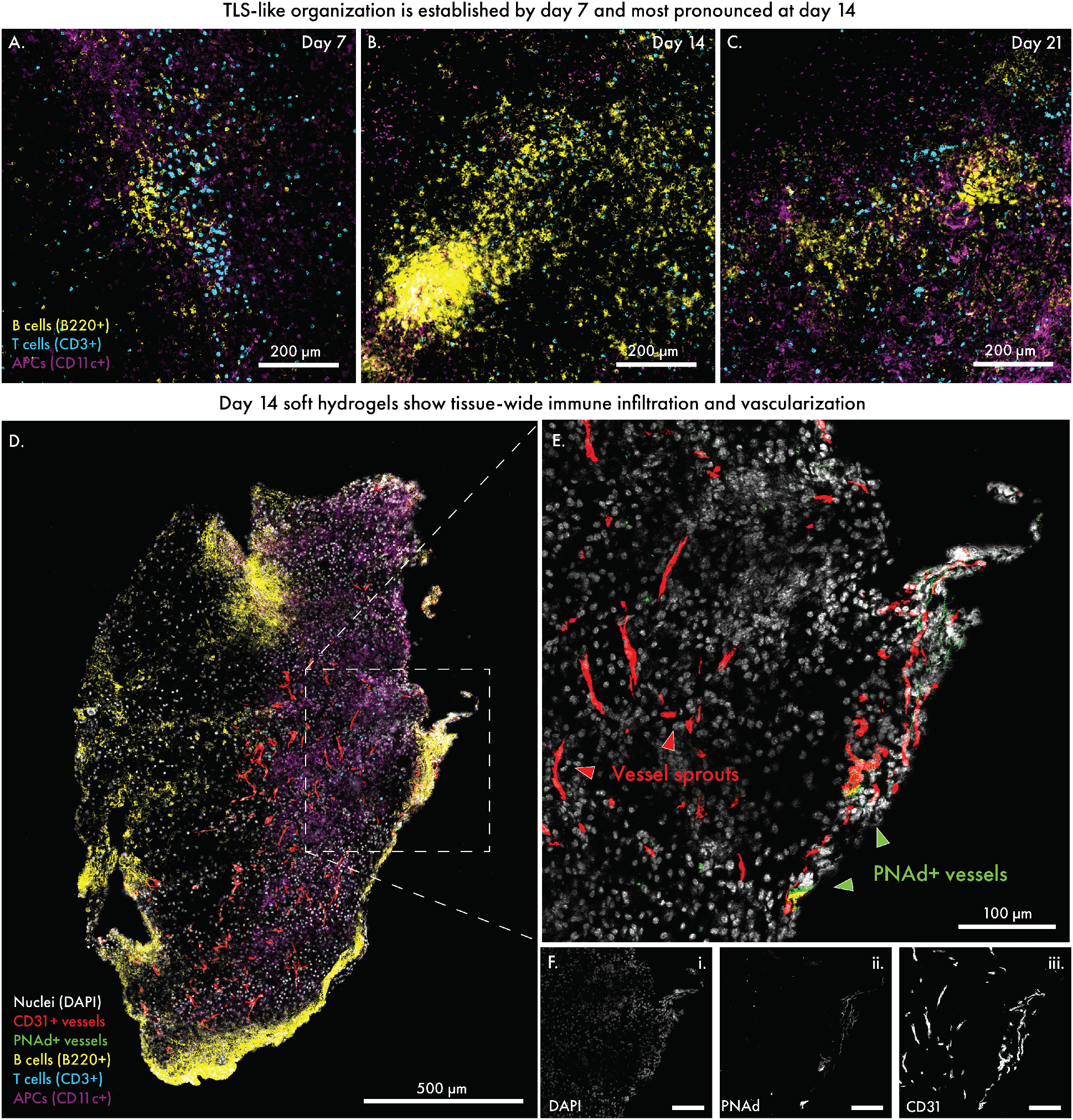
Soft hydrogels support temporal evolution of lymphoid organization and vascularization in vivo. Soft hydrogels containing ovalbumin (OVA; 35 μg), CCL21 (0.4 μg), and LIGHT (0.4 μg) (OCL) were implanted subcutaneously in BALB/c mice and retrieved at days 7, 14, and 21. Explants were sectioned through their central plane for spatial characterization. (**A–C**) Representative immunofluorescence images of soft-hydrogel explants collected at day 7 (A), day 14 (B), and day 21 (C) and stained for B cells (B220⁺, yellow), T cells (CD3⁺, cyan), and CD11c⁺ antigen-presenting cells (magenta). Early infiltrates at day 7 contained small, loosely organized lymphocyte aggregates. B-cell-rich aggregates were most prominent at day 14 and remained detectable at day 21. Scale bars, 200 μm. (**D**) Multiplexed IBEX imaging of a representative central-plane section from a day-14 soft-hydrogel explant stained for nuclei (DAPI, gray), CD31⁺ vessels (red), PNAd (green), B cells (B220⁺, yellow), T cells (CD3⁺, cyan), and CD11c⁺ antigen-presenting cells (magenta). The image was generated through three sequential staining and imaging rounds registered using DAPI as a common spatial landmark. Dashed box denotes the region enlarged in (E–F). Scale bar, 500 μm. (**E**) Higher-magnification view of the same registered field of view from the boxed region in (D). Red arrowheads indicate CD31⁺ vessel sprouts, and green arrowheads indicate PNAd⁺CD31⁺ HEV-like vessels. Scale bar, 100 μm. (**F**) Single-channel images from the same field of view shown in (E) showing (i) DAPI, (ii) PNAd, and (iii) CD31. Single-channel panels are displayed at reduced size relative to (E); scale bars, 100 μm.

Vascularization evolved over the same interval. CD31⁺ vascular sprouts were initially concentrated near the explant periphery but extended throughout the tissue by day 14 (**Fig. 4D**, **Supplementary Fig. G**). Multiplexed imaging at day 14 further identified PNAd signal colocalized with CD31⁺ vascular structures, suggestive of HEV-like vasculature (**Fig. 4E**–**F**). In secondary and ectopic lymphoid tissues, HEVs support lymphocyte entry and are associated with organized lymphoid regions^48,49^.

Together, lymphoid organization, vascular ingrowth and matrix remodeling developed in parallel within soft hydrogels, with the most pronounced TLS-like architecture at day 14 coinciding with widespread vascularization and extensive loss of the original hydrogel. This suggests that organization developed during, rather than after, active material replacement. The subsequent contraction of B cell-rich regions despite established vascularization may reflect early involution in this tumor-free setting as continued hydrogel remodeling progressively depleted the local antigen and cytokine reservoir.

### 3. Soft hydrogels support a highly organized TLS-like state by day 14

To define the cellular transition underlying this spatial progression, we profiled soft OCL hydrogels longitudinally by flow cytometry. Total live cell recovery remained high during the first two weeks, increasing from 25,200 ± 4,300 cells at day 7 to 36,200 ± 7,900 cells by day 14, before falling to 2,100 ± 300 cells by day 21 (p = 0.0105; **Fig. 5A**). Over the same interval, the log_10_ lymphoid-to-myeloid ratio shifted from -0.93 ± 0.2 at day 7 to 0.04 ± 0.11 at day 14 and 0.57 ± 0.18 by day 21 (**Fig. 5B**), capturing a progressive transition from an early myeloid-rich infiltrate toward lymphoid dominance.

**Figure 5.**
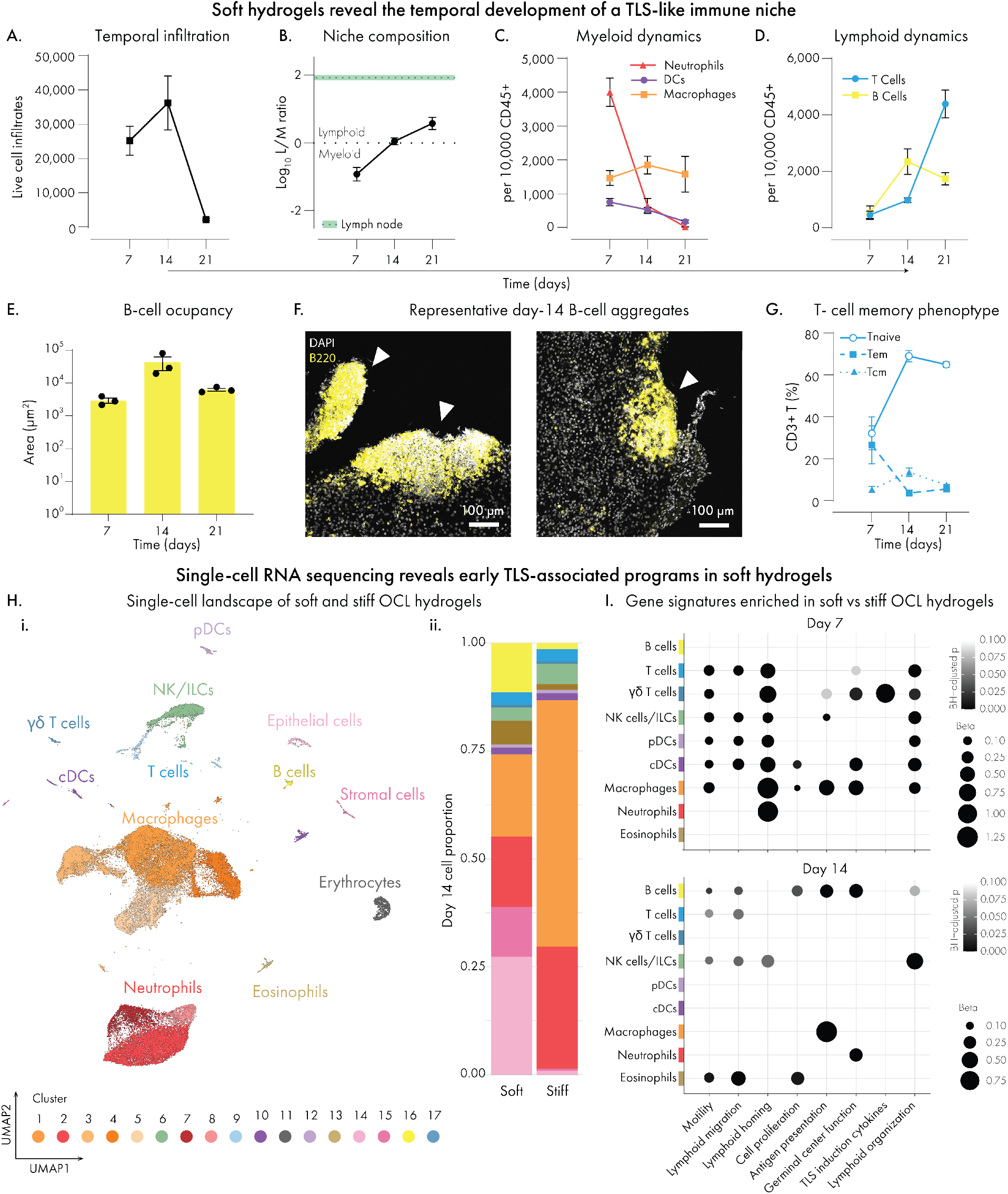
Temporal evolution and transcriptional organization of TLS-like lymphoid niches in soft OCL hydrogels. Soft OCL hydrogels were profiled longitudinally by flow cytometry and immunofluorescence imaging (A–G), and soft and stiff OCL hydrogels were analyzed by single-cell RNA sequencing at days 7 and 14 (H–I). (**A**) Total live-cell recovery from soft OCL hydrogel explants at days 7, 14, and 21. (**B**) Log₁₀-transformed lymphoid-to-myeloid ratio over time. Lymphoid cells were defined as B and T cells, whereas myeloid cells comprised macrophages, neutrophils, and dendritic cells. The green band indicates the mean ± SEM calculated from lymph node samples using the same population definitions and transformation (n = 5 biologically independent mice). (**C**) Neutrophils, dendritic cells, and macrophages per 10,000 CD45⁺ cells over time. (**D**) T and B cells per 10,000 CD45⁺ cells over time. (**E**) B-cell occupancy within manually outlined lymphoid aggregates, quantified as the thresholded B220⁺ area at each timepoint. All identifiable B-cell-rich aggregates within each analyzed hydrogel section were quantified and averaged to obtain a single value per hydrogel. Each point represents one biologically independent hydrogel (n = 3 per timepoint). (**F**) Representative B-cell-rich aggregates within soft OCL hydrogels at day 14. B220, yellow; DAPI, gray. Scale bars, 100 μm. (**G**) Frequencies of naïve, effector-memory (Tem), and central-memory (Tcm) T cells among CD3⁺ cells over time. (**H**) Single-cell landscape and composition of soft and stiff OCL hydrogels. (**i**) Uniform Manifold Approximation and Projection (UMAP) of 69,350 cells recovered from soft and stiff OCL hydrogels at days 7 and 14 after quality control (Supplementary Fig. 11). (**ii**) Relative cell-type composition of soft and stiff OCL hydrogels at day 14. Single-cell RNA-sequencing analysis included n = 3 independently processed replicates per condition, each comprising 1-4 hydrogel explants (Supplementary Table 3); mean ± SEM shown in Supplementary Fig. 12. (**I**) Cell type-specific enrichment of transcriptional gene signatures in soft relative to stiff OCL hydrogels at day 7 (top) and day 14 (bottom). Signatures encompassed motility, lymphoid migration, lymphoid homing, cell proliferation, antigen presentation, germinal-center function, TLS-induction-associated cytokines, and lymphoid organization. Signature definitions and gene sets are provided in Supplementary Table 4. β coefficients and FDR-adjusted p values were obtained by fitting signature scores as a function of hydrogel condition and experiment using Gaussian regression within each cell type. Positive β coefficients indicate signature enrichment in soft relative to stiff hydrogels. Only signatures enriched in soft hydrogels with a Benjamini–Hochberg-adjusted *p* value < 0.1 and model coefficient β > 0.1 are shown. Dot size indicates the magnitude of the model coefficient (β), and color indicates the Benjamini-Hochberg-adjusted *p* value. Data in (A–E) and (G) are presented as mean ± SEM. For longitudinal flow-cytometry analyses, n = 5 biologically independent explants at days 7 and 14 and n = 3 at day 21. For (E), statistical comparisons were performed using one-way ANOVA followed by Tukey’s multiple-comparisons test. Exact *p* values are shown where indicated.

Analysis of the myeloid compartment revealed a transient neutrophil-rich phase. Neutrophils accounted for 40 ± 4% of CD45⁺ cells at day 7, declined approximately 6-fold by day 14, and were nearly undetectable by day 21 (**Fig. 5C**). Macrophages persisted at lower levels, whereas dendritic cells gradually declined. Absolute counts of all three populations decreased after day 7, indicating a contraction of the myeloid infiltrate rather than proportional redistribution alone (**Supplementary Fig. 10**). As the myeloid compartment contracted, lymphoid populations increased with distinct kinetics. B cells rose from 5 ± 2% of CD45⁺ cells at day 7 to 22 ± 5% at day 14, when they were approximately twice as abundant as T cells. By day 21, T cells dominated the lymphoid compartment at 44 ± 5%, while B cells remained at 17 ± 2% (**Fig. 5D**).

The transient B-cell-rich phase coincided with the most pronounced spatial organization. To quantify this organization from immunofluorescence images, we measured B-cell occupancy, defined as the total B220⁺ area within each aggregate, and aggregate compactness, defined as the fraction of aggregate area occupied by B220⁺ signal. B-cell occupancy increased nearly 20-fold between days 7 and 14 to 43,500 ± 19,300 μm² and then decreased by day 21 (**Fig. 5E**–**F**). B-cell aggregate compactness followed a similar trend, increasing from 6.4 ± 1.3% at day 7 to 51 ± 10.5% at day 14 (p = 0.0062; **Supplementary Fig. 10H**). Day-14 aggregates also reached a tissue-relevant spatial scale, with a mean cross-sectional area of 78,300 ± 19,300 μm², comparable to the median area reported for early TLSs in a recent pan-cancer analysis^17^ (**Supplementary Fig. 10I**). In parallel, the T-cell compartment shifted away from an early effector-memory phenotype. Effector-memory T cells (CD62L⁻CD44⁺) declined from 27 ± 9% to 4 ± 1% between days 7 and 14 (p = 0.0219), whereas naïve T cells (CD62L⁺CD44⁻) increased from 32 ± 8% to 69 ± 3% (p = 0.0003) (**Fig. 5G**; **Supplementary Fig. 10J**–**M**). This shift toward a naïve phenotype is consistent with the emergence of a more lymphoid-like trafficking environment, in which CD62L⁺CD44⁻ naïve T cells preferentially home to organized lymphoid tissue^48^.

To define the transcriptional programs accompanying this day-7-to-day-14 transition, we performed single-cell RNA sequencing of pooled soft and stiff OCL hydrogels at both timepoints (**Supplementary Table 3**). After quality control, 69,350 cells were retained across the four conditions (**Fig. 5Hi**; **Supplementary Fig. 11**). At day 14, soft hydrogels showed greater B-cell representation than stiff hydrogels (q = 0.0255), whereas neutrophils (q = 0.0126) and macrophages (q < 0.0001) were more prominent in stiff hydrogels (**Fig. 5Hii**; **Supplementary Fig. 12A**). Soft hydrogels also contained a greater representation of stromal (q = 0.0162) and epithelial populations (q < 0.001). Given the extensive remodeling and integration of soft hydrogels with surrounding subcutaneous tissue by this timepoint, these populations may reflect both cells associated with the remodeled implant and adjacent host tissue recovered during explantation.

We next scored transcriptional gene signatures encompassing motility, lymphoid migration and homing, proliferation, antigen presentation, germinal-center function, lymphoid organization and TLS-induction-associated cytokines (**Fig. 5I**; **Supplementary Table 4**). Within each cell type, signature scores were compared between soft and stiff hydrogels, by fitting a Gaussian model of scores as a function of condition and experiment, such that positive β coefficients indicate higher scores (enrichment) in soft hydrogels relative to stiff hydrogels. At day 7, soft hydrogels were already enriched for multiple programs associated with lymphoid recruitment and organization across multiple immune populations. Lymphoid-homing signatures were strongest in macrophages and neutrophils, and were also enriched in cDCs, pDCs, NK cells/ILCs, and T-cell populations. Lymphoid-organization signatures were similarly enriched across macrophages, dendritic-cell, NK/ILC and T-cell populations, but not neutrophils, while γδ T cells showed prominent enrichment for TLS-associated cytokine signatures. B cells showed no material-dependent enrichment at this early stage.

By day 14, however, enrichment had broadened into the B-cell compartment. B cells in soft hydrogels showed higher scores for lymphoid migration, cell proliferation, antigen presentation, germinal-center function and lymphoid organization, representing the broadest set of enriched signatures among the populations examined at this timepoint. Macrophages retained antigen presentation enrichment, while NK cells/ILCs continued to show selected lymphoid-organizing programs.

Across both timepoints, germinal-center-function enrichment in non-B-cell populations was driven largely by *BclC*, *Cxcr4* and *Fas*, genes with roles in immune-cell trafficking, activation and survival.

Because these comparisons were performed between materials at each timepoint, they do not establish how the soft niche itself changed between days 7 and 14. We therefore directly compared soft hydrogels across these timepoints (**Supplementary Fig. 12B**). Lymphoid-homing signatures were significantly enriched at day 7 in pDCs, macrophages, and neutrophils, while lymphoid-organization and TLS-associated cytokine signatures were enriched across T cells, B cells and several innate populations. By day 14, antigen-presentation programs were significantly enriched in pDCs, and showed higher scores in B cells. Thus, the soft niche transitioned from an early recruitment and organizing program toward a more antigen-processing and lymphoid-focused state by day 14.

Consistent with that transition, CD95⁺GL7⁺ GC-like B cells increased from 0.2 ± 0.1% at day 7 to 1.1 ± 0.2% at day 14, reaching approximately half the frequency observed in lymph nodes from the same soft OCL cohort (2.1 ± 0.1%), and remained detectable at day 21 (**Supplementary Fig. 12C**). OVA-specific serum IgG responses were also higher at day 14 relative to bolus delivery of the same cargo and OVA-only soft hydrogels **(Supplementary Fig. 12D**–**G)**, indicating that the locally organized niche was accompanied by antigen-specific humoral activity.

Together with the imaging and flow-cytometry data, these results track the longitudinal evolution of the immune niche in soft hydrogels in which early lymphoid-homing and TLS-induction programs at day 7 precede the organized, B-cell-rich TLS-like state observed at day 14. The earliest material-dependent divergence was already evident at day 7 and was concentrated predominantly within myeloid populations.

### 4. Early interferon-associated myeloid remodeling distinguishes the soft niche

Because the earliest material-dependent divergence was concentrated within myeloid populations at day 7, we next asked whether the early innate response could help explain the divergent immune trajectories of soft and stiff hydrogels. We compared empty and OCL-loaded formulations at this timepoint to distinguish material-dependent from cargo-dependent features of early immune recruitment.

At day 7, macrophages, neutrophils and dendritic cells together comprised a combined 85 ± 3% of CD45⁺ cells in soft OCL hydrogels and 78 ± 3% in stiff OCL hydrogels (**Fig. 6A**). By day 21, this combined compartment had fallen to 18 ± 5% in soft OCL hydrogels but remained at 56 ± 2% in stiff hydrogels. Comparison with empty controls suggested that the similar myeloid-dominated states observed in soft and stiff OCL hydrogels at day 7 arose through different recruitment drivers. Stiff hydrogels remained myeloid-dominant regardless of cargo, whereas OCL loading increased the myeloid fraction significantly in soft hydrogels from 59 ± 4% to 85 ± 3% (p < 0.0001; **Supplementary Fig. 13A**).

**Figure 6.**
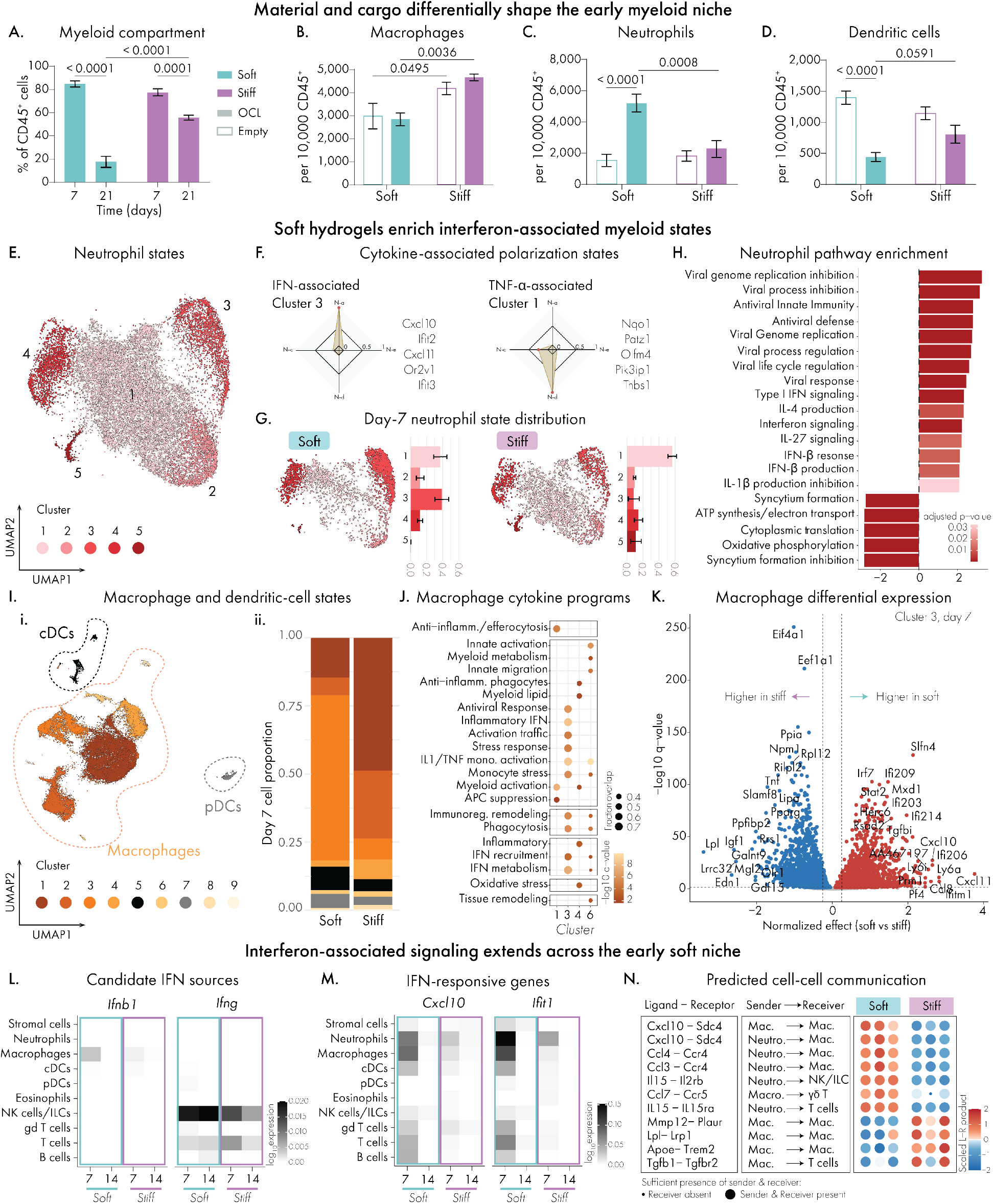
Early material-dependent myeloid remodeling and interferon-associated immune programs in TLS-inducing hydrogels. Flow cytometry was used to quantify myeloid composition and abundance in (A–D), whereas single-cell RNA sequencing was used to profile molecular states in (E–N). (**A**) Combined macrophage, neutrophil, and dendritic-cell compartment as a percentage of CD45⁺ cells in soft and stiff OCL hydrogels at days 7 and 21. (**B**–**D**) Day-7 abundance of macrophages (**B**), neutrophils (**C**), and dendritic cells (**D**) per 10,000 CD45⁺ cells in empty and OCL-loaded soft and stiff hydrogels. (**E**) UMAP of reclustered neutrophils from soft and stiff OCL hydrogels at days 7 and 14, resolving five transcriptionally distinct populations. (**F**) Immune Response Enrichment Analysis^50^ and representative cluster-associated genes, identifying Cluster 3 as IFN-associated and Cluster 1 as TNF-α-associated. (**G**) Distribution and relative representation of neutrophil transcriptional states (Leiden clusters) in soft and stiff OCL hydrogels at day 7. Bars represent the mean ± SEM across n = 3 independently processed biological pools per condition. (**H**) Gene set enrichment analysis (GSEA) of day-7 neutrophils from soft versus stiff OCL hydrogels. Positive normalized enrichment scores indicate pathways associated with soft hydrogels, whereas negative scores indicate pathways associated with stiff hydrogels; bar color indicates Benjamini–Hochberg-adjusted *p* value. (**I**) Reclustering of macrophages, conventional dendritic cells (cDCs), and plasmacytoid dendritic cells (pDCs). (**i**) UMAP resolving seven macrophage populations together with distinct cDC and pDC populations. (**ii**) Relative representation of the reclustered macrophage, cDC, and pDC populations in soft and stiff OCL hydrogels at day 7. (**J**) Enrichment of monocyte-derived Cytokine-Induced Programs across macrophage clusters. Dot size indicates the overlap between each program and the cluster-specific DEG set, and color indicates the -log_10_-transformed Benjamini-Hochberg-adjusted *q* value. (**K**) Volcano plot of condition-specific differential expression within day-7 macrophage Cluster 3 comparing soft and stiff OCL hydrogels. Positive effect sizes indicate higher expression in soft hydrogels, whereas negative values indicate higher expression in stiff hydrogels; selected differentially expressed genes are labeled. (**L**) Mean normalized expression of *Ifnb1* and *Ifng* across cell populations and conditions. (**M**) Mean normalized expression of the IFN-responsive genes *Cxcl10* and *Ifit1* across cell populations and conditions. (**N**) Selected MultiNicheNet-predicted ligand–receptor interactions originating from innate sender populations in soft and stiff OCL hydrogels at day 7. Interactions were curated from prioritized differential ligand–receptor interactions shown in Supplementary Fig. 16 after the application of the expression, cell-abundance, and differential-expression filtering criteria described in the Methods. Color indicates the scaled ligand–receptor pseudobulk expression across n = 3 independently processed biological pools per condition. For day-7 flow-cytometry analyses in (A–D), n = 5 independently analyzed hydrogel explants per condition were obtained from n = 3 mice. Day-21 flow-cytometry analysis in (A) included n = 3 biologically independent soft hydrogel explants and n = 5 biologically independent stiff hydrogel explants. Data are presented as mean ± SEM. Statistical significance was assessed using two-way ANOVA followed by Šídák’s multiple-comparisons test. Exact *p* values are shown where indicated.

Macrophage and neutrophil recruitment showed distinct material- and cargo-dependent dynamics. Macrophages were enriched under stiff conditions, comprising 47 ± 2% of CD45⁺ cells in stiff OCL hydrogels versus 28 ± 3% in soft OCL hydrogels (p = 0.0036; **Fig. 6B**), with the same trend observed in empty controls, indicating that macrophage enrichment was primarily material dependent. Absolute cell counts nevertheless increased approximately 2.3-fold following OCL loading in stiff hydrogels (28,068 ± 6,347 to 64,661 ± 9,116 cells; p = 0.0003), showing that cargo increased macrophage accumulation without substantially changing its relative representation (**Supplementary Fig. 13B**).

Neutrophils instead showed selective cargo-dependent enrichment within soft hydrogels. OCL increased neutrophil representation by approximately 3.4-fold, from 15 ± 4% to 52 ± 6% of CD45⁺ cells (p < 0.0001), compared with 23 ± 5% in stiff OCL hydrogels (**Fig. 6C**). Absolute neutrophil counts increased in both materials following OCL loading, indicating that the selective proportional enrichment in soft hydrogels reflected a broader shift in immune composition rather than neutrophil recruitment alone (**Supplementary Fig. 13C**). Dendritic cell frequency decreased in both materials following OCL loading without a corresponding reduction in absolute counts, indicating that the proportional decrease largely reflected increased abundance of other populations (**Fig. 6D**; **Supplementary Fig. 13D**).

Day-7 neutrophils showed a prominent material-dependent transcriptional divergence, characterized in soft hydrogels by an interferon-associated state. Reclustering of neutrophils from the single-cell dataset resolved five transcriptional states (**Fig. 6E**; **Supplementary Fig. 14**), of which Immune Response Enrichment Analysis^50^ identified three distinct cytokine-associated polarization states: Cluster 1 with a TNF-α response, Cluster 2 with an IL-1 response, and Cluster 3 with an IFN response (**Fig. 6F**). At day 7, the IFN-responsive state represented 39 ± 9% of soft-hydrogel neutrophils but only a small fraction of the stiff population, whereas the TNF-responsive state predominated in the stiff hydrogels (**Fig. 6G**). IL-1-responsive neutrophils represented a smaller and comparable fraction in both materials. Consistent with this polarization, gene set enrichment analysis (GSEA)^51^ showed enrichment of type I interferon signaling, interferon-responsive and antiviral innate immune programs in soft-hydrogel neutrophils (**Fig. 6H**). Stiff-hydrogel neutrophils instead favored oxidative phosphorylation, ATP synthesis-coupled electron transport and cytoplasmic translation, together with gene sets associated with the regulation of syncytium formation. IFN-responsive neutrophil states have been reported in diverse inflammatory contexts^52–54^, and here they represent a distinguishing feature of the early soft niche.

Macrophages showed a parallel material-dependent split. Reclustering resolved seven macrophage populations together with cDC and pDC populations (**Fig. 6I**; **Supplementary Fig. 15A**–**C**). Cytokine-Induced Program^55^ enrichment analysis showed that Cluster 3 macrophages, which predominated in soft hydrogels at day 7 (61 ± 6%), were enriched for antiviral and interferon programs, immune-cell trafficking, IL-1/TNF-associated monocyte activation, phagocyte-complement activity and immunoregulatory remodeling. In contrast, Cluster 1 macrophages, which predominated in stiff hydrogels (49 ± 7%), showed higher anti-inflammatory/efferocytosis, myeloid-activation and APC-suppression scores (**Fig. 6J**). Within Cluster 3, soft hydrogels showed differential upregulation of interferon-response genes *Ifi203*, *Ifi20C*, *Ifi20S*, *Ifi214, Ifitm1*, *Slfn4* and *LyCa*, together with IFN-inducible chemokines *Cxcl10* and *Cxcl11* (**Fig. 6K**), whereas stiff hydrogels showed upregulation of *Tnf*, *Slamf8*, *Lpl*, *Igf1* and *Pparg*, separating an interferon-associated macrophage state from that favored by the persistent stiff material.

Candidate interferon sources were then examined within the early niche. *Ifnb1* expression was detected primarily in macrophages in soft hydrogels and concentrated within the dominant IFN-associated macrophage population, with additional strong expression in a rare Cluster 6 macrophage state characterized by mixed IFN-associated, inflammatory, and tissue-remodeling programs (**Fig. 6L**; **Supplementary Fig. 15D**). Together, these findings support macrophages in soft hydrogels as a candidate local source of IFN-I. *Ifng*, representing a distinct interferon pathway, was instead expressed primarily by NK/ILC cells and was not restricted to the soft condition. Consistent with the broader interferon-responsive state, *Cxcl10* and *Ifit1* were most highly expressed at day 7 in soft hydrogels, particularly in neutrophils and macrophages but also across T and B cells (**Fig. 6M**). Both IFN-I and IFN-γ have been implicated in TLS development^56,57^, but the material-dependent distinction in our system was most evident in the early IFN-I-associated myeloid response.

This divergence also extended to broader cell-cell communication within the early niche. To determine whether the innate populations distinguishing the two materials engaged different signaling networks, MultiNicheNet^58^ was used to compare interactions arising from selected innate sender populations in soft and stiff day-7 hydrogels (**Supplementary Fig. 16**), most involving myeloid senders or receivers. In soft hydrogels these included *Cxcl10*- and *Ccl3/Ccl4*-associated signaling to macrophages, *Ccl7*/*Ccr5* signaling from macrophages to γδ T cells, and IL-15 signaling from neutrophils to T cells and NK/ILCs, consistent with a network emphasizing immune recruitment and myeloid-lymphoid crosstalk (**Fig. 6N**). Stiff hydrogels instead showed prominent macrophage-centered *Apoe*-*Trem2*, *Lpl*-*Lrp1* and *Mmp12*-*Plaur* signaling, associated with lipid handling and tissue remodeling, together with predicted *Tgfb1*-*Tgfbr2* signaling from macrophages to T cells. Thus, networks reflected the broader distinction between an immune-recruiting soft niche and a remodeling-dominant stiff niche. These interactions provide additional candidate mechanisms through which the early myeloid response may shape the subsequent divergence in TLS-like organization.

Concurrent type I interferon and LTβR-associated signaling has been linked to the development of mature B-cell-rich TLSs^59^. In the OCL system, LIGHT can engage LTβR, which was broadly expressed across myeloid and stromal populations in both materials (**Supplementary Fig. 17A**). HVEM (*Tnfrsf14*), an alternative LIGHT receptor with a more lymphoid-associated expression pattern, was most prominent in T and γδ T cells at day 14 in soft hydrogels. Endogenous *Ltb* and *Tnfsf14* were also detectable across immune populations (**Supplementary Fig. 17B**), indicating that both niches contained components capable of supporting LTβR-associated signaling. Against this shared background, soft hydrogels developed a prominent early IFN-I-associated myeloid state characterized by interferon-responsive macrophages and neutrophil programs, with macrophages emerging as a candidate local source of IFN-I and broader interferon-responsive activity extending across the niche. Together, these features provide a potential mechanistic link between the early soft-hydrogel myeloid state and subsequent B-cell-rich TLS-like organization, while the upstream trigger responsible for macrophage IFN-I production remains unresolved.

### 5. Engineered TLS-like niches support prophylactic tumor control

Having established that soft and stiff hydrogels generate distinct immune niches, we next asked whether these differences translated into antitumor activity. C57BL/6J mice received soft and stiff OCL hydrogels, or corresponding empty controls, 14 days before challenge with antigen-matched B16F10-OVA melanoma in the same anatomical compartment, allowing for the engineered niches to form before challenge. Anti-PD-L1 treatment began 14 days after tumor inoculation (**Fig. 7A**) and continued twice-weekly throughout the study.

**Figure 7.**
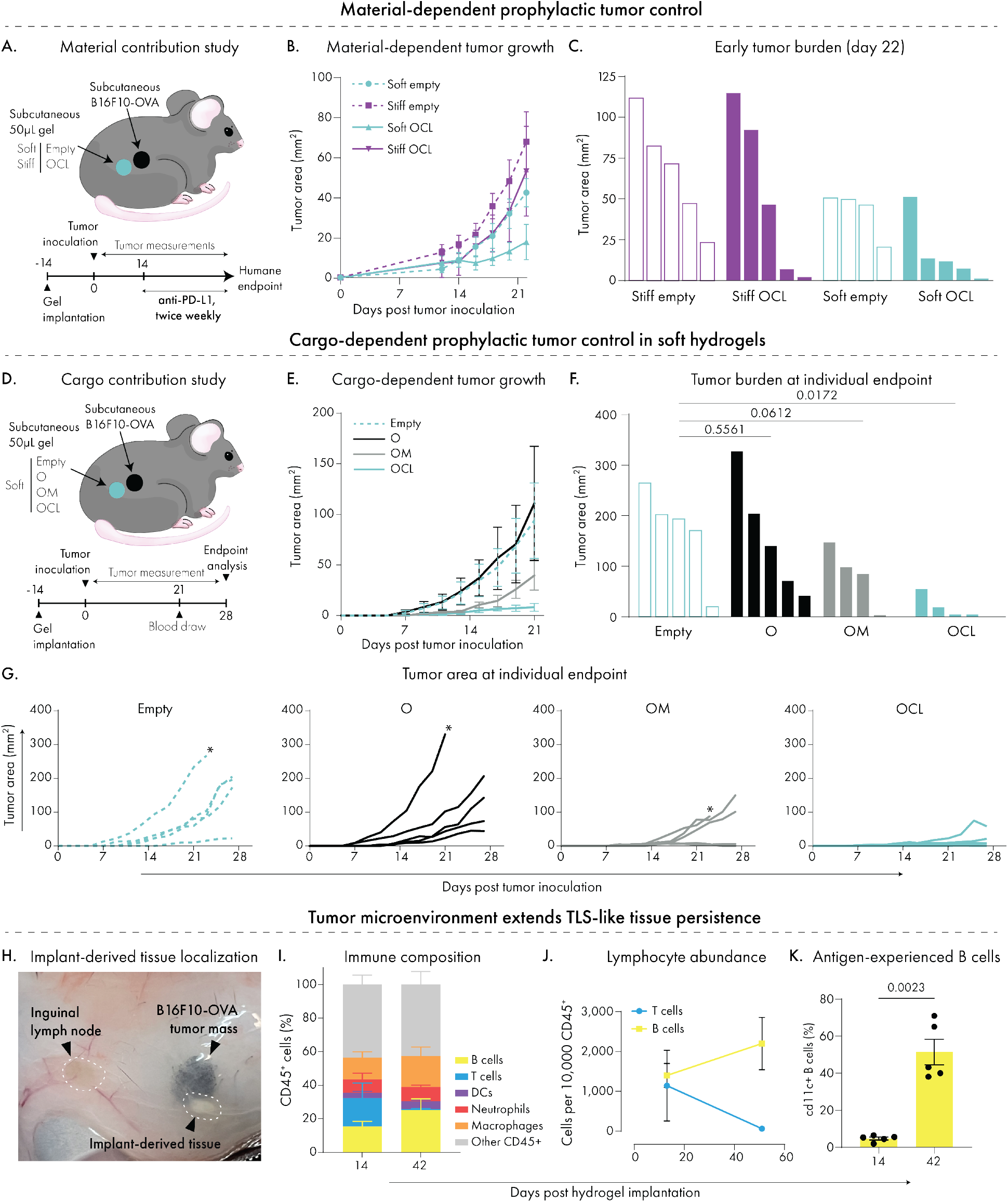
Prophylactic antitumor activity and post-challenge remodeling of engineered TLS-like tissues. (**A**) Experimental schematic comparing soft and stiff empty or OCL hydrogels implanted subcutaneously 14 days before B16F10-OVA tumor challenge. Anti-PD-L1 was administered twice weekly beginning 14 days after tumor inoculation. (**B**) Tumor growth in mice receiving soft empty, stiff empty, soft OCL, or stiff OCL hydrogels. (**C**) Individual tumor areas at day 22 from the material-comparison prophylaxis study shown in (A–B). (**D**) Experimental schematic comparing cargo formulations within soft hydrogels. Empty, OVA-only (O), OVA plus MPLA (OM), or OCL hydrogels were implanted 14 days before B16F10-OVA tumor challenge. Blood was collected at day 21, and endpoint analyses were performed at day 28 after tumor inoculation. (**E**) Tumor growth across soft-hydrogel cargo formulations. (**F**) Individual tumor areas at day 28 or at the humane endpoint for mice euthanized before the scheduled endpoint. (**G**) Individual tumor-growth trajectories across cargo formulations. Asterisks indicate animals euthanized before the scheduled day-28 endpoint. (**H**) Representative endpoint image showing the spatial relationship among the B16F10-OVA tumor mass, draining inguinal lymph node, and implant-derived tissue in a soft OCL-treated mouse. (**I**) Immune composition of soft OCL-derived tissues collected from parallel cohorts at day 14 after implantation, before tumor challenge, and at day 42 after implantation, corresponding to 28 days after tumor challenge. (**J**) B- and T-cell abundance per 10,000 CD45⁺ cells within soft OCL-derived tissues at days 14 and 42 after implantation. (**K**) CD11c⁺ B cells as a percentage of total B cells within soft OCL-derived tissues at days 14 and 42. Data are presented as mean ± SEM. For the soft-versus-stiff prophylaxis study (A–C), n = 4 biologically independent mice in the soft-empty group and n = 5 mice in each of the remaining groups. For the cargo-comparison study (D–G), n = 5 mice per group at study initiation. For tissue analyses in (I–K), n = 5 biologically independent mice per time point; day-42 tissues were recovered from the OCL-treated cohort shown in (D-G). Statistical significance in (F) was assessed using one-way ANOVA followed by the two-stage Benjamini, Krieger and Yekutieli multiple-comparisons procedure, with comparisons made against the empty-hydrogel group. Values shown in (F) are FDR-adjusted *q* values. Statistical significance in (K) was assessed using an unpaired two-tailed Student’s t-test. Exact *q* and *p* values are shown where indicated.

Soft OCL hydrogels improved early tumor control. Over the first three weeks following tumor challenge, mice receiving soft OCL hydrogels carried the lowest tumor burden of any group, with a mean tumor area of 17.8 ± 8.8 mm^2^ at day 22 compared with 53.2 ± 22.5 mm^2^ for stiff OCL hydrogels, which more closely resembled empty controls (**Fig. 7B**–**C**). The separation narrowed at later timepoints and survival did not differ significantly between soft and stiff OCL formulations, although survival curves for both OCL groups were delayed relative to their corresponding empty controls (**Supplementary Fig. 18A**–**B**). Systemic antibody responses did not track with this protection. In a separate immunization study, soft and stiff OCL hydrogels produced comparable OVA-specific serum IgG through week 2, after which titers were higher in the stiff cohort (p = 0.0168; **Supplementary Fig. 18C**). The prolonged persistence of stiff hydrogels may have extended antigen exposure and APC engagement^60^, however, this greater antibody response was not accompanied by superior tumor control. Because these measurements were obtained in separate studies, they do not establish a direct mechanistic relationship, but they indicate that systemic antibody magnitude alone does not account for the early protection associated with soft OCL hydrogels.

To isolate antitumor activity arising from the hydrogel formulation itself, we next removed checkpoint blockade and varied cargo within the soft formulation. Empty, OVA-only (O), OVA plus MPLA (OM), and OCL hydrogels were implanted 14 days before tumor challenge (**Fig. 7D**), allowing us to compare the complete TLS-inducing formulation against antigen alone and conventional adjuvanted vaccine within the same material context. OCL produced the lowest endpoint tumor burden, with a mean tumor area of 18.8 ± 10.4 mm^2^ compared to 172.7 ± 50.7 mm^2^ for empty control (q = 0.0172; **Fig. 7E**–**F**). OVA alone provided no measurable benefit, whereas OM produced an intermediate response (68.7 ± 28.8 mm^2^, q = 0.0612 vs empty). All OCL-treated mice remained below the humane endpoint whereas one OM-treated mouse required earlier euthanasia (**Fig. 7G**). Measured systemic responses again did not mirror tumor control. Both OCL and OM increased OVA-specific IgG relative to empty controls, with the larger response observed in OM, and responses were predominantly IgG1 (**Supplementary Fig. 18D**–**E**). Splenic T-cell profiles likewise did not clearly distinguish OCL from OM, both showing increased CD4:CD8 ratios and reduced PD-1⁺LAG-3⁺ CD4⁺ T cell frequencies relative to empty controls (**Supplementary Fig. 18F**–**G**). Thus, the greater tumor control observed with OCL was not mirrored by the measured systemic humoral or splenic T-cell responses. Although systemic immunity may still contribute to the overall antitumor effect ^61^, these findings point to a potential additional local contribution from the TLS-like niche.

The engineered TLS-like tissue also persisted and remained dynamic after tumor challenge. At the day-42 endpoint, residual OCL-derived tissues were identifiable at all implantation sites (5/5), compared with 1/4 OVA-only and 2/4 OM sites (**Fig. 7H**; **Supplementary Fig. 1GA**). This was more frequent than in the tumor-free day-21 study (60%), raising the possibility that the tumor-bearing environment contributes to maintenance of the engineered OCL-derived tissue. Comparing OCL-derived tissues at day 42, after 28 days of tumor exposure, with soft OCL 14 immediately before challenge showed increased B-cell representation in both relative abundance and absolute number (**Fig. 7I**–**J**; **Supplementary Fig. 1GB**–**C**). Germinal center-associated B cells declined over this interval, while CD11c⁺ B cells expanded from 5 ± 1% of B cells at day 14 to 43 ± 7% by day 42 (p = 0.0023), and dendritic cell numbers within OCL-derived tissues remained stable over this interval (**Fig. 7K**; **Supplementary Fig. 1GD**–**E**). CD11c-expressing B cells have been described as memory-like states arising outside germinal centers under sustained antigenic or inflammatory stimulation, and can exhibit potent antigen-presenting activity^62–64^. Although these data do not establish that persistence or continued remodeling of the engineered tissue directly mediates tumor control, they show that the engineered niche remains immunologically dynamic after tumor challenge, with sustained local tissue persistence and substantial remodeling of its B-cell compartment.

## Conclusions and Outlook

Our results indicate that hydrogel mechanical properties regulated not only material persistence but the identity and organization of the immune niche that formed within it, making the scaffold an active participant in TLS neogenesis rather than a passive vehicle for soluble cues. Despite release of identical TLS-associated cues, only soft hydrogels underwent distributed cellular infiltration and matrix remodeling, whereas stiff hydrogels remained comparatively poorly infiltrated, resisted distributed remodeling and persisted longer *in vivo*. Soft-hydrogel niches subsequently progressed toward vascularized, lymphoid-dominant TLS-like tissues, whereas stiff hydrogels remained myeloid-rich with limited lymphoid organization within the examined timeframe. Single-cell transcriptomics further identified transient interferon-associated neutrophil and macrophage states in the early soft niche that preceded the most pronounced B-cell-rich TLS-like organization observed at day 14.

These findings shift the design question for biomaterial-mediated TLS induction from how long a scaffold can retain TLS-inducing factors to how its viscoelastic properties govern the extent to which host cells can infiltrate, remodel and replace it. Within this supramolecular system, the properties that set resistance to cell-mediated deformation, including yield strain and creep, tracked with both material persistence and the extent of infiltration, and prolonged persistence was not advantageous for TLS neogenesis. Instead, the softer network acted as a sacrificial template that permitted host cells, matrix and vasculature to progressively replace the material as TLS-like architecture emerged. This interpretation remains specific to the mechanical range, material class and subcutaneous context examined here and does not imply that the least persistent scaffold will always be optimal. Rather, effective TLS-inducing materials may require a balance between providing a temporary organizing framework and supporting host-driven remodeling and replacement of the material.

Several questions follow from this view. The first concerns the lifespan of engineered TLS: how long these tissues persist, and whether their longevity depends on continued antigen or other inflammatory cues, remains to be defined, a longitudinal question that tunable platforms such as this one are well suited to address. A second concerns tissue context: because we examined a single subcutaneous compartment, it is unclear whether the requirements for TLS induction differ within tumors or other tissues, and here injectable systems are a particular asset, as they can be introduced across anatomical sites with minimal modification. A third concerns mechanism: although our transcriptomic analyses identified cell populations and states that track with the TLS-inducing outcome, their causal contributions remain to be confirmed. Whether the engineered TLS-like niche contributes through local antigen presentation, coordination with the draining lymph node, or other mechanisms of immune communication also remain unresolved, motivating future studies to directly test its function in therapeutic settings. Lastly, because these relationships were defined within a single class of dynamic supramolecular hydrogels, their generalizability to other injectable materials remains to be established.

More broadly, the liposomal hydrogel platform offers a tractable in situ model in which material properties and immune cues can be varied independently to perturb candidate pathways and define their roles in TLS initiation, maturation, and persistence. Positioning biomaterial design as both a tool for investigating TLS neogenesis and a strategy for engineering local immune tissues, this work provides a framework for developing TLS-inducing biomaterials that could ultimately complement checkpoint blockade, vaccination, and other cancer immunotherapies.

## Materials and Methods

### Polymer and hydrogel fabrication

#### Synthesis of hydrophobically modified HPMC polymers

Hydroxypropyl methylcellulose modified with dodecyl or octadecyl alkyl chains, hereafter referred to as HPMC-C12 and HPMC-C18, respectively, was synthesized as previously described, with minor modifications^65^. Briefly, HPMC (1.0 g; Sigma-Aldrich) was dissolved in 40 mL N-methyl-2-pyrrolidone (NMP; Sigma-Aldrich) by stirring overnight. Either 1-dodecyl isocyanate or 1-octadecyl isocyanate was dissolved in 5 mL NMP and added at 0.52 mmol isocyanate per gram of HPMC together with an equimolar amount of N,N-diisopropylethylamine. The reaction mixture was stirred at 50 °C during reagent addition and subsequently at room temperature for 24 h. Modified HPMC was precipitated in acetone, recovered by centrifugation, and dialyzed against water for 4 days at room temperature using 3.5-kDa molecular-weight-cutoff dialysis tubing. Purified polymers were lyophilized, reconstituted in sterile 1× PBS at 6% w/v, and stored at 4 °C until use. The degree of functionalization was quantified by ^1H nuclear magnetic resonance spectroscopy from the integration of alkyl-chain proton peaks relative to the HPMC backbone.

#### Liposome fabrication

Liposomes were generated using two established methods depending on experimental scale: microfluidic synthesis using an Automated Nanoparticle System (ANP) and thin-film hydration followed by extrusion. Both approaches produced small unilamellar vesicles in the 80–140-nm size range with comparable polydispersity. Microfluidic synthesis was used for all main experiments, whereas extrusion was used for small-volume or rapid-turnaround experiments because of its lower dead volume. Liposomes were composed of 1,2-distearoyl-sn-glycero-3-phosphocholine (DSPC), 1,2-distearoyl-sn-glycero-3-phospho-(1′-rac-glycerol) (DSPG), cholesterol, and 1,2-dimyristoyl-rac-glycero-3-methoxypolyethylene glycol-2000 (DMG-PEG2000) at a molar ratio of 46:12:40:2. All lipids were obtained from Avanti Research. For fluorescent-tracking experiments, the liposome formulation was supplemented with PE-Cy5 at 0.1 mol% (IVIS) or 0.05 mol% (IF).

#### Microfluidic synthesis and tangential-flow filtration

DSPC, DSPG, cholesterol, and DMG-PEG2000 were dissolved in ethanol and combined at a molar ratio of 46:12:40:2. The lipid solution was mixed with 1× PBS at an organic-to-aqueous ratio of 1:4 using a five-input three-dimensional microfluidic chip (Dolomite) installed in an Automated Nanoparticle System (Unchained Labs). Liposomes were subsequently concentrated and diafiltered using a KrosFlo KR2i tangential-flow filtration system. A two-step process consisting of concentration/diafiltration followed by a second concentration step enabled liposomes to be concentrated to up to 10% w/v. Liposome batches were characterized by dynamic light scattering and nanoparticle-tracking analysis.

#### Thin-film hydration and extrusion

Lipids were dissolved in chloroform or methanol, depending on their solubility, and transferred to a round-bottom flask. Organic solvent was removed under vacuum at 40 °C using a rotary evaporator (Büchi) until a uniform lipid film was formed. The film was rehydrated in sterile PBS at a total lipid concentration of 60–150 mg mL⁻¹ under rotary agitation for 30 min at 40 °C. The resulting suspension was manually extruded through 400-, 200-, and 100-nm polycarbonate membranes, with 10 passes through each membrane, using an Avanti Mini-Extruder. Liposomes were used immediately or stored at 4 °C until use.

#### Hydrogel formulation

Hydrogels were formed by combining HPMC-C12 or HPMC-C18 with anionic liposomes to obtain a final polymer-to-liposome concentration ratio of 2:5 w/v. HPMC-C12 and HPMC-C18 formulations are referred to as soft and stiff hydrogels, respectively. OCL hydrogels contained 35 μg ovalbumin (InvivoGen, Cat# vac-pova), 0.4 μg CCL21 (BioLegend, Cat# 586404), and 0.4 μg LIGHT (BioLegend, Cat# 557604) per 50-μL hydrogel. Empty hydrogels contained no added protein cargo. OVA-only hydrogels contained 35 μg ovalbumin, and OVA plus MPLA hydrogels contained 35 μg ovalbumin and 20 μg of MPLA.

### Material characterization

#### Dynamic light scattering and zeta-potential measurements

Liposome hydrodynamic diameter, polydispersity, and zeta potential were measured using a Zetasizer Nano instrument (Malvern Instruments). Samples were diluted 1,000-fold in 0.22-μm-filtered Milli-Ǫ water and analyzed at room temperature. Hydrodynamic diameter was measured in disposable plastic cuvettes and reported as the intensity-weighted particle-size distribution. Zeta potential was measured using DTS1070 folded capillary cells. Three technical measurements were averaged to obtain one value for each independently prepared liposome batch.

#### Nanoparticle-tracking analysis

Liposome particle-size distributions and concentrations were measured by nanoparticle-tracking analysis using a NanoSight NS300. Samples were diluted in 0.22-μm-filtered PBS to obtain an appropriate particle concentration. Three technical measurements were collected and averaged for each independently prepared liposome batch.

#### Rheological characterization

Shear rheology was performed using a stress-controlled DHR-2 rheometer (TA Instruments) equipped with a 20-mm-diameter serrated parallel-plate geometry, humidity chamber, and solvent trap. Approximately 700 μL hydrogel was loaded onto the lower plate and tested at 25 °C using a 700-μm gap. Frequency sweeps were performed from 0.1 to 100 rad s⁻¹ at 1% strain. Oscillatory amplitude sweeps were performed at 10 rad s⁻¹, and the yield strain was defined as the strain at which the storage modulus, G′, and loss modulus, G″, crossed. Steady-shear flow sweeps were performed from high to low shear rates using steady-state sensing. Step-strain measurements incorporated a 120-s recovery interval following each high-strain period. Rheological data were acquired and processed using TRIOS software.

#### Stress-relaxation and creep analysis

Stress-relaxation measurements were performed by applying a constant 10% strain and monitoring the decay in stress over time. Stress values were normalized to the measured stress at 1 s, and the stress-relaxation half-time (τ₁/₂) was defined as the time required for normalized stress to decrease to 0.5.

Creep measurements were performed under a constant applied stress of 12.5 Pa, selected based on previous estimates placing the mechanical stress of the murine subcutaneous space between approximately 12 and 25 Pa^43^. Strain was monitored as a function of time under sustained loading and retained in percent-strain units. The soft hydrogel underwent rapid deformation under sustained loading and reached the measurement limit earlier than the stiff hydrogel, resulting in a shorter creep acquisition window. Steady-state creep strain rates were calculated as the change in percent strain over time between consecutive measurements and are therefore reported as % s⁻¹. Following the initial transient deformation, strain rates were averaged over a region of sustained deformation in which the point-to-point strain rate remained stable (approximately 35 consecutive measurements) to obtain the reported steady-state creep strain rate for each sample.

#### Loss-modulus overshoot analysis

To evaluate nonlinear network rearrangement near yielding, G″ was normalized to its low-strain plateau value within the linear viscoelastic region. The loss-modulus overshoot observed in the stiff formulation was analyzed using the nanoparticle-uncaging framework described by Yu et al.^66^. The normalized G″ peak was fitted with a log-normal distribution, and the associated dissipated energy was calculated from the area under the curve (AUC) of the fitted peak.

#### Protein charge and hydrodynamic-size prediction

Theoretical isoelectric points were calculated using the protein sequence ranges listed in Supplementary Table 2. Charge at physiological pH was classified according to the difference between the theoretical pI and pH 7.4. Proteins were classified as positively charged when pI − 7.4 ≥ 1, negatively charged when pI − 7.4 ≤−1, and approximately neutral when −1 < pI − 7.4 < 1. Effective hydrodynamic diameters were estimated from molecular weight using the empirical mass-to-radius relationship for globular proteins implemented in the FIDA Bio molecular-weight-to-size calculator. These values represent predicted effective hydrodynamic diameters and not experimentally measured structural dimensions.

#### In vitro protein labeling and release

OVA (InvivoGen, Cat# vac-pova), LIGHT (SinoBiological, Cat# 50980-M07B), and CCL21 (BioLegend, Cat# 586404) were fluorescently labeled with NHS-ester Alexa Fluor dyes prior to incorporation into the hydrogel formulations. OVA was labeled with AF647 (Vector Laboratories, Cat# FP-1129-5) using an approximately 5-fold molar excess of dye relative to protein, whereas LIGHT and CCL21 were labeled with AF555 (Vector Laboratories, Cat# FP-1166-5) using an approximately 10-fold molar excess. Sodium bicarbonate was added from a 1 M stock solution at approximately one-tenth of the reaction volume, yielding a final bicarbonate concentration of approximately 0.1 M to provide mildly basic conditions for NHS-ester coupling. Labeling reactions were incubated overnight at 4°C on a tube rotator.

Following labeling, unreacted dye was initially removed using desalting spin columns equilibrated in PBS (ThermoScientific, Cat# 3352585). Columns were centrifuged at 1,500 × *g* for 1 min to remove the storage solution, samples were loaded onto the resin bed, and a 15 µL PBS stacker was added for sample volumes below 70 µL. Labeled protein was recovered by centrifugation at 1,500 × *g* for 2 min. The eluate was subsequently concentrated and further purified by repeated centrifugal diafiltration against PBS using 3-kDa Amicon Ultra centrifugal filters (MilliporeSigma, Cat# UFC500324). Samples were centrifuged at 14,000 × *g* for approximately 30 min per cycle, replenished with PBS, and subjected to approximately 3–4 diafiltration cycles, or until residual free-dye signal was no longer detectable in the filtrate by NanoDrop measurement at the corresponding fluorophore wavelength.

Protein release was evaluated in two separate paired experiments using soft and stiff hydrogel formulations. In each experiment, 100 µL hydrogels contained 7.5 µg OVA together with either 2 µg LIGHT or 2 µg CCL21. OVA and the corresponding TLS-inducing protein were labeled with spectrally distinct fluorophores to permit simultaneous quantification of both cargos within each experiment. Hydrogels (100 µL) were deposited at the bottom of 3 in. glass tubes (2.700 mm inner diameter) sealed at the lower end with epoxy and overlaid with 200 μL PBS. Hydrogels were incubated at 37 °C for 21 days. At designated timepoints, the entire 200 μL release medium was collected and replaced with 200 μL of fresh PBS.

Fluorescence of the released proteins was quantified using a BioTek Synergy Neo2 plate reader. Detection settings were empirically optimized using separate labeled protein standards to minimize spectral crossover between fluorophores. Alexa Fluor 555 fluorescence was measured using 540/10 nm excitation and 580/10 nm emission, with a gain of 100. Following a 20-s delay, Alexa Fluor 647 fluorescence was measured using 650/10 nm excitation and 687/20 nm emission, with a gain of 150. Standard curves were generated using the same fluorescently labeled protein stocks used to prepare the hydrogels and were used to convert fluorescence intensity to released protein mass. Because the complete release medium was exchanged at each sampling point, cumulative release was calculated by summing the protein mass recovered at each successive collection and expressing this value as a percentage of the initial protein mass loaded into the hydrogel.

### In vivo hydrogel studies

#### Animal ethics and husbandry

All animal procedures were approved by the Columbia University Institutional Animal Care and Use Committee (IACUC; protocol no. AC-AABW7658) and were performed in accordance with institutional guidelines. Female BALB/cJ and C57BL/6J mice were obtained from The Jackson Laboratory (#000651 and #000664, respectively). Mice were allowed to acclimate for at least 1 week before experimentation, and animal procurement was timed such that the ages reported for each study correspond to the age at hydrogel implantation. Mice were euthanized according to approved institutional procedures at scheduled experimental endpoints or upon reaching predefined humane-endpoint criteria.

#### Animal cohorts and hydrogel implantation

Unless otherwise stated, non-tumor hydrogel studies were conducted in 6-week-old female BALB/cJ mice. C57BL/6J mice were used for all B16F10 tumor studies. C57BL/6J mice were additionally included in the day-14 non-tumor hydrogel study to compare hydrogel responses across strains before subsequent tumor studies. The C57BL/6J day-14 cohort was subsequently used as the non-tumor reference for the day-14 versus day-42 flow-cytometry comparison with C57BL/6J hydrogel-derived tissues collected from the tumor study.

#### Hydrogel immune-profiling and immunization studies

Six-week-old female BALB/cJ mice received two 50 μL subcutaneous flank implants of OCL hydrogels containing 35 μg ovalbumin (OVA; InvivoGen, Cat# vac-pova), 0.4 μg CCL21 (BioLegend, Cat# 586404), and 0.4 μg LIGHT (BioLegend, Cat# 557604) per hydrogel, with one implant placed on each flank. For day-7 and day-21 analyses, cohorts were staggered such that all mice were 6 weeks old at the time of implantation and day-7 and day-21 tissues were harvested and processed concurrently. Soft OCL hydrogel-derived tissues were collected at day 7, whereas soft and stiff OCL hydrogel-derived tissues were collected at day 21. Each experimental group contained five mice. At day 21, six of the ten implanted soft OCL hydrogels were identifiable and recoverable at explantation, yielding three explants for flow-cytometry analysis and three for histological and immunofluorescence analysis. Explants were assigned by flank for flow-cytometry or histological/immunofluorescence analysis, respectively. Six inguinal lymph nodes were additionally collected for flow-cytometry analysis, comprising three inguinal lymph nodes from soft OCL hydrogel-treated mice and three from stiff OCL hydrogel-treated mice. Mice assigned to the day-21 endpoint were bled weekly at weeks 1, 2 and 3 after implantation. Serum was isolated and stored frozen until analysis of OVA-specific IgG by ELISA.

The day-21 experiment additionally included mice receiving soft hydrogels containing OVA alone and were bled in parallel with the OCL groups at weeks 1, 2 and 3 after implantation. Because OVA-only hydrogels were not identifiable at the day-21 endpoint, this group was not included in hydrogel-derived tissue analyses. Serum from these animals was retained for comparison of OVA-specific IgG responses between the OVA-only and OCL groups.

Day-14 samples were obtained from a separate experiment in which 6-week-old female BALB/cJ and C57BL/6J mice received two 50 μL subcutaneous flank implants of OCL hydrogels, with one implant placed on each flank. Each experimental group contained five mice. Hydrogel-derived tissues were collected 14 days after implantation. For each mouse, the explant from one flank was processed for flow-cytometry analysis, whereas the contralateral explant was processed for histological and immunofluorescence analyses. For temporal comparisons of soft OCL hydrogels across days 7, 14, and 21, only BALB/cJ samples were included.

In a separate day-7 experiment, 6-week-old female BALB/cJ mice were used to evaluate the early effects of hydrogel mechanics and OCL cargo on immune-cell recruitment. Mice received two 50 μL flank implants of the same assigned formulation: soft empty, soft OCL, stiff empty or stiff OCL hydrogels. Each group contained three mice, yielding six explants per condition. Five explants per condition were analyzed individually by flow cytometry. The remaining explant from each of the four conditions was combined and used as a pooled control sample for flow-cytometry staining and gating.

#### Hydrogel persistence and remodeling studies

To evaluate hydrogel persistence in vivo, four 6-week-old female BALB/cJ mice received paired 50 μL subcutaneous flank implants of empty soft and stiff hydrogels containing Cy5-labeled liposomes, with the soft hydrogel implanted in the left flank and the stiff hydrogel implanted in the right flank. Mice were imaged longitudinally at the indicated timepoints over 21 days using an IVIS Spectrum Optical Imaging System at the Oncology Precision Therapeutics and Imaging Core (OPTIC), Herbert Irving Comprehensive Cancer Center, Columbia University. Fluorescence images were acquired using 640-nm excitation and 680-nm emission filters with auto-exposure mode and analyzed using Living Image software. Regions of interest of identical size were positioned over each implant site and maintained across animals, formulations, and imaging timepoints. Background total radiant efficiency was subtracted from each measurement, and the background-corrected signal for each implant was normalized to its corresponding value at the initial imaging timepoint (t_0_). Each implant time course was fitted independently in GraphPad Prism using a one-phase exponential decay model, and signal half-life was calculated from the fitted decay constant. Individual normalized decay profiles and corresponding fits are shown in Supplementary Fig. 3B–C. At day 21, mice were euthanized and implant sites were examined macroscopically for residual hydrogel-derived material.

For analysis of local hydrogel remodeling, separate cohorts of 6-week-old female BALB/cJ mice received soft or stiff OCL hydrogels prepared using liposomes containing PE-Cy5. Explants were collected at days 7, 14, and 21 after implantation, with three hydrogel explants collected per formulation at each timepoint. Explants were fixed overnight in 10% formaldehyde and cryoprotected by incubation in 15% sucrose for 24 h followed by 30% sucrose for 24 h. Samples were embedded in OCT compound, cryosectioned, processed for hematoxylin and eosin or Masson’s trichrome staining, or imaged by confocal microscopy to detect retained Cy5 signal. To preserve hydrogel integrity during direct fluorescence imaging, sections were imaged immediately after drying and mounting without aqueous washing steps, thereby preventing detachment of hydrated hydrogel cores during processing. Explants were sectioned through the center of the hydrogel implant. Cy5-positive regions were segmented in Fiji (ImageJ)^67^, and the retained hydrogel fraction was calculated as the Cy5-positive area normalized to the total explant cross-sectional area.

#### Hydrogel-versus-bolus immunization study

In a separate immunization experiment, 6-week-old female BALB/cJ mice received two subcutaneous OCL administrations, one on each flank, either incorporated within 50 μL soft hydrogels or delivered as a soluble 50 μL bolus in PBS using matched cargo doses. Blood was collected at the indicated timepoints, and serum was isolated and stored frozen until analysis of OVA-specific IgG by ELISA.

### Animal tumor studies

#### B16F10-OVA melanoma cells

B16F10 melanoma cells expressing the OVA-derived SIINFEKL epitope fused to GFP (B16F10-OVA) were generously obtained from the laboratory of Nicholas Arpaia (Columbia University), where they had originally been provided by Andrea Schietinger (MSKCC), as previously described^68^. The B16F10-OVA cell line was authenticated by mouse short tandem repeat (STR) profiling (CD Genomics), which showed a 96.55% match to the B16-F10 reference profile (CVCL_0159) and no evidence of human-cell cross-contamination. Cells were maintained in Gibco DMEM, high glucose, pyruvate supplemented with 10% fetal bovine serum (FBS) and 1% penicillin-streptomycin at 37 °C and 5% CO₂. Cells were routinely passaged twice weekly.

#### Soft-versus-stiff prophylactic study

Six-week-old female C57BL/6J mice received a single 50 μL subcutaneous flank implantation of a soft or stiff empty hydrogel or a soft or stiff OCL hydrogel. OCL hydrogels contained 35 μg OVA, 0.4 μg CCL21, and 0.4 μg LIGHT. Fourteen days after hydrogel implantation, mice were challenged subcutaneously with 100,000 B16F10-OVA melanoma cells. Anti-PD-L1 (BioXCell, Cat# BE0101-R025mg) antibody was administered intraperitoneally at 200 μg per dose twice weekly beginning 14 days after tumor inoculation and continuing until the experimental or humane endpoint. Tumors were measured every other day using calipers, and tumor area was calculated as the area of an ellipse based on width and length of tumor. Mice were euthanized when tumor diameter approached 20 mm along one axis or earlier when investigator assessment indicated that humane-endpoint criteria would be reached before the next scheduled monitoring timepoint.

#### Soft-hydrogel cargo-composition study

Six-week-old female C57BL/6J mice received 50 μL subcutaneous implants of soft hydrogels containing no cargo, OVA alone, OVA plus MPLA, or OCL cargo. Hydrogels were implanted 14 days before subcutaneous challenge with 100,000 B16F10-OVA cells. Blood was collected 21 days after tumor inoculation, and mice were euthanized for endpoint analyses on day 28 after tumor inoculation or earlier upon reaching humane-endpoint criteria. Tumor growth was monitored as described above. At endpoint, spleens and identifiable implant-derived tissues were collected for flow-cytometry analysis.

### Histological and immunofluorescence analysis

#### Immunofluorescence staining of immune niches

Explanted hydrogels were embedded in Tissue-Tek O.C.T. compound (Sakura, Cat# 4583) within 15 x 15 x 5 mm disposable molds (Fisher Scientific, Cat# 22-363-553) and cryopreserved by rapid freezing in a precooled (-80°C) aluminum can of isopentane, within a liquid-nitrogen bath. Samples were stored at −80 °C until cryosectioning. Frozen specimens were sectioned at 12 μm and mounted on glass slides. Slides were brought to room temperature for 30 min, and hydrophobic barriers were drawn around samples using a PAP pen. Sections were fixed in 4% paraformaldehyde for 20 min at room temperature and washed twice with PBS-Tween. Fc-receptor blocking was performed using BlockAid (Invitrogen, Cat# B10710) for 2 h at room temperature or overnight at 4 °C in a humidified chamber.

For soft-versus-stiff imaging, antibodies included B220-BV421, clone RA3-6B2 (BioLegend, Cat# 103239); CD3-AF647, clone 17A2 (BioLegend, Cat# 100209); and CD11c-AF594, clone N418 (BioLegend, Cat# 117356). For temporal imaging of soft hydrogels, antibodies were applied in separate iterative staining rounds as described below and included B220-AF647, clone RA3-6B2 (BioLegend, Cat# 103229); CD3-AF647, clone 17A2 (BioLegend, Cat# 100209); CD11c-AF594, clone N418 (BioLegend, Cat# 117356); PNAd, clone MECA-79 (BioLegend, Cat# 120803), followed by a DyLight-conjugated streptavidin secondary antibody (BioLegend, Cat# 405129); and CD31-AF647, clone 390 (BioLegend, Cat# 102415). For PNAd staining, an additional streptavidin/biotin blocking step was performed using a commercial blocking kit (Thermo Scientific, Cat# R37628) before primary-antibody incubation.

Primary antibodies were diluted in 2% FBS in PBS and applied for 2 h at room temperature or overnight at 4 °C in a dark, humidified chamber. Slides were washed with PBS-Tween and incubated with secondary detection reagents diluted in 5% FBS in PBS for 1 h at room temperature in the dark. After washing, slides were briefly rinsed in PBS and mounted using ProLong Gold antifade reagent with or without DAPI (Invitrogen, Cat# P10144 or P36941), sealed with glass coverslips, cured overnight at room temperature in the dark, and stored at 4 °C until imaging.

For histological analysis, 12-μm cryosections were sent to the Molecular Pathology Core at Columbia University Irving Medical Center for hematoxylin and eosin staining using standard protocols.

#### Iterative immunofluorescence imaging

For iterative bleaching and staining, sections mounted using Fluoromount-G mounting medium (SouthernBiotech, Cat# 0100-01) were incubated in PBS for 2 h at room temperature or overnight at 4 °C to facilitate coverslip removal. Slides were washed twice with PBS-Tween and incubated for 10 min at room temperature under ambient light with freshly prepared 1 mg mL−1 lithium borohydride in deionized water (Strem Chemicals, Cat# 393-0397), filtered through a 0.22-μm nylon membrane. Sections were then washed three times with PBS-Tween before subsequent staining cycles. Images from successive cycles were registered in Fiji using the TurboReg plugin^69^ in accurate manual mode. DAPI was included in each cycle and used as the common reference channel for image registration and cellular localization.

#### Confocal imaging

Confocal images were acquired using a Nikon AXR resonant-scanning confocal microscope mounted on an Ti2 inverted microscope with a 20x/0.75 PlanApo violet-corrected objective lens (Nikon, Melville, NY). DAPI and BV421 were excited with a 405 nm laser and emission was collected from 425-475 nm. DyLight was excited with a 488 nm laser and emission was collected from 500-550 nm. AF594 was excited with a 561 nm laser and emission was collected from 570-625 nm. AF647 was excited with a 647 nm laser and emission was collected from 650-720 nm. Images were acquired using gallium arsenide phosphide (GaAsP) (green and red fluorophores) and conventional silicon (blue and far-red fluorophores) PMTs, sequentially using line-series acquisition with minimal crosstalk, resonant bi-directional scanning, 2x line averaging, a pixel size of approximately 0.49 μm and a scan zoom of 1.25x. Z stacks were collected at intervals of 3 µm over a total depth of 12 μm. Maximum intensity projections were generated, and adjacent fields of view were stitched into large images using the Optimal Path stitching method. Stitched images were subsequently denoised using the Denoise.ai function in NIS-Elements software. The microscope was controlled by NIS Elements software version 6 (Nikon).

#### B-cell aggregate image analysis

B-cell aggregate organization was quantified from confocal images using Fiji. All images were processed using the same Denoise.ai workflow in NIS-Elements software prior to quantitative analysis via Fiji. The B220 channel was separated from the remaining fluorescence channels and processed using a Gaussian blur with σ = 1. B220⁺ signal was segmented using Otsu automatic thresholding, followed by binary opening to remove isolated background pixels.

The entire local cellular aggregate containing the B-cell-rich region was manually outlined to define the aggregate cross-sectional area. B-cell occupancy was defined as the total thresholded B220⁺ area within the manually outlined aggregate. Aggregate compactness was calculated as:

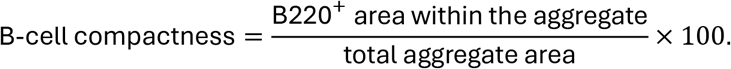

All identifiable B-cell-rich aggregates within each analyzed hydrogel section were quantified. Three biologically independent hydrogels were analyzed at each timepoint. Where multiple aggregates were identified within a section, measurements were averaged to obtain a single hydrogel-level value.

#### HGE nuclear-density quantification

Nuclear density within hydrogel-derived tissue explants was quantified from hematoxylin and eosin (HCE)-stained cross sections using ǪuPath, an open-source platform for digital pathology image analysis^70^. For each section, the hydrogel-associated tissue region was manually annotated to exclude background and surrounding non-implant tissue, including adjacent muscle and skin. Automated nuclear detections were then performed within the annotated region based on hematoxylin nuclear staining. Nuclear density was calculated as the number of detected nuclei normalized to the annotated tissue area and reported as nuclei per mm².

Images were analyzed at a fixed resolution of 0.5 μm pixel⁻¹. Local background subtraction was performed using an 8-μm radius, and morphological opening by reconstruction was applied to reduce uneven background staining. No median filtering was applied, and Gaussian smoothing was performed with σ = 1.5 μm. Detected objects with nuclear areas below 10 μm² or above 400 μm² were excluded to remove small staining artifacts and large non-nuclear structures, respectively. Nuclei were identified using a hematoxylin optical-density threshold of 0.1, with a maximum background intensity of 2. Shape-based splitting was enabled to separate adjacent nuclei.

### Flow cytometry

#### Tissue dissociation

Hydrogel-derived tissues and lymph nodes were mechanically dissociated using Kimble BioMasher II tissue grinders (Cat# 749625-0020) and enzymatically digested in DMEM containing 100 μg mL⁻¹ DNase I (Sigma Aldrich, Cat# 10104159901) and 1 mg mL⁻¹ Collagenase/Dispase (Sigma Aldrich, Cat# 10269638001). Samples were incubated for 45 min at 37 °C with constant agitation on a temperature-controlled shaker. Cell suspensions were passed through 35-μm strainers into 5-mL polystyrene round-bottom tubes (Falcon, Cat# 352235) and centrifuged at 400 × g for 10 min at 4 °C. Spleens were processed separately by mechanical dissociation without enzymatic digestion. Splenic tissue was dissociated through sterile 70-μm strainers (Thermo Fisher Scientific, Cat# 229483) using the back of a syringe plunger. Cell suspensions were centrifuged at 400 × g for 10 min at 4 °C, resuspended in 3 mL ACK red-blood-cell lysis buffer per spleen, and incubated for 10 min at room temperature. Samples were then centrifuged at 400 × g for 5 min. Cell pellets were transferred to U-bottom 96-well plates for staining.

#### Cell staining and acquisition

For immune-cell profiling of hydrogel-derived tissues and lymph nodes, cells were stained in 100 μL PBS containing Zombie NIR viability dye at a 1:1,000 dilution (BioLegend, Cat# 423105) for 20 min at room temperature. Cells were washed sequentially with 100 and 200 μL FACS buffer consisting of PBS containing 2% FBS. Fc receptors were blocked using 50 μL TrueStain FcX diluted 1:100 in FACS buffer for 20 min at 4 °C. An equal volume of antibody cocktail was then added. The cocktail contained anti-CD45–BV510 (clone 30-F11, BioLegend, Cat# 103137), anti-CD3–PE (clone 17A2, BioLegend, Cat# 100205), anti-CD4–Alexa Fluor 700 (clone GK1.5, BioLegend, Cat# 100429), anti-CD8–Spark PLUS UV395 (clone 53-6.7, BioLegend, Cat# 104011), anti-CD19–BV421 (clone 6D5, BioLegend, Cat# 115537), anti-CD95–BV605 (clone SA367H8, BioLegend, Cat# 152612), anti-GL7–PE-Cy7 (clone GL7, BioLegend, Cat# 144619), anti-CD44–PerCP-Cy5.5 (clone IM7, BioLegend, Cat# 103031), anti-CD62L–BV785 (clone MEL-14, BioLegend, Cat# 104440), anti-CD11c–APC (clone N418, BioLegend, Cat# 117309), anti-CD11b–Alexa Fluor 488 (clone M1/70, BioLegend, Cat# 101219), anti-F4/80–RB705 (BD Biosciences, Cat# 570289), and anti-Ly6G–APC-Cy7 (clone 1A8, BioLegend, Cat# 127623). Individual antibodies were prepared at 1:100 in the cocktail, yielding a final dilution of 1:200 after addition to the Fc-blocking solution.

For splenic T-cell profiling in the soft-hydrogel cargo-composition tumor study, a separate staining panel was used containing Zombie Aqua viability dye at 1:1,000 dilution (BioLegend, Cat# 423101), anti-CD3–PE-Fire 700 (clone 17A2, BioLegend, Cat# 100272), anti-CD4–Spark PLUS UV395 (clone GK1.5, BioLegend, Cat# 100496), anti-CD8–APC-Fire 750 (clone 53-6.7, BioLegend, Cat# 100766), anti-CD25–PE (clone 3C7, BioLegend, Cat# 101904), anti-CD69–PE-Cy7 (clone H1.2F3, BioLegend, Cat# 104512), anti-PD-1–BUV615 (clone 29F.1A12, BD Biosciences, Cat# 568601), and anti-LAG-3–Alexa Fluor 647 (clone C9B7W, BioLegend, Cat# 125242).

For both panels, cells were stained with the corresponding antibody cocktail for 30 min at 4 °C, washed, and fixed in 4% paraformaldehyde for 10 min at room temperature. Samples were washed twice, resuspended in FACS buffer, and acquired the following day using a Sony ID7000 spectral flow cytometer.

#### Flow-cytometry UMAP analysis

Dimensionality reduction of flow-cytometry data was performed in FlowJo using the UMAP_R plugin. Following exclusion of debris, doublets, and nonviable cells, CD45⁺ leukocytes were embedded using expression of CD45, CD11b, CD3, CD4, CD8, CD19, Ly6G, F4/80, and CD11c. UMAP was performed using cosine distance, 30 nearest neighbors, a minimum distance of 0.1, and two output dimensions. Immune-cell populations were subsequently identified using conventional manual gating based on marker expression, and the corresponding gated populations were overlaid onto the UMAP embedding to visualize their distribution across the phenotypic landscape.

#### Flow-cytometry analysis

Debris, doublets, and nonviable cells were excluded before identification of CD45⁺ leukocytes. For hydrogel-derived tissues, the entire processed cell suspension from each explant was acquired, allowing total recovered cell numbers and absolute immune-cell counts to be determined from the acquired events. Immune populations were identified using a sequential gating strategy. Macrophages were first identified as CD11b⁺F4/80⁺ cells. Within the remaining F4/80⁻ population, neutrophils were identified as CD11b⁺Ly6G⁺ cells. After macrophages and neutrophils were excluded, CD11b⁻ cells were selected, and T and B cells were identified as CD3⁺CD19⁻ and CD3⁻CD19⁺ cells, respectively. Dendritic cells were identified within the macrophage- and neutrophil-depleted population by excluding CD3⁺ and CD19⁺ lymphocytes and subsequently gating CD11c⁺ cells. Naïve, effector-memory, and central-memory T cells were defined as CD44⁻CD62L⁺, CD44⁺CD62L⁻, and CD44⁺CD62L⁺ cells within the CD3⁺ T-cell population, respectively. GC-like B cells were defined as CD95⁺GL7⁺ within CD19⁺ B cells. CD11c⁺ B cells were defined as CD11c⁺ within CD19⁺ B cells.

For splenic T-cell analysis, following the exclusion of debris, doublets, and nonviable cells, CD3⁺ T cells were separated into CD4⁺ and CD8⁺ populations. CD4:CD8 ratios were calculated from the relative frequencies of these populations. CD25 and CD69 expression was quantified within the CD8⁺ T-cell compartment. PD-1 and LAG-3 expression was assessed within the CD4⁺ and CD8⁺ T-cell compartments, and PD-1⁺LAG-3⁺ cells were quantified as a co-expressing population.

### OVA-specific serum IgG ELISA

OVA-specific serum antibody responses were quantified by enzyme-linked immunosorbent assay (ELISA). Standard high-binding 96-well plates (Nunc-Immuno MicroWell, Thermo Scientific, Cat# 439454) or high-binding half-area 96-well plates (MICROLON 600, Fisher Scientific, Cat# 07-000-720) were coated with OVA at 5 μg mL⁻¹ in PBS and incubated overnight at 4 °C. Coating volumes were 50 μL per well for standard plates and 25 μL per well for half-area plates. Plates were washed twice with PBS containing 0.05% Tween-20 (PBS-T; MilliporeSigma, Cat# 524653) and blocked with 1% bovine serum albumin (BSA) in PBS for 2 h at room temperature or overnight at 4 °C. Serum samples were initially diluted 1:25 or 1:50 and subsequently serially diluted threefold in non-treated 96-well plates. Following blocking, ELISA plates were washed twice with PBS-T, serum dilutions were transferred to the OVA-coated plates, and samples were incubated for 2 h at room temperature. Plates were then washed three times with PBS-T.

OVA-specific total IgG and IgG subclass responses were detected using horseradish peroxidase (HRP)-conjugated secondary antibodies diluted 1:5,000 in 1% BSA and incubated for 1 h at room temperature. Total IgG was detected using AffiniPure goat anti-mouse IgG (H+L)-HRP (Jackson ImmunoResearch, Cat# 115-035-003). IgG1 responses were detected using goat anti-mouse IgG1-HRP (SouthernBiotech, Cat# 1071-05). IgG2a responses were measured for BALB/c serum samples using goat anti-mouse IgG2a-HRP (SouthernBiotech, Cat# 1081-05), whereas IgG2c responses were measured for C57BL/6J serum samples using goat anti-mouse IgG2c-HRP (SouthernBiotech, Cat# 1078-05). Following secondary-antibody incubation, plates were washed four times with PBS-T and developed in the dark using Pierce 1-Step Ultra TMB-ELISA Substrate Solution (Thermo Scientific, Cat# PI34029). Reactions were stopped with 1 N sulfuric acid (Fisher Chemical, Cat# SA2121-1). Absorbance was measured at 450 and 570 nm using a microplate reader, and background-corrected absorbance was calculated as A₄₅₀ − A₅₇₀. Antibody titers were determined by fitting background-corrected absorbance values across the serum dilution series using a four-parameter logistic (4PL) nonlinear regression model in GraphPad Prism. The reciprocal serum dilution corresponding to A₄₅₀ − A₅₇₀ = 0.5 was interpolated from the fitted curve and reported as OVA-specific titer.

### Single-cell RNA sequencing

#### Tissue dissociation for single-cell RNA sequencing

Hydrogel explants were mechanically and enzymatically dissociated using the same BioMasher, DNase I, and Collagenase/Dispase procedure described above. Cell suspensions were passed through 35-μm strainers and centrifuged at 400 × g for 10 min at 4 °C. Before cell counting, residual erythrocytes were lysed for 5 min at room temperature using blood-cell lysis buffer. Samples were centrifuged at 400 × g for 5 min, and cell pellets were resuspended for cell counting and loading.

#### Single-cell cDNA-library construction and sequencing

Single-cell RNA sequencing (scRNA-seq) was performed across three independent experiments using the 10x Genomics Chromium Single Cell 3′ platform. Cells were collected from pools of soft or stiff gels, with the number of gels per pool varying across conditions and experiments to achieve sufficient cell concentrations (Supplementary Table 3). For all samples except day 14 soft gel pools, viable cell concentrations were determined using an automated cell counter, and cell suspensions were loaded onto a Chromium X instrument using either GEM-X OCM 3′ Chips (10x Genomics; PN-1000747) or Chromium GEM-X Single Cell 3′ Chips v4 (10x Genomics; PN-1000690), following the manufacturer’s protocols and targeting the cell recoveries specified in Supplementary Table 3. Cell suspensions from day 14 soft gel pools were too dilute for reliable counting; therefore, the maximum allowable suspension volume was loaded per lane. Libraries were prepared according to the Chromium GEM-X Single Cell 3′ Reagent Kits v4 user guide (10x Genomics; CG000731 Rev B) or the GEM-X Universal 3′ Gene Expression v4 4-plex user guide (10x Genomics; CG000768 Rev B). Fragment size distributions of amplified cDNA and final sequencing libraries were assessed using a 2200 TapeStation system (Agilent) with D5000 reagents. Final libraries were normalized to 4 nM and pooled for sequencing by Novogene using its standard workflow on an Illumina NovaSeq X Plus platform, with a paired-end 150 bp read configuration and a target depth of ≥ 20,000 read pairs per cell.

#### Single-cell RNA-sequencing processing, clustering, and annotation

Raw base call files were processed using the 10x Genomics Cell Ranger Gene Expression pipeline v8.0.1 or v10.0.0. Reads were aligned to a mouse reference genome, and gene-by-cell count matrices were generated for each sample. The SoupX^71^ R package was used to remove ambient mRNA from each count matrix separately. For each sample, a cell_data_set (CDS) object was created using the R package monocle3^72–74^ from the count matrix, gene metadata, and cell annotations. CDS objects were then combined, and cells with fewer than 500 unique molecular identifiers (UMIs) or more than 20% mitochondrial content were filtered out. Doublets were identified using the Single-Cell Remover of Doublets (Scrublet^75^) framework in Python, and cells with Scrublet scores greater than 0.2 were excluded. The filtered dataset was normalized by size factors, preprocessed by principal component analysis using the top 35 principal components and top highly variable genes, and aligned using mutual nearest neighbor correction^76^ to reduce experiment-specific batch effects. The aligned PCA space was clustered using Leiden community detection^77,78^ and visualized using Uniform Manifold Approximation and Projection^45^. Clustering resolution was selected using silhouette score analysis with the silhouette() function from the cluster R package^79^. Cluster-specific marker genes were identified using the top_markers() function from the monocle3 package and evaluated together with canonical cell-type marker expression to assign cell-type identities to each cluster. For phenotypic characterization of cell populations and downstream analyses, specific clusters were subjected to a second round of preprocessing, clustering, and annotation using the same parameters as specified above.

#### Gene-expression analysis

Expression of selected genes of interest was assessed across cell types and experimental conditions. Size factor-normalized expression values were extracted from the monocle3 CDS object, log_10_-transformed following addition of a pseudocount (+1), and averaged across cells within each cell type and condition group. Mean expression values were visualized as heatmaps using the ggplot2 R package, with each panel representing an individual gene.

#### Gene-signature scoring

Functional gene signatures related to cell motility, lymphoid organization, TLS-induction-associated cytokines, lymphoid migration, germinal center function, and antigen presentation were assembled from genes selected based on their reported association with these biological processes (Supplementary Table 4). A cell proliferation signature was obtained from a curated list of cell cycle genes^80^. A lymphoid homing signature was derived from the cytokine-induced immune programs described in the Human Cytokine Dictionary^55^. For each cell, aggregate expression scores for each signature were calculated using the calculate_aggregate_expression_score function, where genes for each signature are size-factor normalized, aggregated, and log-transformed to generate a signature score. Signature scores were then joined with the cell metadata and compared between selected conditions within each annotated cell type. For this, signature aggregate scores were fit to a Gaussian regression model using the speedglm^81^ function, specifying condition and experiment as covariates. Multiple hypothesis correction was performed using the Benjamini-Hochberg (BH) method. Significant condition-specific signatures were defined as those with BH-adjusted *P* value < 0.1 and model coefficient (*β*-coefficient) > 0.1 and visualized as dot plots or heatmaps of model coefficients, with non-significant values set to zero. For the temporal comparison of soft hydrogels between days 7 and 14, coefficients with an absolute *β* > 0.25 were displayed irrespective of adjusted *P* value, with statistical significance indicated by the corresponding Benjamini–Hochberg-adjusted *q* values.

#### Differential gene-expression analysis

##### Condition-specific differential expression

Condition-specific DEGs were calculated by fitting gene expression to a quasi-Poisson regression model of a gene as a function of condition and experiment using the monocle3 R package fit_models function. Genes expressed in fewer than 1% of cells were excluded. *P* values were FDR corrected using Benjamini-Hochberg correction for multiple hypotheses, and significant DEGs for a condition were defined as genes with BH-adjusted P value < 0.05 and an absolute normalized effect of ≥ 0.05. DEGs were visualized using volcano plots. The top 40 significant DEGs, ranked by absolute normalized effect, were additionally visualized using a heatmap. For each selected gene, log-normalized expression was averaged across cells within each gel-pool sample and the resulting sample-level means were standardized to Z-scores across samples.

Heatmaps were generated using the Heatmap() function from the ComplexHeatmap^82^ R package, with genes as rows and gel-pool samples as columns. Z-scores were capped at ±2 for display.

##### Cluster-specific differential expression for cytokine-induced cell polarization

Cluster-specific DEGs were calculated by fitting gene expression to a quasi-Poisson regression model of a gene as a function of Cluster using the monocle3 R package fit_models function where each cluster was compared to all other clusters, and condition was not specified as a covariate. Genes expressed in less than 1% of cells were excluded from the test. *P* values were FDR corrected using Benjamini-Hochberg correction for multiple hypotheses. Significant DEGs for a cluster were defined as genes with BH-adjusted *P* value < 0.05 and an absolute normalized effect of ≥ 0.05. DEGs were used for downstream analyses of cytokine-induced cell polarization.

##### Cluster-specific differential expression for CIP enrichment analysis

Cells were randomly subsampled to a maximum of 2,000 per cluster–condition–gel-pool combination using a fixed random seed. Genes detected in at least 5% of cells in the subsampled dataset were retained for analysis. Cluster-specific DEGs were calculated by fitting gene expression to a quasi-Poisson regression model of a gene as a function of Cluster using the monocle3 R package fit_models function where each cluster was compared to all other clusters and condition was not specified as a covariate. *P* values were FDR corrected using Benjamini-Hochberg correction for multiple hypotheses. Significant DEGs for a cluster were defined as genes with BH-adjusted *P* value < 0.05 and an absolute normalized effect of ≥ 0.25. DEGs were used for downstream program enrichment analyses.

##### Gene-set enrichment analysis

GSEA was performed using the R package clusterProfiler^83^ GSEA function using MSigDB^51,84^ C5 Gene Ontology biological process gene sets for Mus musculus, obtained through the msigdbr R package^85^. Differentially expressed genes were ranked by normalized effect size. For duplicated gene symbols, the entry with the largest absolute normalized effect was retained. GSEA was run with Benjamini-Hochberg correction, a minimum gene set size of 10, and a maximum gene set size of 500. Enriched pathways were summarized using normalized enrichment scores and adjusted *P* values. Gene-set names were simplified for visualization while retaining the underlying MSigDB gene-set definitions.

##### Cytokine-Induced Programs (CIPs) enrichment analysis

Myeloid cell-specific CIPs were obtained from the Human Cytokine Dictionary GitHub repository^55^. Each program was defined by its top 75 genes ranked by effect size. Human gene annotations were mapped to their mouse orthologs using the orthologs() function from the babelgene R package^86^. Unmapped genes were excluded, and duplicate mapped genes were removed within each program. For each cluster, DEGs with a BH-adjusted *P* value < 0.05 and a normalized effect > 0.25 formed the query gene set. The background comprised all genes tested for differential expression, corresponding to genes detected in at least 5% of cells in the subsampled dataset. Ǫuery and program gene sets were restricted to genes present in this background. Program over-representation was assessed using one-sided hypergeometric tests implemented with the R function phyper(). *P* values were adjusted using the Benjamini–Hochberg procedure across programs within each cluster, with an adjusted *P* value < 0.05 indicating significant enrichment. For visualization, significantly enriched programs were ranked by the fraction of program genes overlapping each cluster’s query gene set, and up to 10 programs per cluster were visualized using a dot plot. Program names were simplified for visualization while retaining the underlying gene-program definitions. Programs were grouped into broad functional categories and using all significantly enriched programs, the proportion belonging to each category was calculated within each cluster.

##### Cell–cell communication analysis

Differential ligand-receptor cell-cell communication across conditions was inferred using the R framework MultiNicheNet^58^ with the NicheNet-v2^87^ mouse ligand-receptor network and ligand-target prior knowledge model. Analyses were performed using the sample-agnostic MultiNicheNet workflow. The cell identity-annotated CDS object was converted to a Seurat object, log-normalized, and subsequently converted to a SingleCellExperiment object for downstream analysis. Communication analysis was performed across soft and stiff gels at day 7. Erythrocytes and epithelial cells were excluded from the sender and receiver cell-type sets. Cell type abundance and expression filtering were performed prior to communication analysis. Cell type-condition combinations were required to contain at least 10 cells, and genes were considered expressed if detected in at least 5% of cells. Ligand activity prediction was performed using the NicheNet ligand-target matrix, with differential expression thresholds of logFC > 0.25 and adjusted *P* value < 0.05. Ligand-receptor pairs were prioritized under the no_frac_LR_expr scenario. Prioritized ligand-receptor interactions for specific sender and receiver cell types were visualized using ligand-receptor product activity plots.

##### Immune Response Enrichment Analysis for cytokine-induced cell polarization

The top 100 cluster-specific DEGs, ranked by normalized effect size, were used to infer cytokine-induced polarization states for each cell subcluster. Enrichment analysis was performed using the Immune Dictionary’s Immune Response Enrichment Analysis software^50^ with a hypergeometric test.

### Statistical analysis

Statistical analyses of material characterization, imaging, flow-cytometry, serological, and tumor-study data were performed using GraphPad Prism. Unless otherwise indicated, data are presented as mean ± SEM. Comparisons between two groups were performed using unpaired two-tailed Student’s *t*-tests or Mann– Whitney *U* tests, as indicated. Comparisons among multiple groups were performed using one-way or two-way analysis of variance (ANOVA), followed by Tukey’s or Šídák’s multiple-comparisons tests as specified for the corresponding analysis. For the cargo-comparison tumor study and associated systemic immune analyses, one-way ANOVA was followed by the two-stage Benjamini, Krieger and Yekutieli procedure to control the false discovery rate, with adjusted values reported as *q* values. Survival curves were compared using the log-rank (Mantel–Cox) test. Statistical significance was defined as *p* < 0.05, or *q* < 0.05 for FDR-adjusted comparisons, unless otherwise specified. Exact *p* and *q* values are shown where indicated.

Nonlinear regression analyses were performed in GraphPad Prism as described for the corresponding assays. Longitudinal IVIS signal was fitted using a one-phase exponential decay model to determine signal half-life, whereas OVA-specific serum antibody titers were determined using four-parameter logistic regression of serum dilution curves. Statistical methods specific to single-cell RNA-sequencing analyses, including regression models, differential-expression testing, gene-set enrichment, and multiple-hypothesis correction, are described in the corresponding analysis subsections.

## Author contribution

R.R.H. and S.C. conceptualized the study and designed the experiments. R.R.H. developed the experimental methodology, led the investigation, performed formal analysis and data interpretation, prepared the visualizations, and coordinated the project. A.M. contributed to methodology and investigation. J.S.W. contributed to methodology and investigation. E.K. and C.W. contributed to investigation. M.K. contributed to flow-cytometry methodology and formal analysis. O.M.M. contributed to single-cell RNA-sequencing methodology, software, formal analysis, data curation, and visualization. J.L.M.-F. supervised and guided the single-cell RNA-seq profiling and analysis. O.M.M. and R.R.H. contributed to biological interpretation of the single-cell data. R.R.H. and S.C. wrote the original draft. A.M., O.M.M., and J.L.M.-F. contributed to review and editing of the manuscript. S.C. supervised the research and acquired funding. All authors reviewed and approved the final manuscript.

## Acknowledgments

We thank Dr. Robert Hincapie for guidance on protein labeling and purification, and the Columbia Stem Cell Initiative Flow Cytometry Core for technical support and advice. We thank Dr. Theresa Swane and Dr. Istvan Boldogh of the Confocal and Specialized Microscopy Shared Resource at the Herbert Irving Comprehensive Cancer Center (HICCC) for access to confocal microscopy instrumentation and assistance with immunofluorescence imaging. We also thank the Molecular Pathology Shared Resource at HICCC for histological staining and imaging, and the Oncology Precision Therapeutics and Imaging Core (OPTIC; HICCC) for access to IVIS instrumentation and guidance with image acquisition and analysis. The Confocal and Specialized Microscopy Shared Resource, Molecular Pathology Shared Resource, and OPTIC Shared Resource are supported in part by the NIH/NCI Cancer Center Support Grant P30CA013696. We thank Dr. Mathieu Rouanne, from the laboratory of Dr. Nicholas Arpaia, for providing the B16F10-OVA cell line.

## Declaration of generative AI and AI-assisted technologies in the writing process

During the preparation of this work, the authors used ChatGPT and Claude to assist with edits for grammar and clarity. The final text was reviewed and edited by the authors, who take full responsibility for the published content.

