## Supplemental data for "Supramolecular hydrogel viscoelasticity regulates in situ tertiary lymphoid neogenesis"

### Supplementary Information

#### Supplementary Figures

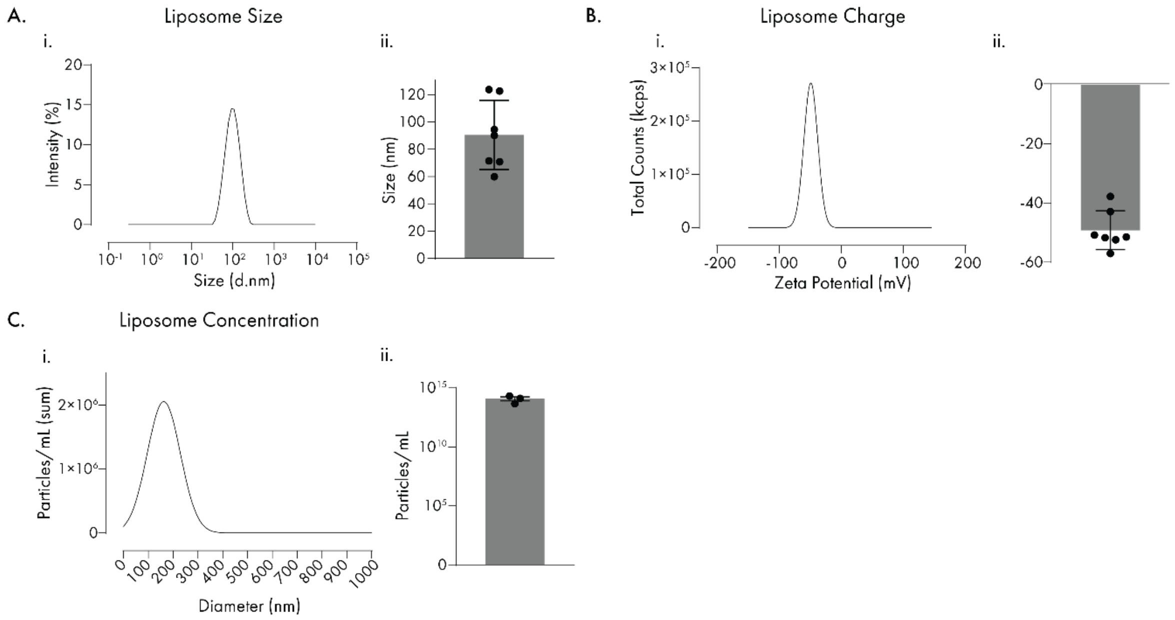

**Supplementary Figure 1. Characterization of liposomal hydrogel crosslinkers**. Liposomes were generated by either microfluidic synthesis using the ANP system or thin-film hydration followed by extrusion. (**A**) Average intensity-weighted particle-size distribution measured by dynamic light scattering and quantification of mean hydrodynamic diameter across independent liposome preparations. Each point represents the average of three technical measurements from one independently prepared liposome batch (n = 7 batches). (**B**) Average zeta-potential distribution and quantification of mean liposome surface charge. Each point represents the average of three technical measurements from one independently prepared batch (n = 7 batches). (**C**) Average particle-size distribution measured by nanoparticle tracking analysis and quantification of liposome particle concentration (n = 3 independent preparations). DLS size and zeta-potential measurements are presented as mean ± SD, and NTA concentration measurements are presented as mean ± SEM.

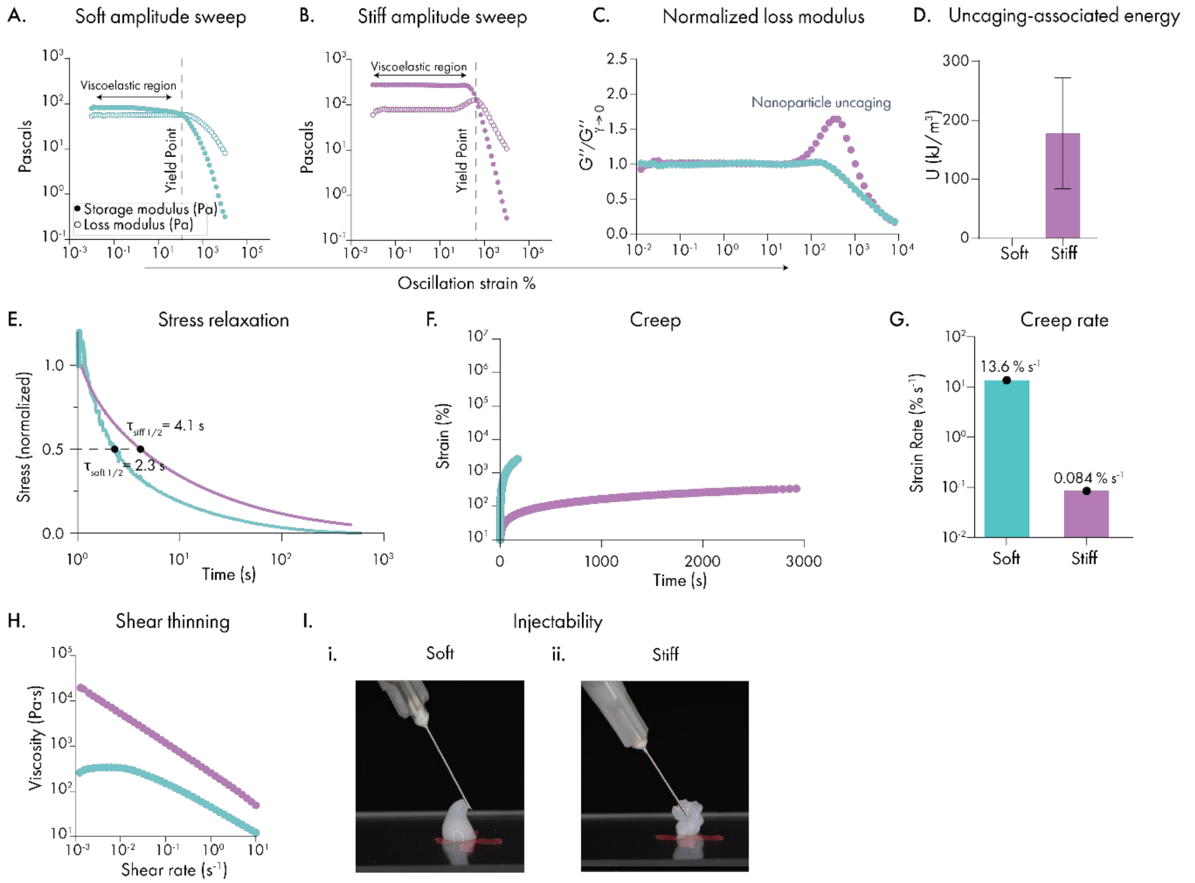

**Supplementary Figure 2. Nonlinear and time-dependent rheological behavior and injectability of soft and stiff supramolecular hydrogels.** (**A–B**) Representative oscillatory amplitude sweeps of soft and stiff hydrogels performed at 10 rad $s^{-1}$, showing storage modulus $G'$ and loss modulus $G''$ as functions of oscillatory strain. The linear viscoelastic region and yield point, defined by the crossover of $G'$and $G''$, are indicated. (**C**) Representative loss modulus normalized to its low-strain plateau value, $G''/G''_{\gamma\to 0}$, where $G''_{\gamma\to 0}$represents the loss modulus within the low-strain linear viscoelastic region. The stiff formulation exhibited a pronounced loss-modulus overshoot near yielding, associated with uncaging behavior, whereas the soft formulation underwent a more gradual decrease. (**D**) Quantification of the energy associated with the loss-modulus overshoot in stiff hydrogels, calculated from the fitted uncaging peak as described previously. No discrete uncaging-associated peak was detected in the soft formulation. Data are presented as mean ± SEM from three independent hydrogel preparations (n = 3). (**E**) Stress-relaxation profiles of soft and stiff hydrogels under a constant 10% strain. Stress was normalized to its initial value, and the stress-relaxation half-time ($\tau_1/_2$), defined as the time required for normalized stress to decrease to 0.5, was 2.3 s for the soft hydrogel and 4.1 s for the stiff hydrogel. (**F**) Creep response of soft and stiff hydrogels under a constant stress of 12.5 Pa, shown as strain as a function of time. (**G**) Steady-state creep strain rates of soft and stiff hydrogels, calculated from the terminal creep response. Stress-relaxation and creep measurements were performed once per formulation (n = 1). (**H**) Representative steady-shear flow sweeps showing shear-thinning behavior in soft and stiff hydrogels, with viscosity decreasing as a function of increasing shear rate. (**I**) Representative photographs of (i) soft and (ii) stiff hydrogels following extrusion through a 27G needle, demonstrating retention of a cohesive hydrogel following injection.

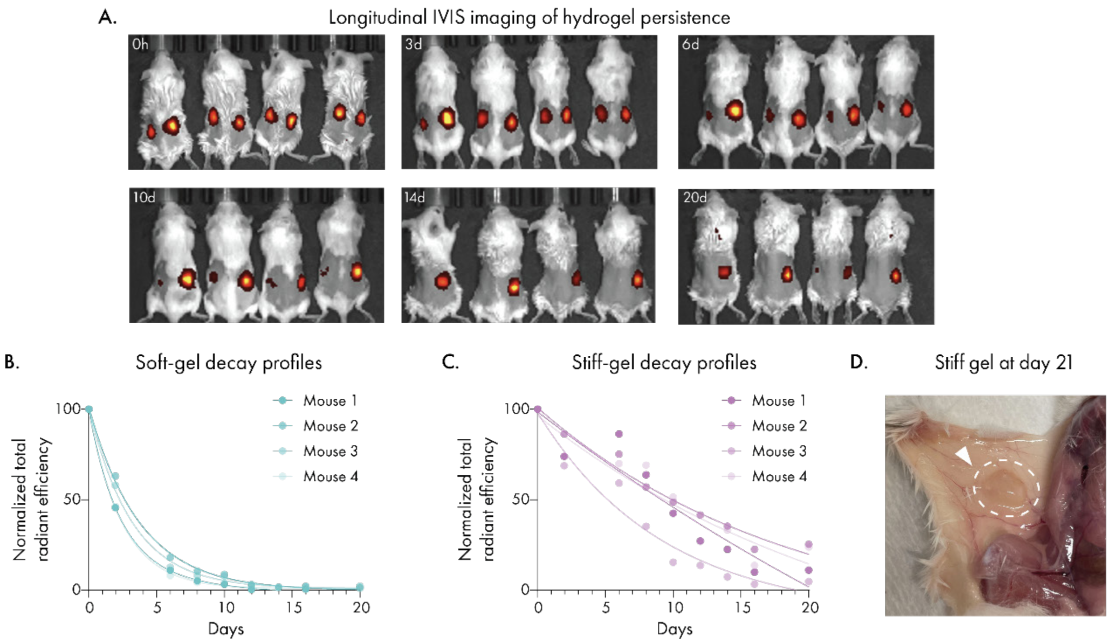

**Supplementary Figure 3. Longitudinal *in vivo* persistence profiles of empty soft and stiff hydrogels.** Six-week-old female BALB/c mice received paired subcutaneous injections of empty soft and stiff hydrogels containing Cy5-labeled liposomes, with soft and stiff hydrogels implanted in the left and right flanks, respectively (n = 4 mice). Liposomes were fluorescently labeled by incorporation of 18:0 Cyanine 5 PE into the liposomal formulation. (**A**) Representative longitudinal IVIS images acquired immediately after injection (0 h) and at days 3, 6, 10, 14, and 20. Cy5 fluorescence was measured using 640-nm excitation and 680-nm emission filters with automatic exposure. Images were generated using automatic display scaling and therefore are intended to show signal localization rather than permit direct visual comparison of fluorescence intensity across timepoints. (**B**-**C**) Individual normalized total radiant-efficiency profiles for soft and stiff hydrogels, respectively. Background fluorescence was subtracted, and each animal was normalized to its own signal at 0 h, defined as 100%. Regions of interest were maintained at a fixed size across animals and timepoints. Curves represent one-phase exponential-decay fits generated in GraphPad Prism. (**D**) Representative gross image of retained stiff hydrogel at day 21. The dashed circle and arrowhead indicate visible material. No corresponding soft hydrogel was grossly detectable at this timepoint.

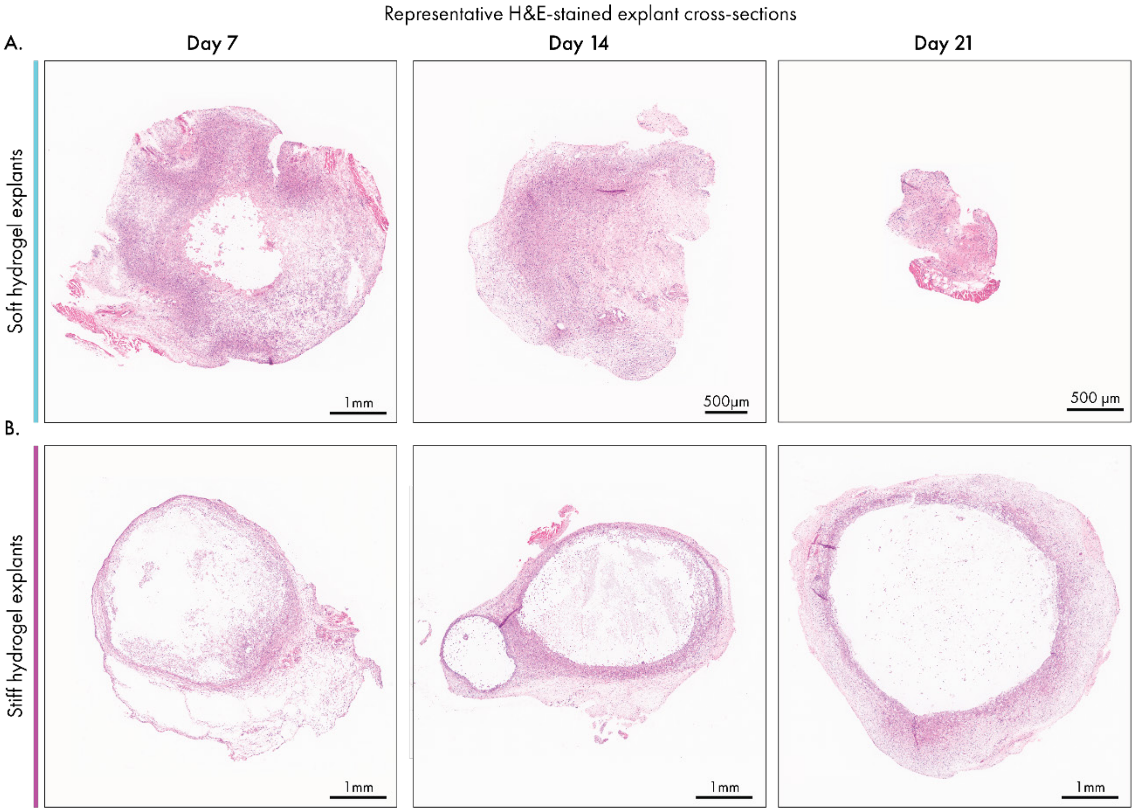

**Supplementary Figure 4. Soft and stiff hydrogels exhibit distinct spatial remodeling patterns over time.** Soft and stiff hydrogels containing ovalbumin (OVA), CCL21, and LIGHT were injected subcutaneously and retrieved at days 7, 14, and 21. Liposomes incorporated into the hydrogels were labeled with 18:0 Cyanine 5 PE. (**A**–**B**) Representative H&E-stained cross-sections of soft (A) and stiff (B) hydrogel explants retrieved at days 7, 14, and 21. Sections were collected at 12 μm thickness through the central plane of each explant. Images were resized by the indicated factors for visualization of differences in explant dimensions. Scale bars are indicated within each image.

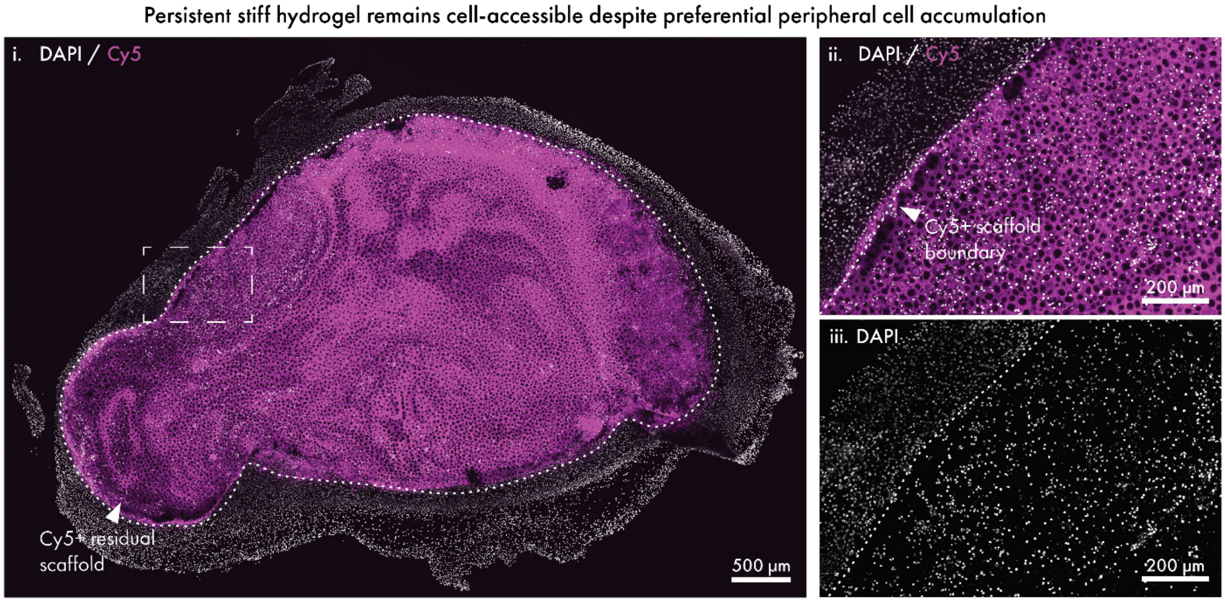

**Supplementary Figure 5. Cellular presence within the persistent stiff hydrogel scaffold.** Cy5 signal (magenta) delineates the residual hydrogel scaffold, and DAPI (gray) marks cell nuclei. The dotted line denotes the approximate boundary of the $Cy5^+$ residual scaffold. (**i**) Representative whole-explant image of a day-14 stiff hydrogel showing the persistent $Cy5^+$ scaffold. The boxed region is shown at higher magnification at right. (**ii**) Higher-magnification merged DAPI/Cy5 image showing $DAPI^+$ nuclei distributed within the $Cy5^+$ scaffold region and along the material interface. (**iii**) Corresponding DAPI-only image of the same region. Scale bars, 500 µm in the whole-explant image and 200 µm in higher-magnification images.

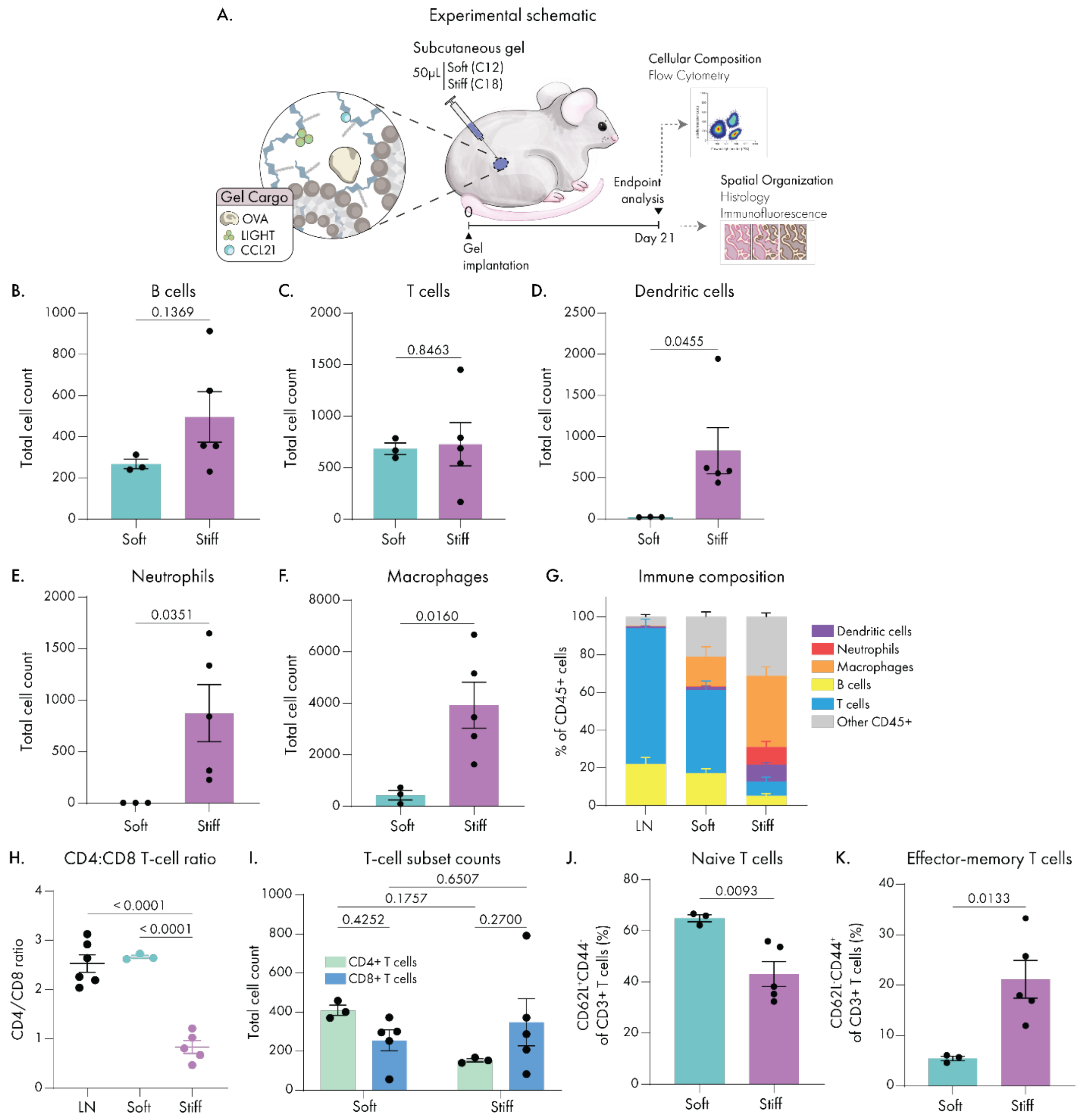

**Supplementary Figure 6. Day-21 immune composition and lymphocyte phenotypes in soft and stiff hydrogels.** Six-week-old female BALB/c mice received subcutaneous injections of either 50 μL soft (C12) or stiff (C18) hydrogels containing ovalbumin (OVA; 35 μg), CCL21 (0.4 μg), and LIGHT (0.4 μg). Hydrogel explants and inguinal lymph nodes were collected after 21 days for flow-cytometric and spatial analysis. (**A**) Experimental schematic illustrating hydrogel implantation and endpoint analysis by flow cytometry, histology, and immunofluorescence. (**B**-**F**) Total numbers of B cells (B), T cells (C), dendritic cells (D), neutrophils (E), and macrophages (F) recovered from soft and stiff hydrogel explants. These raw immune-cell counts underlie the lymphoid-to-myeloid ratio balance summarized in Figure 3F. (**G**) Immune composition of lymph nodes, soft hydrogels, and stiff hydrogels, expressed as the percentage of $CD45^+$ cells. Bars show the mean contribution of B cells, T cells, macrophages, neutrophils, dendritic cells, and other $CD45^+$ cells, with error bars representing SEM. This analysis provides the variability associated with mean immune-composition profiles shown in Figures 3G–I. (**H**) $CD4^+/CD8^+$ T-cell ratio in lymph nodes, soft hydrogels, and stiff hydrogels. (**I**) Total $CD4^+$ and $CD8^+$ T-cell counts recovered from soft and stiff hydrogel explants. (**J**–**K**) Frequencies of naïve $CD62L^+CD44^-$ T cells (J) and effector-memory $CD62L^-CD44^+$ T cells (K), expressed as percentages of $CD3^+$ T cells. Flow-cytometric analysis included three recovered soft-hydrogel explants (n = 3) and five stiff-hydrogel explants (n = 5) with lymph nodes collected across experimental conditions combined into a shared reference group (n = 6). Data are presented as mean ± SEM. Statistical significance in (B–F) and (J–K) was assessed using unpaired two-tailed Student’s t-tests; (H) was assessed using one-way ANOVA with Tukey’s multiple-comparisons test; and (I) was assessed using two-way ANOVA followed by Šídák’s multiple-comparisons test. Exact *p* values are shown.

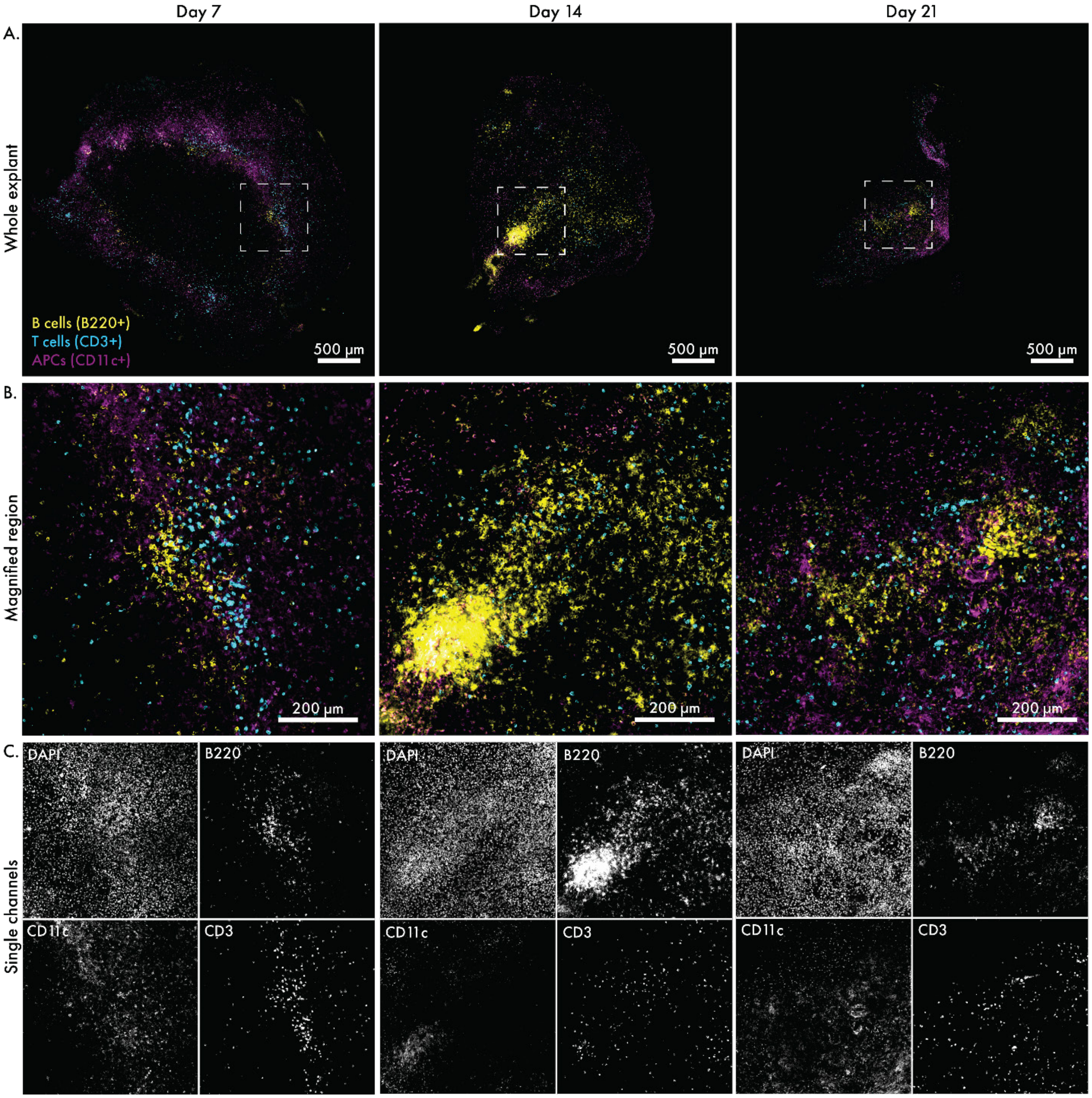

**Supplementary Figure 7. Whole-explant context and single-channel visualization of temporal lymphoid organization in soft hydrogels.** Soft hydrogels containing ovalbumin (OVA), CCL21, and LIGHT were implanted subcutaneously and retrieved at days 7, 14, and 21 for immunofluorescence imaging. Explants were sectioned through their central plane for spatial characterization. Explants were stained for B cells ($B220^{+}$, yellow), T cells ($CD3^{+}$, cyan), and $CD11c^{+}$ antigen-presenting cells (magenta), with DAPI marking nuclei. (**A**) Representative whole-explant overviews of soft-hydrogel explants at days 7, 14, and 21. Dashed boxes indicate the regions shown at higher magnification in (B). (**B**) Higher-magnification views of the boxed regions in (A), corresponding to the representative day-7, day-14, and day-21 images shown in Fig. 4A–C, respectively. (**C**) Corresponding single-channel images showing DAPI, B220, CD11c, and CD3 for each timepoint, displayed at reduced size relative to the corresponding merged images in (B). Scale bars, 500 µm in (A) and 200 µm in (B).

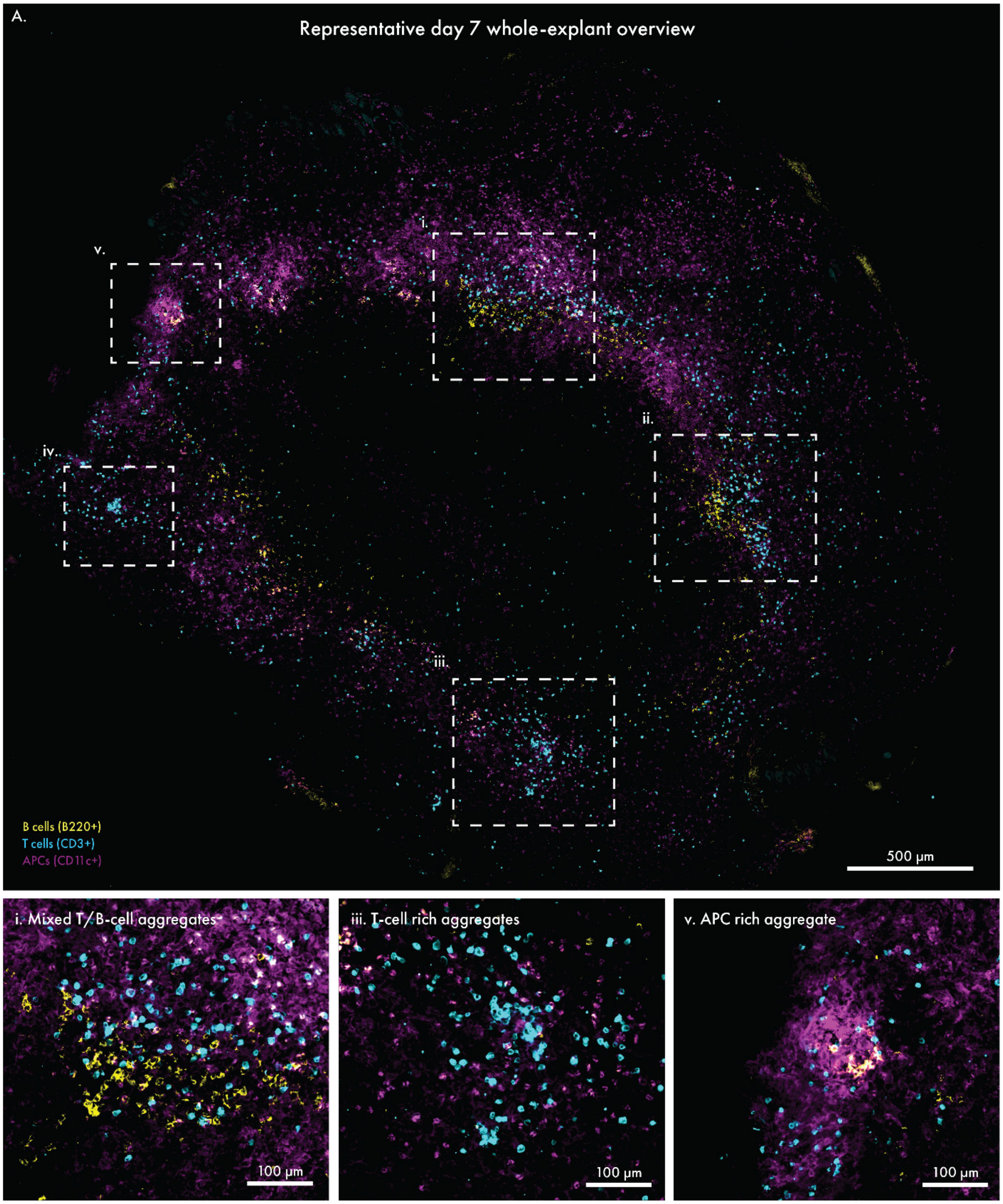

**Supplementary Figure 8. Spatial heterogeneity of early lymphoid organization within a day-7 soft hydrogel.** The same representative day-7 soft-hydrogel explant shown in Supplementary Figure 7A is displayed as a whole-explant overview, with boxed regions highlighting additional patterns of local immune organization. Central plane sections were stained for B cells (B220$^+$, yellow), T cells (CD3$^+$, cyan), and CD11c$^+$ antigen-presenting cells (magenta). (**A**) Representative whole-explant image showing the broad distribution of infiltrating immune cells. Boxed regions (i–v) identify multiple examples of local immune aggregates, including mixed T/B-cell aggregates (i–ii), T-cell-rich aggregates (iii–iv), and an APC-rich aggregate (v). Representative examples of these patterns (i, iii, and v) are enlarged below. Scale bars, 500 µm in (A) and 100 µm in the enlarged images.

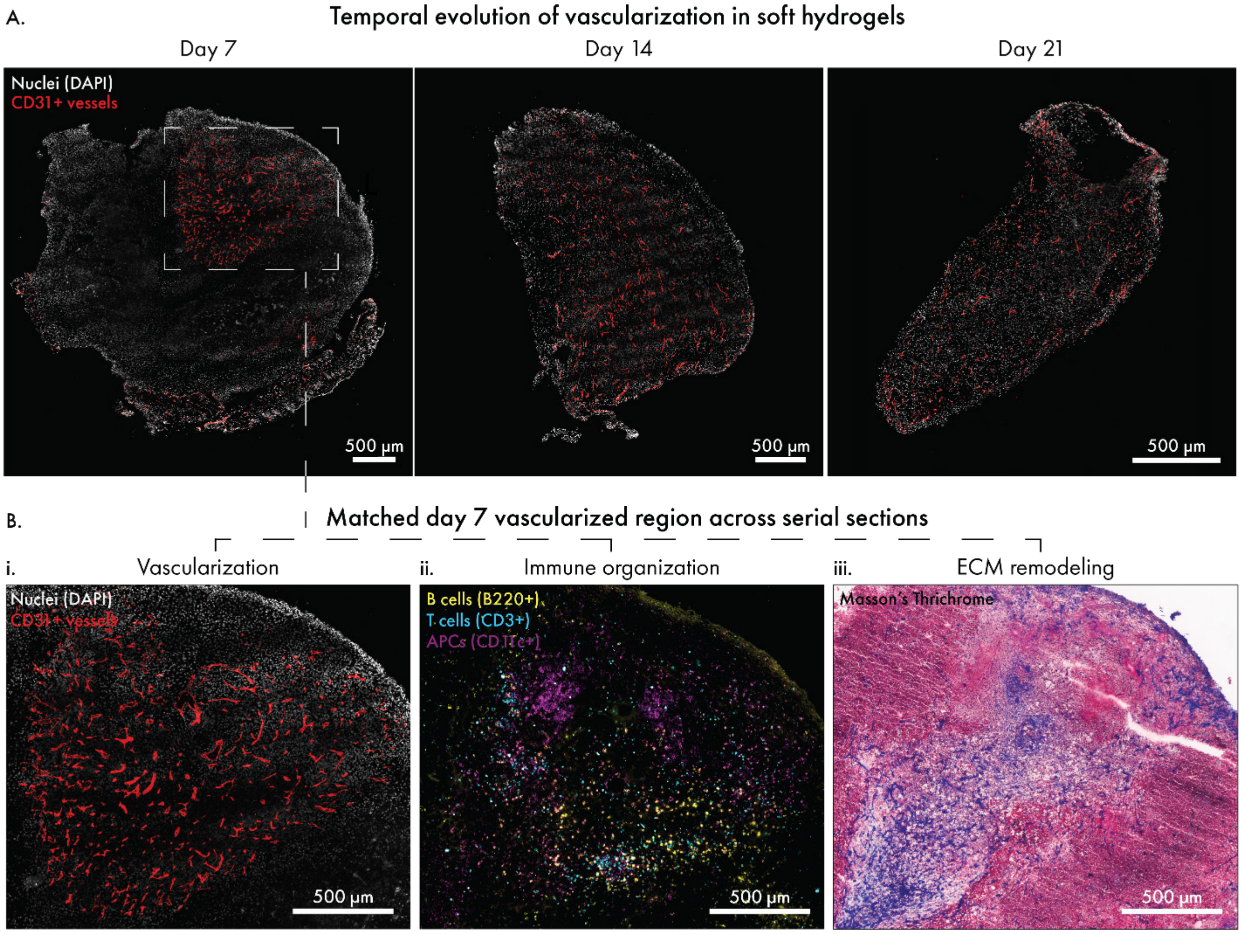

**Supplementary Figure 9. Temporal vascularization of soft hydrogels and spatial association with immune organization and ECM remodeling.** Soft hydrogels containing ovalbumin (OVA), CCL21, and LIGHT were implanted subcutaneously and retrieved at days 7, 14, and 21. Explants were sectioned through their central plane for spatial characterization. (**A**) Representative whole-explant sections collected at days 7, 14, and 21, stained for nuclei (DAPI, gray) and blood vessels ($CD31^+$, red), illustrating the temporal distribution of vasculature across soft-hydrogel explants. The dashed box in the day-7 explant indicates the vascularized region examined across matched serial sections in (B). (**B**) Matched day-7 vascularized region examined across serial sections. (i) CD31 and DAPI staining showing local vascularization. (ii) Corresponding immunofluorescence section stained for B cells ($B220^+$, yellow), T cells ($CD3^+$, cyan), and $CD11c^+$ antigen-presenting cells (magenta), showing spatial association of the vascularized region with local immune-cell infiltration and organization. (iii) Corresponding Masson's Trichrome-stained section showing collagen-rich extracellular matrix deposition within the same region. Scale bars, 500 μm.

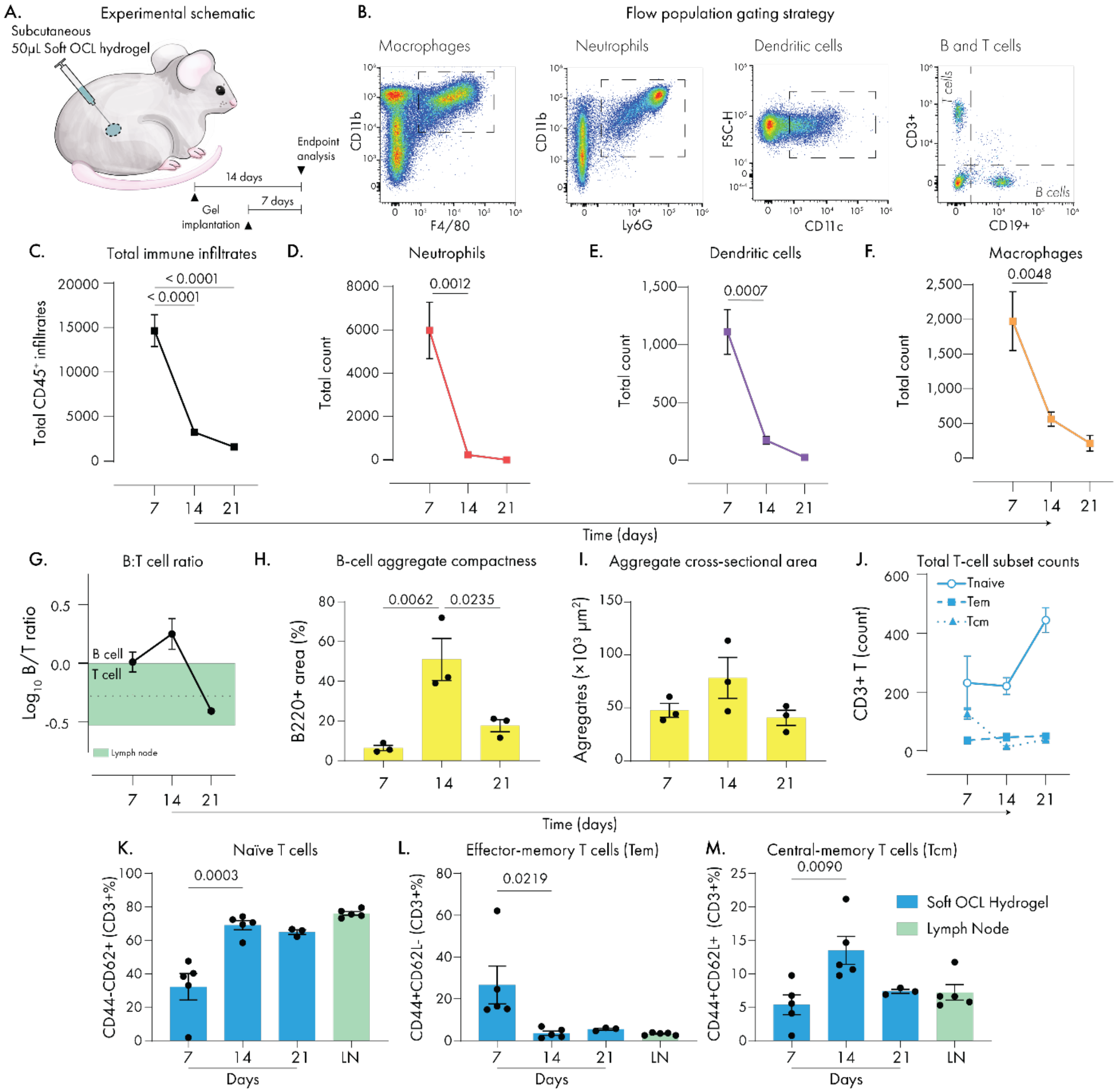

**Supplementary Figure 10. Temporal flow-cytometric and morphometric characterization of TLS-like niche development in soft OCL hydrogels.** (**A**) Experimental schematic illustrating collection of soft OCL hydrogels containing ovalbumin, CCL21, and LIGHT at days 7 and 14 after implantation for temporal comparison with day-21 explants collected in the preceding study. (**B**) Representative flow-cytometry gating strategy used to identify immune-cell populations within live CD45$^+$ cells. Macrophages were identified as CD11b$^+$F4/80$^+$ cells. Within the remaining F4/80$^-$ population, neutrophils were identified as CD11b$^+$Ly6G$^+$ cells. Following exclusion of macrophages and neutrophils, CD11b$^-$ cells were used to identify CD3$^+$CD19$^-$ T cells and CD3$^-$CD19$^+$ B cells. Dendritic cells were identified as CD11c$^+$ cells after exclusion of macrophages, neutrophils, T cells, and B cells. Representative plots are shown for a day-7 soft OCL hydrogel. (**C**) Total CD45$^+$ leukocyte recovery from soft OCL hydrogel explants over time. (**D**–**F**) Total counts of neutrophils (D), dendritic cells (E), and macrophages (F) recovered at days 7, 14, and 21. (**G**) Log$_2$-transformed B-cell-to-T-cell ratio over time. Values above zero indicate B-cell predominance, whereas values below zero indicate T-cell predominance. The green band indicates the mean ± SEM calculated from lymph node samples using the same population definitions and transformation (n = 5 biologically independent mice). (**H**) B-cell aggregate compactness, calculated as the percentage of the manually outlined aggregate area occupied by thresholded B220$^+$ signal. (**I**) Cross-sectional area of manually outlined cellular aggregates at each timepoint. (**J**) Absolute numbers of naïve, effector-memory (Tem), and central-memory (Tcm) T cells recovered from soft OCL hydrogel explants over time. (**K**–**M**) Frequencies of naïve T cells (K), Tem cells (L), and Tcm cells (M) among CD3$^+$ T cells in soft OCL hydrogels at days 7, 14, and 21 and in lymph nodes. Naïve, Tem, and Tcm cells were defined as CD44$^-$CD62L$^+$, CD44$^+$CD62L$^-$, and CD44$^+$CD62L$^+$, respectively. Data are presented as mean ± SEM. For flow-cytometry analyses, n = 5 biologically independent mice at days 7 and 14 and n = 3 at day 21. Lymph node data represent n = 5 biologically independent mice. For aggregate analyses, all identifiable B-cell-rich aggregates within each analyzed hydrogel section were quantified and averaged to obtain a single value per hydrogel. Each point represents one biologically independent hydrogel (n = 3 per timepoint). For (C–F), (H–I), and (K–M), statistical significance was assessed using one-way ANOVA followed by Tukey's multiple-comparisons test. Exact *p* values are shown where indicated.

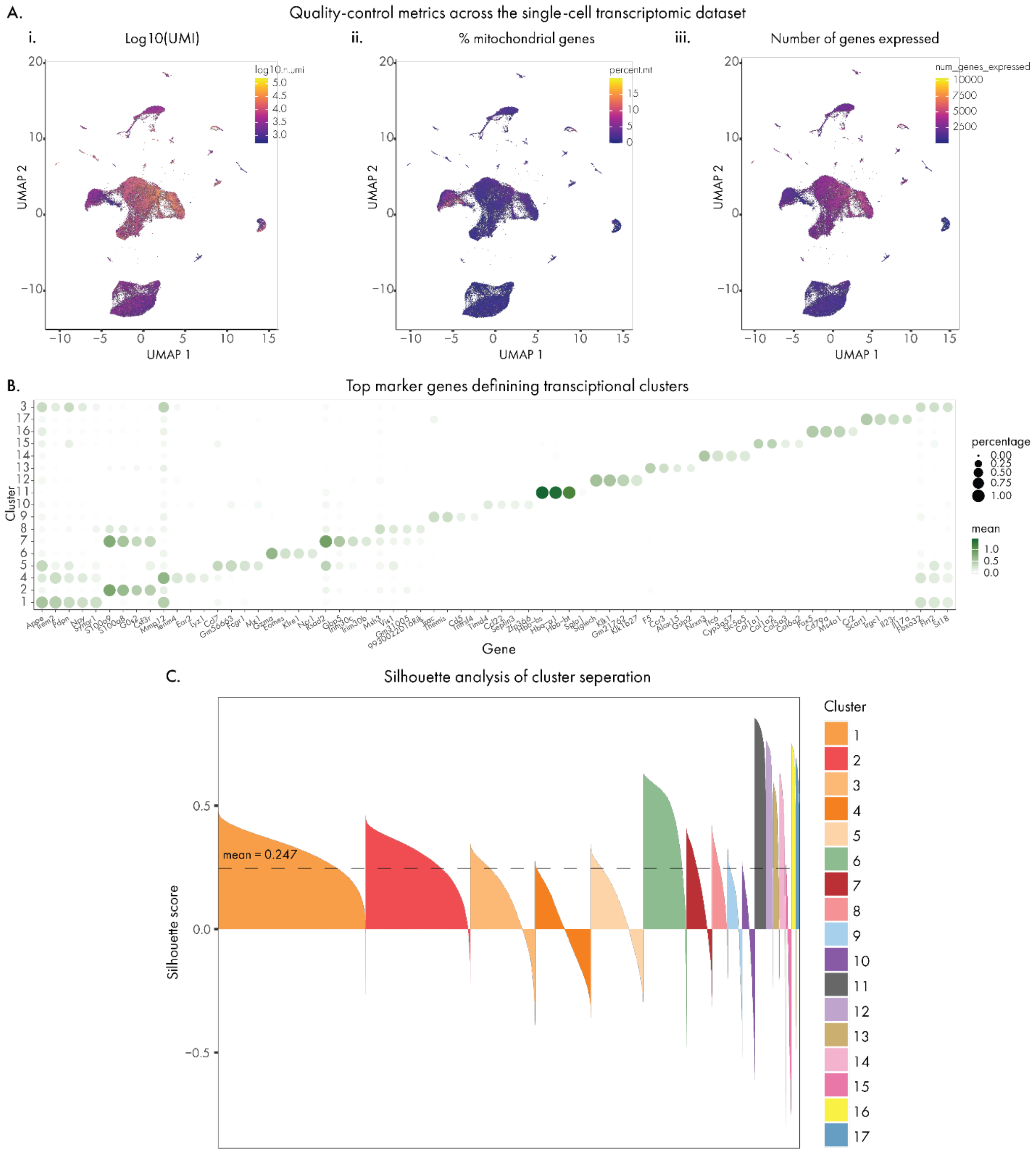

**Supplementary Figure 11. Quality control and cluster annotation of the single-cell RNA-sequencing dataset.** (**A**) Uniform Manifold Approximation and Projection (UMAP) visualizations of the 69,350 cells retained after quality control, colored by $log_{10}$-transformed unique molecular identifier (UMI) count, percentage of mitochondrial transcripts, and number of detected genes. (**B**) Dot plot showing top-ranked cluster-specific genes identified using the top_markers() function in monocle3. Dot size indicates the percentage of cells expressing each gene, and color indicates mean normalized expression. (**C**) Silhouette scores of the 17 transcriptional clusters. Individual cells are grouped by cluster and ordered by silhouette score; positive values indicate greater similarity to cells within the assigned cluster than to the nearest alternative cluster, whereas negative values indicate greater overlap with neighboring clusters. The dashed red line indicates the mean silhouette score across all cells (0.247). Colors denote cluster identity.

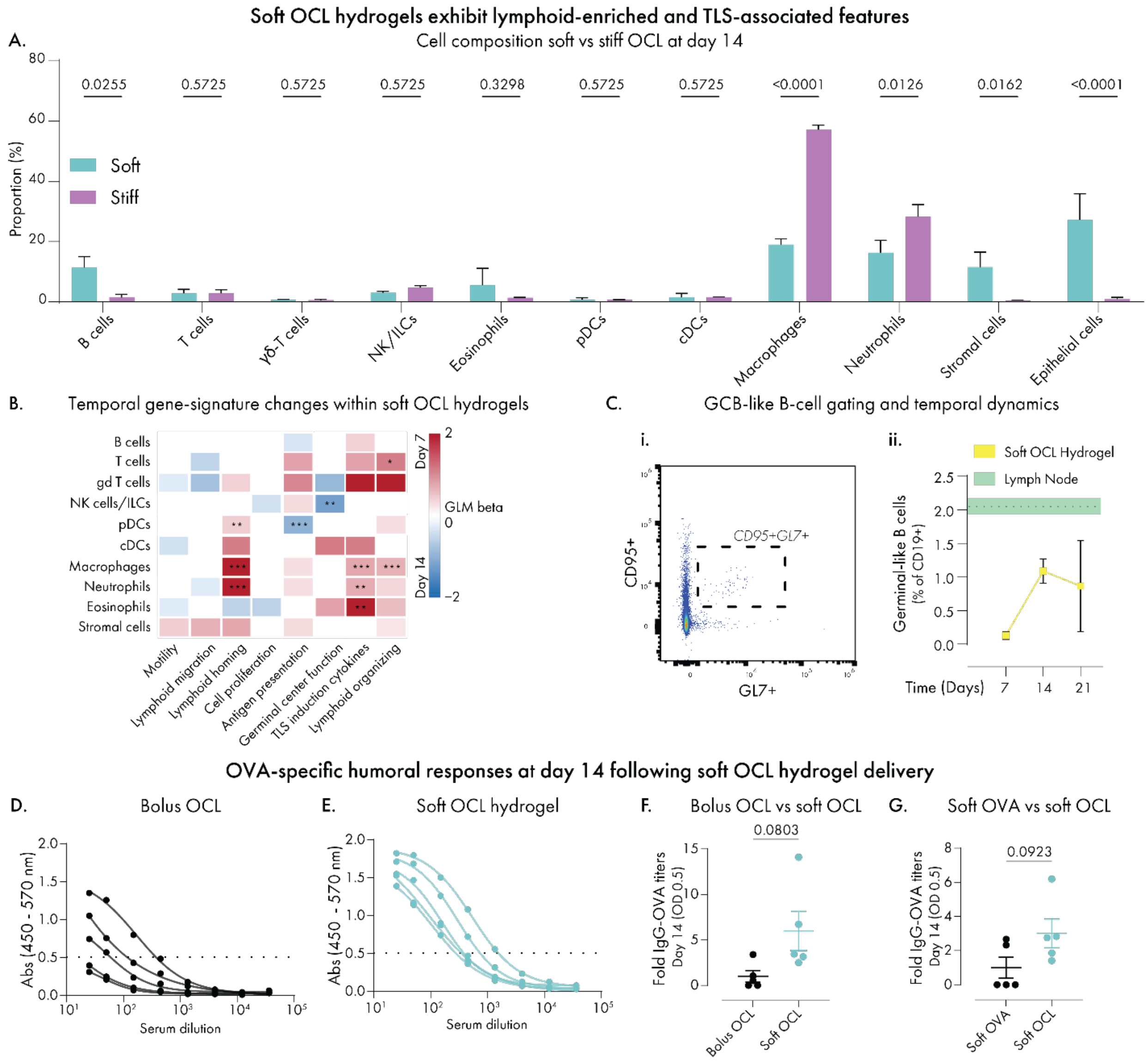

**Supplementary Figure 12. Cellular composition, temporal TLS-associated transcriptional programs, and adaptive immune responses in soft OCL hydrogels.** (**A**) Cell-type proportions in soft and stiff OCL hydrogels at day 14 following single-cell RNA sequencing. (**B**) Cell type-specific differences in aggregate expression scores for TLS-associated transcriptional signatures within soft OCL hydrogels at day 7 relative to day 14. Positive model coefficients indicate higher signature scores at day 7, whereas negative coefficients indicate higher scores at day 14; coefficients with an absolute $\beta \leq 0.25$ were set to zero. (**C**) Germinal-center-like B-cell identification and temporal dynamics. (**i**) Representative flow-cytometry gating strategy used to identify $CD95^+GL7^+$ germinal-center-like B cells among $CD19^+$ B cells in a lymph node sample. (**ii**) Frequencies of germinal-center-like B cells among $CD19^+$ B cells in soft OCL hydrogels at days 7, 14, and 21, with lymph nodes shown as a reference. (**D–F**) OVA-specific total IgG responses measured by ELISA at day 14 in an experiment comparing bolus OCL cargo with soft OCL hydrogels. (**D–E**) Serum dilution curves following administration of bolus OCL cargo (**D**) or soft OCL hydrogels (**E**). Each curve represents one biologically independent mouse. Absorbance was measured at 450 nm with background correction at 570 nm. (**F**) Day-14 OVA-specific IgG titers from the same experiment. Titers were defined as the serum dilution corresponding to an optical density of 0.5 and normalized to the bolus OCL reference group. (**G**) Day-14 OVA-specific IgG titers from a separate experiment comparing soft hydrogels containing OVA alone or OCL cargo. Titers were normalized to the OVA-only hydrogel reference group. Normalized values in (F–G) are expressed relative to the respective reference group within each independent experiment. Data in (A, Cii, F–G) are presented as mean ± SEM. For single-cell RNA-sequencing analyses in (A–B), n = 3 independently processed biological pools per condition. Statistical significance in (A) was assessed using two-way ANOVA followed by the two-stage Benjamini, Krieger and Yekutieli multiple-comparisons procedure, with soft and stiff hydrogels compared within each cell type. FDR-adjusted *q* values are shown. For (B), aggregate signature scores were compared between days 7 and 14 within each cell type using Gaussian regression with timepoint and experiment as model terms, followed by Benjamini–Hochberg correction. Coefficients with an absolute $\beta > 0.25$ are shown regardless of adjusted *P* value, with asterisks indicating statistical significance (*q < 0.1, **q < 0.01, ***q < 0.001). For (Cii), n = 5 biologically independent mice at days 7 and 14, n = 3 at day 21, and n = 5 lymph node samples. For (F–G), n = 5 biologically independent mice per group. Statistical significance in (F–G) was assessed using unpaired two-tailed Student's *t*-tests. Exact *p* values are shown for these comparisons.

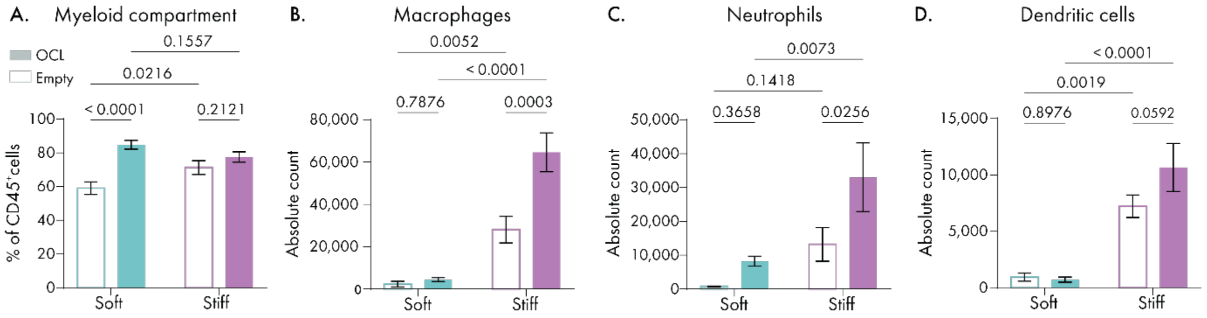

**Supplementary Figure 13. Day-7 myeloid composition and accumulation in empty and OCL-loaded hydrogels.** (**A**) Combined macrophage, neutrophil, and dendritic-cell compartment as a percentage of CD45$^+$ cells in empty and OCL-loaded soft and stiff hydrogels at day 7. (**B**–**D**) Total numbers of macrophages (**B**), neutrophils (**C**), and dendritic cells (**D**) recovered from empty and OCL-loaded soft and stiff hydrogels at day 7. Data are presented as mean ± SEM; n = 5 analyzed hydrogel explants per condition from three mice. Statistical significance was assessed using one-way ANOVA followed by Tukey's multiple-comparisons test. Exact *p* values are shown where indicated.

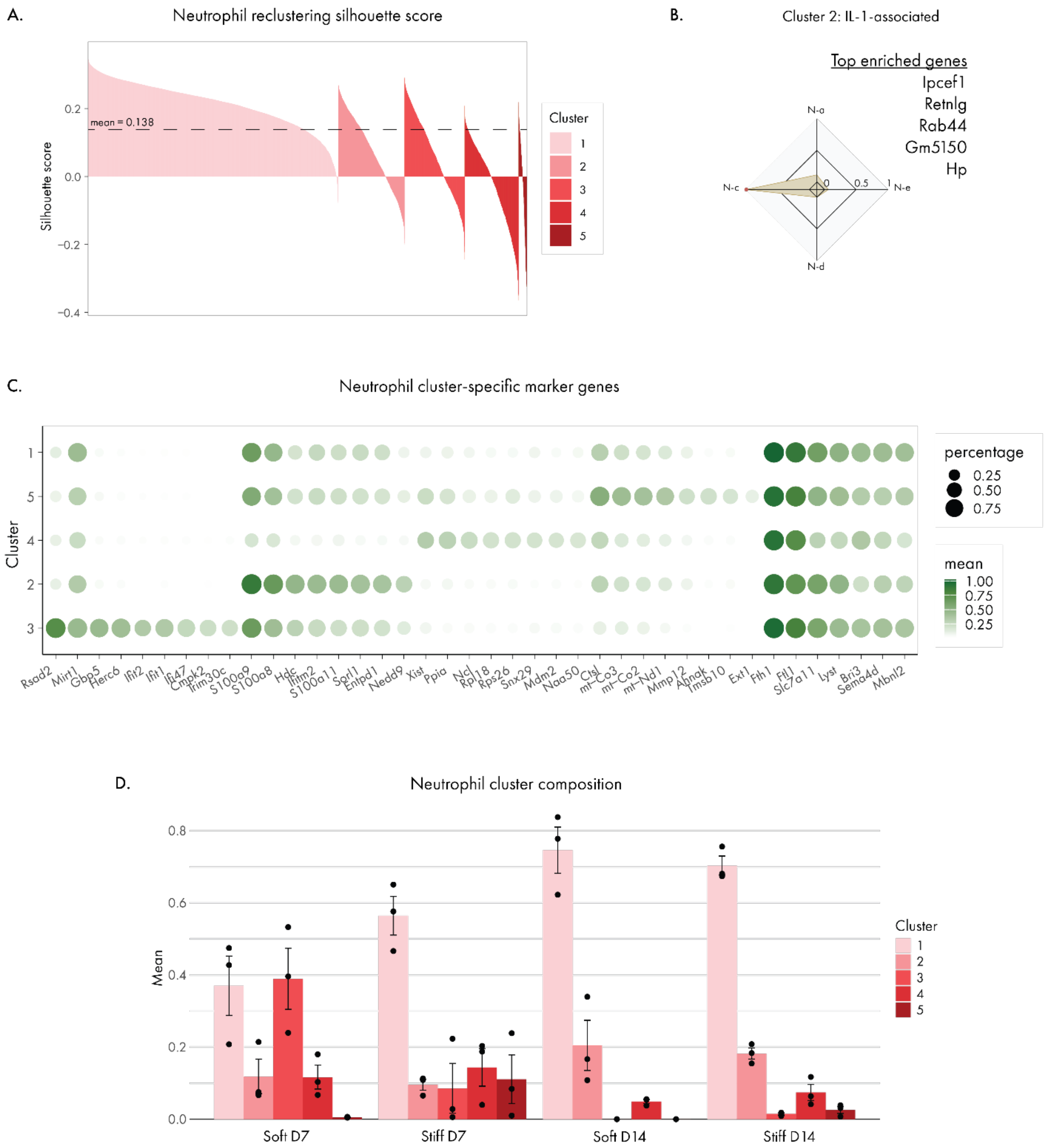

**Supplementary Figure 14. Transcriptional characterization and temporal distribution of neutrophil states in soft and stiff OCL hydrogels.** (**A**) Silhouette scores for the neutrophil reclustering solution, resolving five transcriptional clusters. Colors indicate Clusters 1–5, and the dashed line indicates the mean silhouette score of 0.138. (**B**) Representative Immune Response Enrichment Analysis of neutrophil Cluster 2, identifying an IL-1-associated transcriptional state together with representative cluster-associated genes. (**C**) Top-ranked cluster-specific marker genes identified following neutrophil reclustering. Dot size indicates the percentage of cells expressing each gene, and color indicates mean normalized expression. (**D**) Relative representation of neutrophil Clusters 1–5 in soft and stiff OCL hydrogels at days 7 and 14. Bars show mean ± SEM across n = 3 independently processed biological pools per condition; individual pools are shown as points.

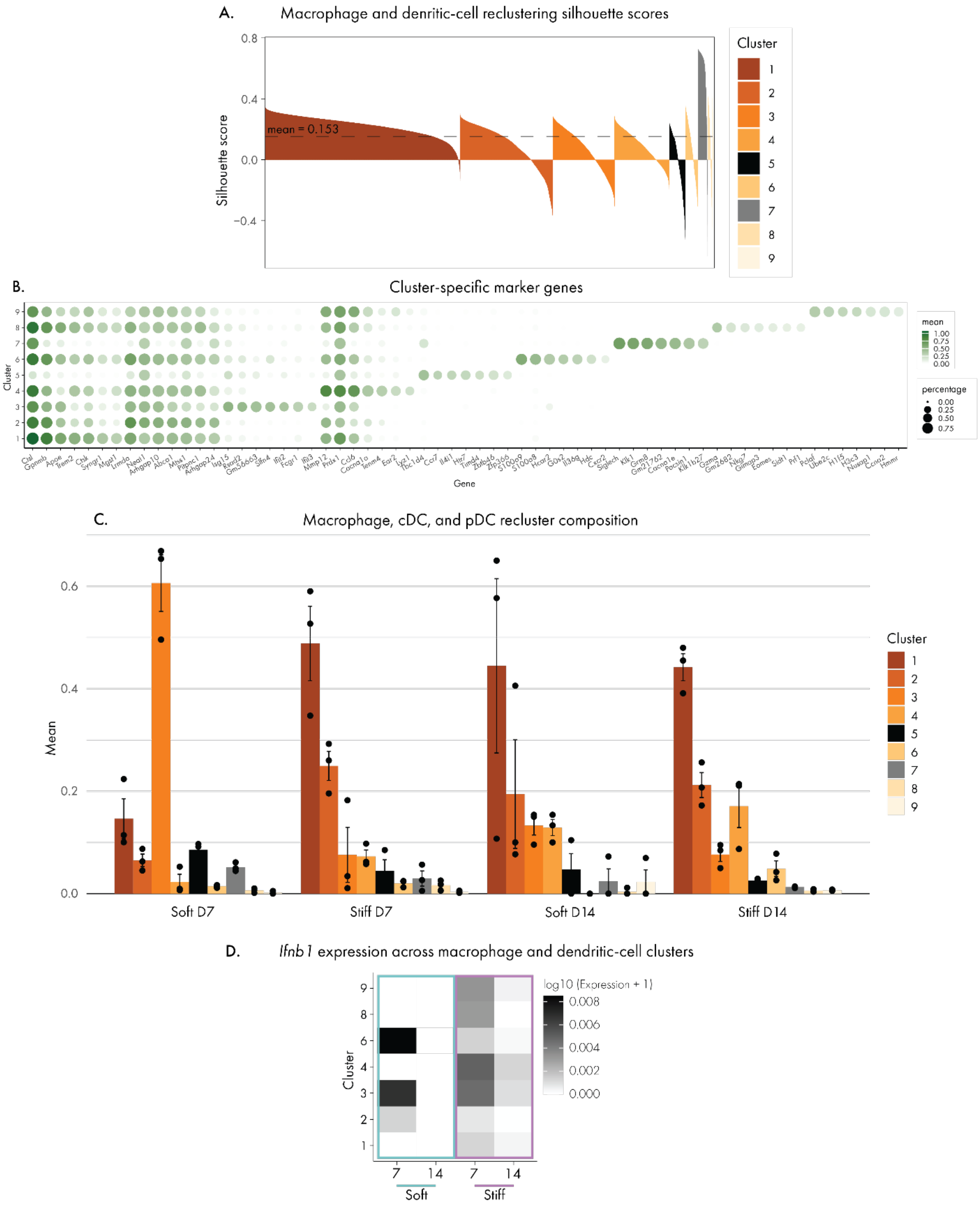

**Supplementary Figure 15. Reclustering and transcriptional annotation of macrophage, dendritic-cell, and pDC populations.** (**A**) Silhouette scores for the joint reclustering of macrophages, cDCs, and pDCs. Colors indicate Clusters 1-9, and the dashed line indicates the mean silhouette score of 0.153. (**B**) Top-ranked cluster-specific marker genes identified following reclustering. Dot size indicates the percentage of cells expressing each gene, and color indicates mean normalized expression. (**C**) Relative representation of the reclustered populations in soft and stiff OCL hydrogels at days 7 and 14. Clusters 1–4, 6, 8, and 9 represent macrophage populations, Cluster 5 represents cDCs, and Cluster 7 represents pDCs. Bars show mean ± SEM across n = 3 independently processed biological pools per condition; individual pools are shown as points. (**D**) Mean normalized *Ifnb1* expression across macrophage, cDC, and pDC clusters in soft and stiff OCL hydrogels at days 7 and 14. Heatmap color represents $\log_{10}$ (expression + 1).

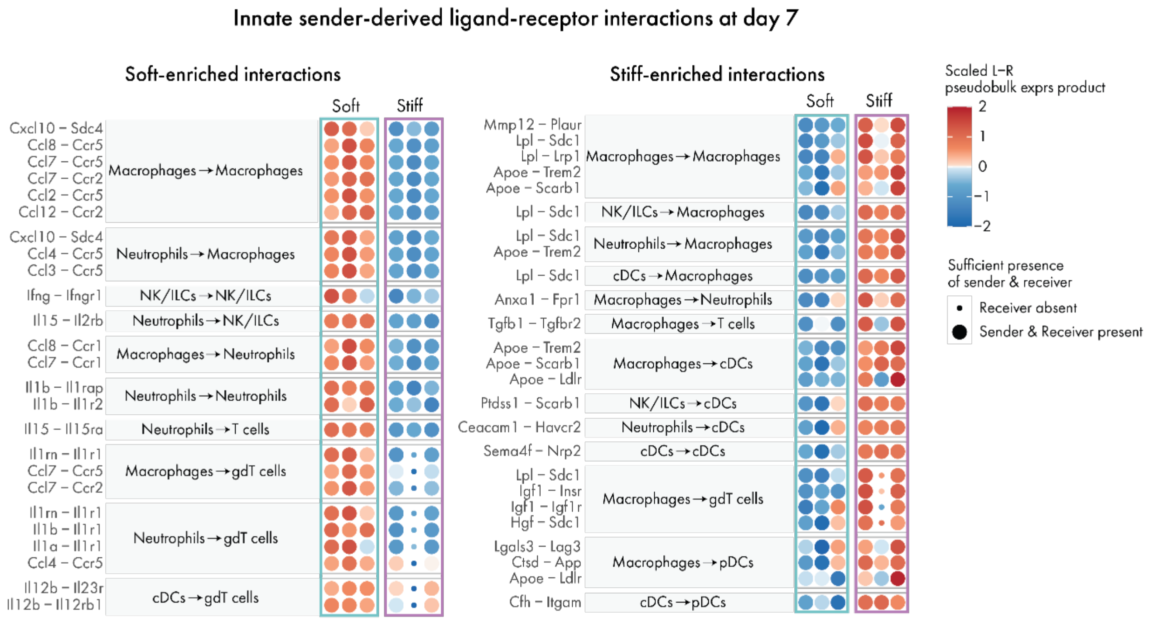

**Supplementary Figure 16. Differential ligand–receptor interactions from selected innate sender populations at day 7.** MultiNicheNet analysis of predicted cell–cell communication originating from selected innate sender populations in soft and stiff OCL hydrogels at day 7. The top 25 prioritized ligand–receptor interactions favoring soft hydrogels (left) and stiff hydrogels (right) are shown, grouped by sender-to-receiver cell pair. Selected innate sender populations included macrophages, neutrophils, cDCs, and NK cells/ILCs, with interactions evaluated across eligible immune-cell receiver populations. Each row indicates a ligand–receptor pair, and the three points shown for each condition represent n = 3 independently processed biological pools. Color indicates the scaled ligand–receptor pseudobulk expression product, with positive and negative values reflecting relatively higher and lower interaction-associated expression, respectively. Dot size indicates sufficient representation of the corresponding sender and receiver populations within each pool; small dots indicate absence of the receiver population, whereas large dots indicate that both sender and receiver populations were present. Selected interactions from this prioritized set are shown in Fig. 6N.

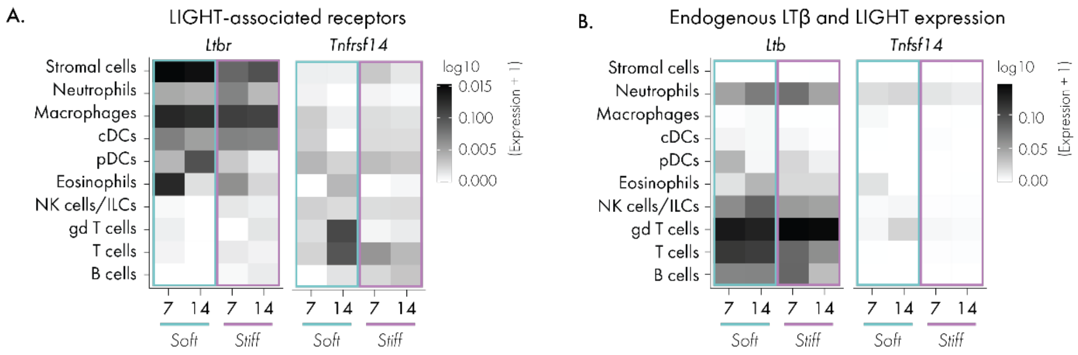

**Supplementary Figure 17. LIGHT receptor and endogenous LTβR-associated ligand expression across soft and stiff OCL hydrogels.** (**A**) Mean normalized *Ltbr* and *Tnfrsf14* expression across stromal and immune cell populations in soft and stiff OCL hydrogels at days 7 and 14. (**B**) Mean normalized *Ltb* and *Tnfsf14* expression across stromal and immune cell populations in soft and stiff OCL hydrogels at days 7 and 14. Heatmap color represents $\log_{10}$(expression + 1).

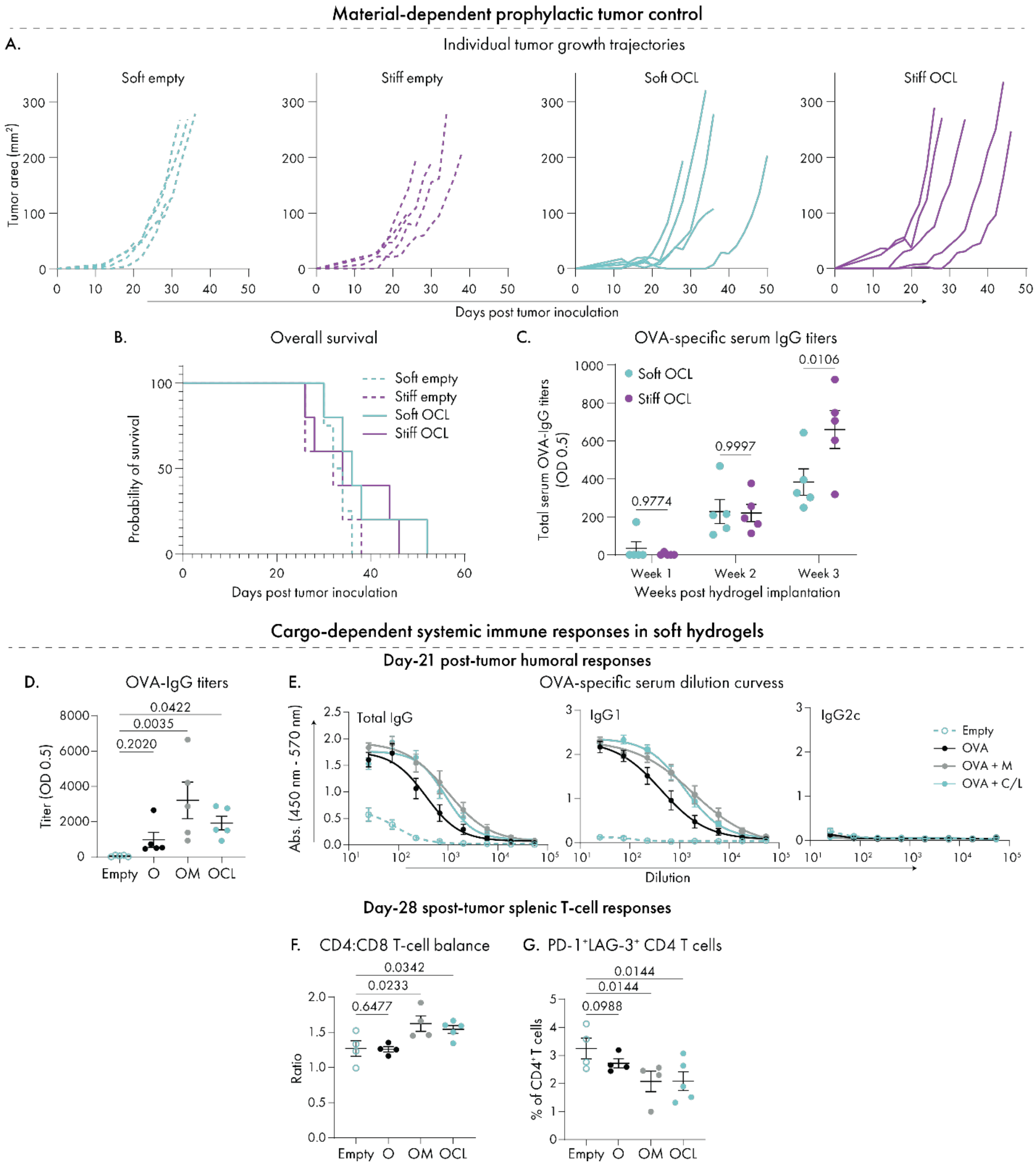

**Supplementary Figure 18. Individual tumor trajectories, survival and systemic immune responses following prophylactic hydrogel implantation.** (**A**) Individual B16F10-OVA tumor-growth trajectories in mice receiving soft empty, stiff empty, soft OCL or stiff OCL hydrogels 14 days before tumor inoculation, followed by twice-weekly anti-PD-L1 treatment beginning at day 14 after tumor challenge. (**B**) Kaplan–Meier survival curves for the corresponding treatment groups. (**C**) OVA-specific serum total IgG titers measured weekly following implantation of soft or stiff OCL hydrogels in a separate immunization study in naïve mice. (**D**) OVA-specific serum total IgG titers at day 21 after tumor inoculation in mice receiving empty, OVA-only (O), OVA plus MPLA (OM) or OCL soft hydrogels. (**E**) OVA-specific total IgG, IgG1 and IgG2c serum dilution curves at day 21 after tumor inoculation. (**F**) Splenic $CD4^+/CD8^+$ T-cell ratios and (**G**) frequencies of PD-$1^+$LAG-$3^+$ cells among splenic $CD4^+$ T cells at day 28 after tumor inoculation. Data are presented as mean ± SEM. In (A–B), n = 4 biologically independent mice for the soft empty group and n = 5 in each of the remaining groups. Survival curves in (B) were compared using the log-rank (Mantel–Cox) test. In (C–E), n = 5 biologically independent mice per group. For the day-28 splenic analyses in (F–G), n = 4 biologically independent mice in the empty, O, and OM groups and n = 5 in the OCL group. Statistical significance in (C) was assessed using two-way ANOVA followed by Šídák's multiple-comparisons test. Statistical significance in (D–G) was assessed using one-way ANOVA followed by the two-stage Benjamini, Krieger and Yekutieli multiple-comparisons procedure, with comparisons made against the empty-hydrogel group. Exact *p* or FDR-adjusted *q* values are shown where indicated.

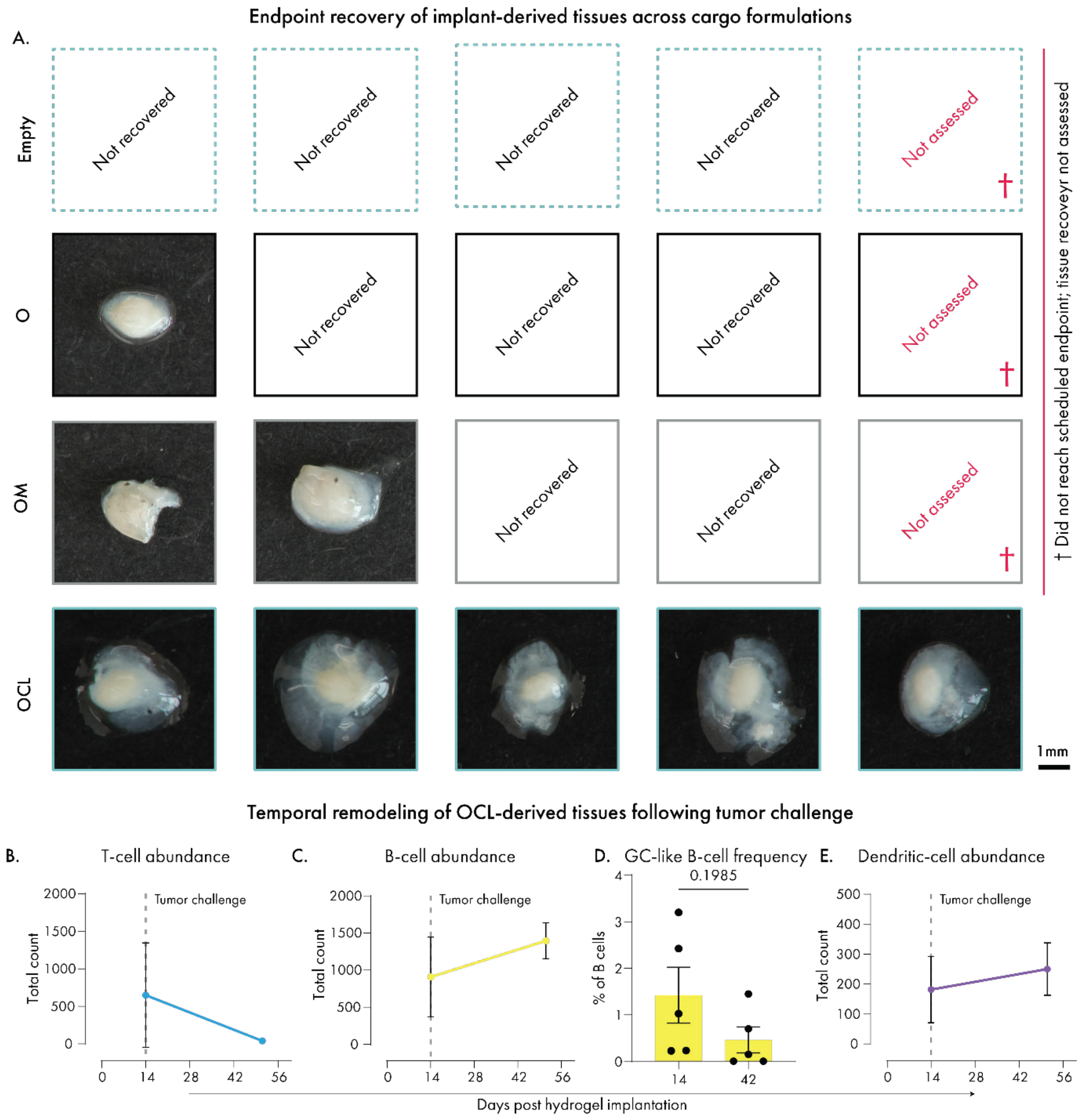

**Supplementary Figure 19. Recovery and temporal immune remodeling of soft OCL-derived tissues following tumor challenge.** (**A**) Recovery of implant-derived tissues at experimental endpoint, corresponding to day 42 after hydrogel implantation and 28 days after tumor challenge. Macroscopic images are shown for all identifiable tissues recovered from mice receiving empty, OVA-only (O), OVA plus MPLA (OM) or OCL soft hydrogels. Boxes labeled "Not recovered" indicate implantation sites from which no identifiable tissue was recovered. One mouse in each of the empty, O and OM groups did not reach the scheduled endpoint; these implantation sites were therefore not assessed for tissue recovery, as indicated by "Not assessed" boxes. Among implantation sites assessed at endpoint, tissues were recovered from 0/4 empty, 1/4 O, 2/4 OM and 5/5 OCL implantation sites. Scale bar, 1 mm; applies to all macroscopic tissue images. (**B**–**E**) Flow-cytometric analysis of soft OCL-derived tissues collected from independent cohorts at day 14 after implantation, before tumor challenge, and at day 42 after implantation, corresponding to 28 days after tumor challenge. Total numbers of $CD3^+$ T cells (B), $CD19^+$ B cells (C), GC-like $CD95^+GL7^+$ B cells as a percentage of total B cells (D), and $CD11c^+$ dendritic cells (E) are shown. Dashed lines in (B, C and E) indicate the timing of tumor challenge. Data are presented as mean ± SEM; n = 5 biologically independent mice per timepoint. Statistical significance in (D) was assessed using an unpaired two-tailed Student's t-test. Exact *p* values are shown where indicated.

**Supplementary Table 1. Frequency dependence of the storage modulus in soft and stiff hydrogels.** The frequency-scaling exponent x was obtained by fitting $G' \sim \omega^x$ to frequency-sweep data. Values are reported as the fitted exponent ± standard error with 95% confidence intervals.

| Hydrogel formulation | Frequency dependence ($G' \sim \omega^x$) | 95% confidence interval |
|---|---|---|
| Soft | 0.4355 ± 0.003806 | (0.4277, 0.4433) |
| Stiff | 0.2287 ± 0.005066 | (0.2183, 0.2390) |

**Supplementary Table 2. Predicted charge classification and estimated hydrodynamic diameter of protein cargos.** Theoretical isoelectric points were determined using the indicated protein sequence ranges. Charge at physiological pH was classified from the difference between theoretical pI and pH 7.4. Proteins were classified as positively charged when pI − 7.4 ≥ 1, negatively charged when pI − 7.4 ≤ −1, and approximately neutral when −1 < pI − 7.4 < 1. Effective hydrodynamic diameters were estimated from molecular weight using the empirical mass–radius scaling relationship for globular proteins implemented in the FIDA Bio molecular weight-to-size calculator. These estimates assume a compact, globular protein geometry and do not represent experimentally measured structural dimensions.

| **Protein** | **Accession** | **Sequence range** | **Theoretical pI** | **Predicted charge at pH 7.4** | **Estimated hydrodynamic diameter** |
|---|---|---|---|---|---|
| Chicken ovalbumin (OVA) | UniProt P01012 | Ser2–Pro386 | 5.19 | Negative | 6.3 nm |
| Mouse CCL21 (CCL21a) | UniProt P84444 | Ser24–Gly133 | 10.00 | Positive | 3.82 nm |
| Mouse LIGHT | UniProt Q9QYH9 | Asp72–Val239 | 9.55 | Positive | 4.68 nm |

**Supplementary Table 3. Single-cell RNA-sequencing pooling and library-loading design.**

| Condition | Biological pool | Experiment | Hydrogels per pool | Lanes loaded | Target cells per lane |
|---|---|---|---|---|---|
| Soft, day 7 | 1 | 2 | 4 | 4 | 5,000 |
| Soft, day 7 | 2 | 3 | 2 | 2 | 5,000 |
| Soft, day 7 | 3 | 3 | 2 | 2 | 5,000 |
| Stiff, day 7 | 1 | 2 | 3 | 4 | 5,000 |
| Stiff, day 7 | 2 | 3 | 1 | 2 | 5,000 |
| Stiff, day 7 | 3 | 3 | 2 | 2 | 5,000 |
| Soft, day 14 | 1 | 1 | 2 | 2 | 20,000 |
| Soft, day 14 | 2 | 3 | 2 | 2 | 5,000 |
| Soft, day 14 | 3 | 3 | 2 | 2 | 5,000 |
| Stiff, day 14 | 1 | 1 | 2 | 2 | 20,000 |
| Stiff, day 14 | 2 | 3 | 1 | 2 | 5,000 |
| Stiff, day 14 | 3 | 3 | 2 | 2 | 5,000 |

**Note:** Each row represents one independently processed biological pool. Three pooled replicates were analyzed per condition, with each pool comprising 1–4 hydrogel explants. The number of hydrogels and lanes varied across experiments to achieve sufficient cell concentrations for library preparation.

**Supplementary Table 4. Gene-signature definitions used for aggregate expression scoring.** Gene sets encompassed cell motility, lymphoid organization, TLS-induction-associated cytokines, lymphoid migration, lymphoid homing, cell proliferation, germinal-center function, and antigen presentation. The genes included in each signature are listed below.

| Signature | Number of unique genes | Gene set |
|---|---|---|
| Cell motility | 28 | *Rac2, Dock2, Dock8, Coro1a, Was, Wipf1, Arpc1b, Arpc2, Arpc3, Itgb2, Itgal, Itgam, Itga4, Ccr7, Cxcr4, Cxcr3, Ccr2, Ccr5, Cx3cr1, S1pr1, Cd44, Sell, Msn, Ezr, Talin1, Vcl, Ptk2, Pxn* |
| Lymphoid organization | 16 | *Ltb, Ltbr, Tnfsf14, Relb, Nfkb2, Traf3, Traf6, Icam1, Vcam1, Ccl19, Ccl21a, Cxcl13, Cxcr5, Sell, Lamp3, Cxcl12* |
| TLS-induction-associated cytokines | 14 | *Il4, Il6, Il7, Il13, Il17a, Il21, Il22, Il23, Il1b, Il36g, Tnf, Tnfsf14, Tnfsf13, Ifnb1* |
| Lymphoid migration | 16 | *Ccr7, Sell, S1pr1, Klf2, Cxcr4, Cxcr5, S1pr2, Itgal, Itgb2, Itga4, Itgb1, Dock2, Gnai2, Rac1, Pik3cd, Pik3cg* |
| Lymphoid homing | 20 | *Ccr7, Ccl19, Cxcl13, Cxcl11, Cxcl10, Cd83, Lamp3, Ido1, Ido2, Enpp2, Nr4a3, Chst7, Chst3, Adam19, Map3k13, Runx2, Plcl1, Spats2l, Ifit2, Rsad2* |
| Cell proliferation | 97 | *unique(unlist(cc.genes.mouse))* |
| Germinal-center function | 12 | *Cd19, Ms4a1, Cd79a, Cd79b, Bcl6, Aicda, Rgs13, Fas, Pax5, Ung, Icosl, Cxcr4* |
| Antigen presentation | 13 | *H2-Aa, H2-Ab1, H2-Eb1, Cd74, H2-DMa, H2-DMb1, Ciita, Cd80, Cd86, Cd40, Icosl, Icam1, Tnfsf9* |

**Note:** The cell proliferation signature was supplied in the source workbook as the R expression unique(unlist(cc.genes.mouse)); the expanded gene-level contents were not included in the workbook.
